# Bifunctional Phagocytic Synapse Enhancers for Cancer Immunotherapy

**DOI:** 10.1101/2025.03.13.642658

**Authors:** Valerio Sabatino, Silvan Fleisch, Cong Tang, Felix M. Müller, Carlos Labão-Almeida, Wei Ting Khaw, Sabrina Hogan, Derrick R. Hicks, Marcia Fontes, Ana R. Coelho, Wei Yang, Irene Sarkar, Debarati Shome, Aldrin Vasco Vidal, Mar Cabeza-Cabrerizo, Theresa Rohm, Rogier M. Reijmers, Deniz Kaymak, Fiona Gerster, David Baker, Rita Fior, Gregor Hutter, Gonçalo J. L. Bernardes

## Abstract

Immunotherapy profoundly impacted cancer treatments by harnessing the patient’s immune system. Phagocytosis, the process whereby immune cells engulf and destroy foreign particles or cells, plays a critical role in tumour cell clearance. Herein, we introduce a novel concept termed “ENPHASYS” – <u>En</u>hancement of <u>Pha</u>gocytic <u>Sy</u>napse<u>s</u> – designed to direct and amplify phagocytosis of cancer cells using heterobifunctional molecules named phagocytic synapse enhancers (PSEs). By engineering a de novo PD-L1 binder linked to a natural phagocytosis promoting peptide, tuftsin, the resulting PSE combines PD-L1 blockade with enhanced tumour cell phagocytosis; in addition, the PSEs induce macrophages to internalize a membrane or extracellular target. Intratumoural treatment of colorectal carcinoma- or glioblastoma-burdened immunocompetent animals resulted in beneficial overall survival, delayed tumour growth and a potent antitumor response driven by T-cell activation and TAM reprogramming, underpinning the translational relevance of ENPHASYS.

**Graphical Abstract:** 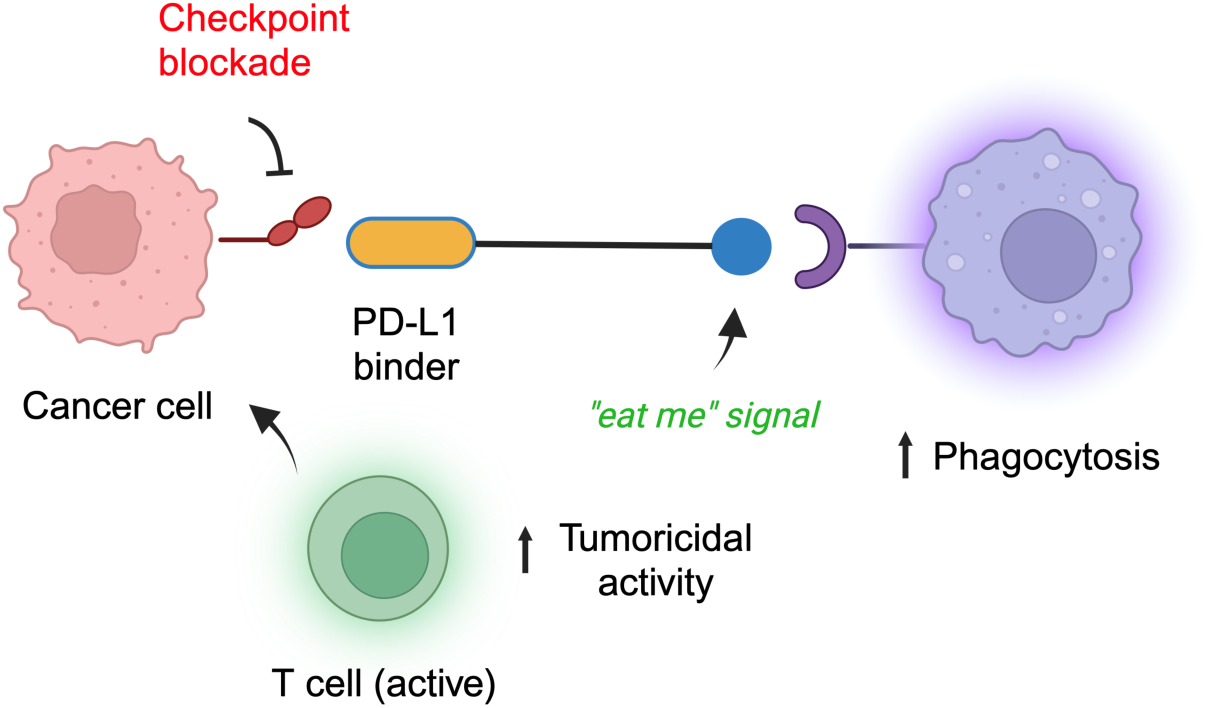

## Introduction

Immunotherapy has revolutionised the standard of care for cancer patients. Among the various hallmarks of cancer, the discovery of immune checkpoints profoundly impacted cancer therapy^1–4^. Nevertheless, only about 20% of cancer treatments rely on immunotherapy, and its effectiveness is hampered by adverse events and/or mechanisms of resistance^5^. The tumour microenvironment (TME) contains a significant amount of myeloid cells that support cancer development. Macrophages (Mϕ) are specialised in clearing pathogens and cancerous cells^6^. However, in the immunosuppressive TME, Mϕ undergo phenotypic changes to differentiate into tumour associated Mϕ (TAMs), which are often associated with poor prognosis. Immunotherapies are neutralised by the complex TME, responsible for the progression and metastasis of solid tumours^7–10^. Since Mϕ represent the largest immune population in solid tumours^11^, their reprogramming to their native state and direction against tumour cells is an attractive strategy which is still in its infancy. Most TAM-targeting approaches are based on a thorough molecular characterization of the immunosuppressive pathways involved in the TME^12–18^. Direct approaches include Mϕ depletion with CSF-1R inhibitors^19–21^, TAM “re-education” with CD40 agonists^22^, disruption of the PD-1/PD-L1 axis^23^, or antibody-based targeting of Mϕ-derived proteins^24^. A breakthrough in Mϕ-mediated immunotherapy was defined by the discovery of the “don’t eat me” signal CD47 overexpressed on the surface of cancer cells to evade immune surveillance. The CD47/SIRP-⍺ axis is an immune checkpoint disrupted by Magrolimab^®^, a CD47 monoclonal antibody that promotes tumour phagocytosis^25^. The molecule progressed to phase III clinical trials for the treatment of hematological malignancies but was discontinued, highlighting the enormous challenges in the field^26^. A variety of immunoconjugates able to alter the immune system have been proposed to modulate the TME^27–34^. Other approaches focus on the disruption of hypersialylated cancer cells engaging the inhibitory lectin receptors (Siglecs) and glycoimmune checkpoint blockade^35–37^.

Herein, we describe a novel class of modular heterobifunctional molecules termed phagocytic synapse enhancers (PSEs). PSEs bind selectively to tumour-associated antigens harbouring the cancer cell surface with an “*eat me*” signal that triggers and directs the phagocytic activity of Mϕ specifically towards cancer cells. Since immunosuppressive Mϕ express immune checkpoints (e.g. PD-L1) similar to cancer cells, PSEs enhance the uptake of membrane and extracellular targets via the endolysosomal pathway. Given their low molecular weight (ca. 14 kDa), PSEs exhibit improved tissue penetration to better reprogram tissue-resident Mϕ in the TME. This technology, termed ENPHASYS (<u>En</u>hancement of <u>Pha</u>gocytic <u>Sy</u>napse<u>s</u>), is highly modular and can be used to target any cancer surface antigen while providing a tumour-directed immunostimulatory signal.

## METHODS

### Protein expression and purification

dnPDL1 and dnEGFR were provided by the Institute of Protein Design, University of Washington. The recombinant proteins were obtained according to literature protocol^38^.

The recombinant plasmids for nbPDL1 were transformed into Shuffle T7 competent *E. coli* (New England Biolabs: C3026J) and an overnight pre-culture of these were grown in LB media (5 mL) containing ampicillin (50 μg/mL). Pre-cultures were used to inoculate 0.5 L of LB medium containing 50 μg/mL of ampicillin in 2 L flasks to reach an initial OD_600_ of 0.02. These were incubated at 37 °C, 180 rpm until an OD_600_ of 0.8 was reached. At which point, expression was induced upon addition of IPTG (250 μM). The temperature was reduced to 30 °C and the cultures left to incubate overnight at 180 rpm. The expression media was subsequently collected after centrifugation at 8000 rpm and 4 °C for 30 min. The pellet was discarded, and the supernatant filtered through a 0.45 μm filter under moderate vacuum to remove cellular debris. The supernatant was loaded onto Ni-NTA resin equilibrated with PBS (pH 7.4) and passed over the resin 3x before the bound protein was eluted with an imidazole gradient The imidazole gradient consisted of 10 mM (30 mL), 40 mM (15 mL), 200 mM (5 mL) and 200 mM (10 mL) (all imidazole solutions were in PBS pH 7.4). The fractions were collected and analysed by SDS–PAGE before combining pure protein containing fractions. Following combination of fractions, all samples were desalted using HiPrepTM 26/10 Desalting column (Cytiva) eluting with PBS (pH 7.4). Protein concentrations were determined by absorbance at 280 nm using a NanoDrop 2000c UV-Vis spectrophotometer. All protein sequences are reported in the Supplementary Information,Table S4.

### Bioconjugation

Fresh batches of cysteine-containing binders in PBS (pH 7.4) were reduced using TCEP (2.0 equivalents) to remove glutathione and homocysteine capping occurred during expression, followed by addition of maleimide linkers (5.0 equivalents) to reach full conjugation monitored by LC-MS analysis. After conjugation, purification was carried out by SEC using a HiLoad 10/300 Superdex 75 column (Cytiva, Little Chalfont, UK) and PBS (pH 7.4) as an elution buffer.

### SDS-PAGE gel

For SDS-PAGE gels, 2 μg of conjugates with SDS dye were loaded onto a Bis-Tris 4–12% Criterion™ XT Bis-Tris Protein Gel, 18 well, (Bio-Rad 3450124) and run with XT-MES buffer at 180 V for 40 min. Proteins were stained with Aquastain (Bulldog Bio AS001000) for 10 min followed by a 10 min destain in water.

### General LC–MS protocols

LC–MS analysis of protein samples was carried out using a Waters SQD2 mass spectrometer using inlet methods A and B, or a Waters Xevo G2-S TOF mass spectrometer using inlet method C, in combination with an Acquity UPLC system with an Acquity UPLC BEH300 C4 column (130 Å1.7 μm, 2.1 × 50 mm). The SQD2 mass spectrometer mobile phase consisted of solvent A (99.9% water with 0.1% formic acid), solvent B (71% acetonitrile, 28.9% water and 0.1% formic acid) and the following gradients were programmed. Inlet method A: 100% A for 2 min, then 100% A to 100% B in 9 min, then 100% B for 5 min followed by 100 % A for 4min. Inlet method B: 100% A for 2min, then 85 % A to 100% B in 6 min, then 100 % B for 2.5 min followed by 85% A for 3.5 min. The Xevo G2-S TOF mass spectrometer mobile phase consisted of solvent A (99.9% water with 0.1% formic acid), solvent B (95% MeCN and 5% water with 0.1% formic acid) and the following gradients were programmed. Inlet method C: 95% A for 0.93 min, then 95% A to 100% B in 4.28 min, then 100% B for 1.04 min, 100% B to 95% A for 1.04 min. The capillary voltage of the electrospray source for the Waters SQD2 mass spectrometer was 3.0 kV with a cone voltage of 30 V and the desolvation gas used was nitrogen, with a flow rate of 800 L h−1. For the Waters Xevo G2-S TOF mass spectrometer the capillary voltage of the electrospray source was 2.0 kV with a cone voltage of 40 V and the desolvation gas used was nitrogen, with a flow rate of 850 L h−1. The ion series was obtained through integration of the major peaks of the chromatogram. Following this, the total mass spectra were reconstructed using the MaxEnt1 algorithm on the MassLynx software (v. 4.1), according to manufacturer’s guidelines.

## In vitro studies

### Cell culture

All cell lines were purchased from the American Type Culture Collection (ATCC). RAW 264.7 and J774A.1 cells were cultured in DMEM + 10% heat-inactivated fetal bovine serum (FBS) with antibiotic selection. MC-38 colon carcinoma cells were cultured in DMEM + 10% heat-inactivated FBS, supplemented with HEPES, pyruvate, non-essential amino acids and antibiotic selection. Cell lines were not independently authenticated beyond the identity provided from the ATCC. Cell lines were cultivated in a humidified incubator at 5% CO_2_ and 37 °C and tested negative for mycoplasma using a PCR-based assay.

### Isolation of peripheral blood mononuclear cells

Peripheral blood leukocytes from healthy donors were obtained from the Blood Donation Center of the University Hospital Basel, Switzerland. Informed consent was obtained from all participants before blood collection. Peripheral blood mononuclear cells (PBMCs) were isolated using Ficoll Paque-PLUS (GE17-1440-02, Cytiva, Germany) and density centrifugation. After two rounds of ACK-lysis (#A10492-01, Gibco, USA) to remove erythrocytes, PBMCs were washed with PBS. Monocytes were magnetically separated from PBMCs by positive selection using human CD14 MicroBeads (#130-050-201, Miltenyi Biotec, Germany). Primary monocytes were taken fresh after magnetic separation from PBMCs and plated at 0.8 x 10^6^ cells per mL in RPMI 1640 (#61870036, Gibco, USA) supplemented with 10% heat-inactivated fetal bovine serum (#P30-3302, PAN-Biotech, Germany), 5% human serum (#H4522, Sigma-Aldrich, USA) and 25 ng/mL Mϕ colony-stimulating factor (M-CSF, #400-25, PeproTech, USA). After 3-4 d at 37 °C and 5% CO_2_, the medium was replaced with medium lacking human serum and refreshed every 2-3 d for a maximum of 14 d until use for downstream assays.

### Isolation and differentiation of bone-marrow derived Mϕ

After removing all muscles from the legs of ≥8-week-old mice, femur and tibia bones were isolated and flushed out with sterile PBS containing 2% FBS. Bone marrow cells were filtered through a 40 μm strainer (#352340, Falcon) and spun down for 5 min at 500 g. Cell pellets were incubated in Red Cell Lysis Buffer (154 mM NH_4_Cl, 10 mM KHCO_3_, 0.1 mM EDTA) for 3 min and then filled with PBS containing 2% FBS. After a 5 min centrifugation at 500 g, cells were resuspended in Complete RPMI (10% FBS, Glutamax, P/S). In the presence of 10-20 ng/mL Recombinant Mouse M-CSF (#576406, Biolegend). Bone marrow cells were differentiated into Bone Marrow-derived Mϕ (BMDMs). Differentiation was started at d0, the same amount of Complete RPMI containing M-CSF was added at d3, and at d6 differentiation medium was removed and exchanged. M1-like (50 ng/mL IFN-γ, 100 ng/mL LPS) or M2-like (10 ng/mL IL-4 and IL-13; # 214-14 or #210-13, Peprotech) polarization started at d6 or 7.

### Cell binding avidity assays

Tumour cells were attached on poly-L-lysine-coated chips for at least 3h prior to testing on the z-Movi Cell Avidity Analyzer (Lumicks). CellTrace far-red labelled (ThermoFisher Scientific) Mϕ treated with GRD peptide (50 mM), were flown inside the chip and, in the presence of the PSEs, incubated with the target tumour cells for 3 min prior to application of acoustic force linearly from 0 to 1000pN. Experiments were performed with RAW 264.7 Mϕ and IFNγ treated MC-38 cells. Cell detachment was analysed using Ocean software. Experiments and analysis were conducted according to manufacturer recommendations.

### Immunofluorescence staining assay

MC-38 cells were cultured as mentioned above. According to the type of assay, the cells were plated in a 96-well plate (20’000 cells, 100 μL) and stimulated with IFNγ (20 ng/mL) 24h prior to the assay. After stimulation, the fluorescent probes (100 nM) are added to the well and incubated for 1h at 37 °C. At the endpoint, the cells are washed with PBS buffer (100 μL x 3) and the cells lifted with TrypLE, centrifuged, resuspended in FACS buffer and analysed by flow cytometry (Cytoflex, Beckman Coulter) to determine histograms and the fluorescence intensity of the cells. Data were analysed with FloJo software. For confocal microscopy analysis, cells were added to poly-L-lysine coated 96 well plates and at the endpoint the cells were washed with PBS buffer (100 μL x 3) and fixed with 4% PFA. Cells were then imaged with a Leica Stellaris confocal microscope. Data were acquired from a 63× oil immersion objective lens, a green fluorescence channel (ex: 460 ± 10, em: 524 ± 10, acquisition time: 300 ms) or yellow-orange fluorescence channel (ex: 565 ± 10, em: 578 ± 10).

### Characterization of PSEs affinity to PD-L1 by BLI

All BLI experiments were performed on an Octet QKe (ForteBio) at 25 °C. Proteins for analysis were prepared in sample buffer (PBS, pH 7.4, 0.02% (v/v) Tween-20 and 0.05% (w/v) bovine serum albumin). The sensorgrams were obtained with a 60 s baseline, 100 s loading, 60 s baseline, 180 s association, and 200 s dissociation. Biotinylated PD-L1-Fc fusion protein (Invitrogen) was diluted to 2 μg/ml in sample buffer and loaded onto Octet Streptavidin biosensors (Sartorius). Individual purified nbPDL1/dnPDL1-conjugates were diluted in sample buffer to between 400 nM and 12.5 nM. Raw data were corrected with double referencing and analysed using Prism v10.0 (GraphPad). The dissociation constants (*K_D_*) for PD-L1 binding were derived from steady-state equilibrium analysis of the sensorgrams using the maximum response observed during association of the PSEs binding to PD-L1 Fc. The binding curves were fitted in Prism to a one-site specific-binding model. Additional kinetic analysis was performed on the sensorgrams to determine k_on_, k_off_, and *K*_D_ values, using the global analysis model for association kinetics within the Prism9 software.

### Phagocytosis assays

Mϕ were washed with PBS and lifted by mechanical pipetting with cold PBS buffer to avoid enzymatic digestion of the surface receptors. Mϕ were pelleted by centrifugation at 300 × g for 5 min and resuspended in PBS buffer containing CMFDA Green and incubated at 37 °C for 30 min. Stained Mϕ (20,000 cells, 100 μL) were added to a 96 well flat-bottom plate (Corning) and incubated in a humidified incubator. Meanwhile, target cells were washed 1x with PBS, detached with TryplE (1 mL) and centrifuged at 300 x g for 5 min. Then cells are resuspended in PBS buffer containing Cell Trace far red dye (Invitrogen) in PBS at 37 °C for 30 min, washed with PBS, centrifuged at 300 x g for 5 min and resuspended in growth medium. The stained MC-38 cells (100 μL, 20’000 cells) are then added to the 96-well plate containing Mϕ and incubated for 30 min at 37 °C. Finally, the media was removed from the wells and a solution of PSEs (100 μL, 100 nM) in serum-free medium is added to the co-culture and incubated at 37 °C for 4 h.

### Confocal microscopy

For phagocytosis assays, cells in co-culture were fixed with 4% PFA after 4 h incubation. Images of fixed cells were taken with a Leica Stellaris 5 confocal microscope. Data were acquired from a 63× oil immersion objective lens at 1x magnification, a green fluorescence channel (ex: 460 ± 20, em: 524 ± 20, acquisition time: 300 ms), and a far-red fluorescence channel (665 ± 40, acquisition time: 400 ms). Two images per well were acquired at the end point. Tiles were acquired using a z-stack of 35 μm and images processed by ImageJ (Fiji). The number of phagocytised cancer particles and the average MFI was calculated by processing the images on QuPath (Cell detection: Channel 1; requested pixel size: 0.5 μM; Background radius: 8 μm; sigma: 1.5; minimum area: 10 μm; maximum area 400 μm^2^; Threshold: 13.49. Channel 2: Threshold: 22.78).

### Flow cytometry analysis

Quantitative analysis of the phagocytosis assay was performed by flow cytometry (Cytoflex, Beckman Coulter). Phagocytosis assay is stopped at the endpoint by adding cold FACS buffer to the wells and placing the 96-well on ice. The cells are transferred to an Eppendorf tube and acquired. FLowJo and gated on intact cells and single cells to determine the percentage of double positives Macs PE^+^APC-750^+^. Phagocytic index is calculated by gating on Mϕ only and analysing the ratio of PE^+^APC-750^+^ Mϕ against total Mϕ (PE^+^ Mϕ + PE^+^APC-750^+^ Mϕ). Data analysis was performed on Prism v10 (GraphPad, USA).

### Imagestream analysis

THP-I cells were differentiated to Mϕ with PMA (50 nM) for 48h in a cell culture treated 24-well plate. After differentiation, PD-L1 expression was induced for 24h with IFNγ (20 ng/ml) and cells treated with the PSEs or relative controls and incubated at 37 °C. At the end of the time point, cells were washed three times with PBS and stained for PDL1 with aPDL1-APC (BioLegend Cat. N. 329707) antibody for 30 min at 4 °C). Finally, cells were washed twice, stained with DAPI for 20min at 4 °C, washed twice with PBS and the single-cell suspensions transferred to 1.5 mL Eppendorf tubes for analysis by Amnis Imagestream. For co-localisation with the lysosomes, Mϕ were pre-treated Lysobrite Red (AAT Bioquest, 22645) following the manufacturer’s protocol. Acquisition settings: Illumination intensity: 405 (20 mW), 488 (100 mW), 561 (50 mW), 642 (50 mW); Magnification: 40X; Speed: low). Cells were gated on Scatter plot of Area vs Aspect Ratio of Brightfield. Live cells were gated on Scatter plots of channel 7 relative to DAPI.

### Western Blot analysis

1×106 THP-I cells were plated 24h before the experiment. Cells were incubated with complete growth medium supplemented with IFNγ (20 ng/ml). On the day of the experiment, cells media was replaced with serum-free media containing the PSE, dnPDL1 or PBS and incubated for 4h and 24h. At the of the assay, the supernatant was removed and the cells lysed using RIPA cell lysis buffer supplemented with protease inhibitors cocktail and phosphatase inhibitors on ice for 30 min. The cells were centrifuged at full speed (13’000 rpm) for 15 min at 4 °C. The supernatant collected, and the protein concentrations were quantified using BCA assay. The supernatants for each condition were split in equal amounts and supplemented with Laemmli buffer (4x)

An equal amount of protein (about 30 μg, determined by BCA assay) was loaded per lane and separated on SDS-PAGE gels (SurePage Bis-Tris, 4-20%, 15-well, Tris-MOPS-SDS running buffer), and then transferred onto nitrocellulose membrane (BioRad TransBlot Turbo Transfer pack). Membranes were blocked for 1h with 5% non-fat dry milk in TBS supplemented with 0.05% Tween-20 (TBST) at room temperature for 1h and then probed with the primary antibody (anti PD-L1, Cell Signaling Technology, N.13684, or anti Actinϕ, BD Biosciences, N. 612657, 1:1000 in blocking buffer) at 4 °C overnight. After three times washing with TBST, secondary antibody (anti-goat or anti-rabbit IgG HRP-conjugated, 1:15000 dilution in blocking buffer) is added to the membrane for 1h at room temperature. All membranes were washed three times and exposed using ECL substrate (West) and imaged at the Fusion X Vilber.

### Statistical Analysis

Statistical analysis was performed in Prism (version 10) and Rstudio. For binding curves, one-site-specific binding curves were used to calculate the dissociation constants of binders and PSEs. In binding assays, phagocytosis experiments, ordinary one-way ANOVAs were performed with Tukey’s multiple comparison’s test to compare treatment groups. In every instance the asterisk * indicates a p<0.05, ** indicates p<0.01, *** indicates p<0.001, and **** indicates p<0.0001.

## Zebrafish Studies

### Zebrafish care and handling

Zebrafish (Danio rerio) model was handled and maintained according to the standard protocols of the European Animal Welfare Legislation, Directive 2010/63/EU (European Commission, 2016) and Champalimaud Fish Platform. All protocols were approved by the Champalimaud Animal Ethical Committee and Portuguese institutional organizations — ORBEA (Órgão de Bem-Estar e Ética Animal/Animal Welfare and Ethics Body) and DGAV (Direção Geral de Alimentação e Veterinária/Directorate General for Food and Veterinary).

### Zebrafish transgenic and mutant lines

According to the purpose of each experiment, different genetically modified zebrafish lines were used in this study: Tg(fli1:eGFP), Tg(mpeg1:mcherry-F) and Tg(mpeg1:mcherryF, tnfa:eGFP-F).

### Cell Culture

TNBC cell line Hs578T was cultured and expanded in Dulbecco’s Modified Eagle Medium (DMEM) High Glucose (Biowest) supplemented with 10% Fetal Bovine Serum (FBS) (Sigma-Aldrich), 1% Penicillin-Streptomycin (P/S) 10,000 U/mL (Hyclone) and supplemented with insulin at 10 g/mL (Sigma-Aldrich). Hs578T cells were cultured in a humidified atmosphere containing 5% CO_2_ at 37°C. Hs578T were authenticated through short tandem repeat (STR) profile analysis and tested routinely for mycoplasma contamination.

### Cell Labelling

Hs578T cells were grown to 70% confluence, washed with Dulbecco’s phosphate-buffered saline (DPBS) 1X (Biowest) and detached with 1 mM EDTA by scrapping. Cell suspension was collected to 1.5 ml eppendorf, stained with lipophilic dye Deep Red Cell Tracker (1 μl/ml in DPBS 1X, 10 mM stock) (Life Technologies), for 10 min at 37 °C, in darkness and washed with DPBS. Cells were centrifuged at 250 × g, for 4 min at 4 °C, and resuspended in DMEM. Cell viability was assessed by trypan blue exclusion method, and cell number was determined by hemocytometer counting. Cells were resuspended in growth medium to a final concentration of 0.50 × 10^6^ cells/μl. Maximum tolerated concentration assay to assess the maximum tolerated concentration (MTC) of each compound in zebrafish larvae, a MTC assay was performed using the in vitro viability assays as reference for the tested concentrations. Groups of 20 noninjected zebrafish larvae were exposed to different concentrations of the compounds (dnPDL1, shortPSE and lonPSE) during 3 consecutive days, with the E3 medium/drug being renewed every day. The tested concentrations of the compounds, in different combinations, were the following 0.1 μM, 0.5 μM and 1 μM. Toxicity was analysed daily by counting the total number of dead larvae and checking for the presence of morphologic changes such cardiac edemas or curved tails.

### Zebrafish xenografts injection

Fluorescently labelled Hs578T cells were injected using borosilicate glass microcapillaries under a fluorescence scope (Zeiss Axio Zoom. V16) with a mechanical micropipette attached (World Precision Instruments, Pneumatic Pico pump PV820). Cells were injected into the perivitelline space (PVS) of 2 days post fertilization (dpf) zebrafish embryos, previously anesthetised with Tricaine 1X (Sigma-Aldrich). After injection, zebrafish xenografts remained for ∼10 min in Tricaine 1X and then transferred to E3 medium and kept at 34 °C. At 1 dpi, zebrafish xenografts were screened according to the presence or absence of tumoral mass. Xenografts with cells in the yolk sac or cellular debris were discarded, whereas successful ones were grouped according to their tumour size, which was classified by comparison with eye’s size. Next, xenografts were exposed to E3 medium (controls) and to the treatment groups: dnPDL1, shortPSE, longPSE. Xenografts were checked daily, and dead ones removed and E3 medium with the different compounds refreshed. Four days after zebrafish xenografts were sacrificed, fixed with 4% (v/v) Formaldehyde (FA) (Thermo Scientific) at 4 °C overnight and preserved at −20°C in 100% (v/v) methanol. For transgenic zebrafish line Tg(mpeg1:mcherry-F) fixation was performed with PIPES for optimal fluorescent signal preservation.

### Zebrafish Xenograft Drug administration

Previous work from the lab showed that, when the treatment compounds are antibodies, adding them to the cell suspension prior to injection increases the effectiveness of the compound. As such, each compound was added to the cell suspension at the same concentration of 0.5 μM. At 1 dpi all zebrafish xenografts were screened regarding the presence of successfully injected tumour mass and distributed in the corresponding group controls (E3 medium) and treatment groups: dnPDL1, shortPSE and longPSE.

### Zebrafish xenografts imaging and analysis

Fixed zebrafish xenografts images were acquired using a LSM 980 Upright confocal laser scanning microscope, with a 5 μm interval. Generated images were processed using the FIJI/ImageJ software. The number of cells was quantified with ImageJ software Cell counter plugin. To assess tumour size, three representative slices of the tumour, from the top (Zfirst), middle (Zmiddle), and bottom (Zlast), per z-stack per xenograft were analysed and a proxy of total cell number of the entire tumour (DAPI nuclei) was estimated as follows: the 1.5 correction number was estimated to human cells that have a nucleus with an average of 10–12 μm of diameter.

### Confocal imaging and analysis of zebrafish xenografts

Mounted xenografts were imaged using an Andor BC43 spinning disk confocal microscope. Tumours were imaged with a 20× objective lens using the z-stack function with an interval of 5 μm between slices. The number of cells was manually assessed with the Cell Counter plugin from ImageJ/Fiji. To assess tumour size, a proxy of the total cell number (DAPI nuclei) was estimated by counting the number of nuclei in three representative slices of the tumour from the top (Zfirst), middle (Zmiddle) and bottom (Zlast) per z-stack per xenograft, as follows:

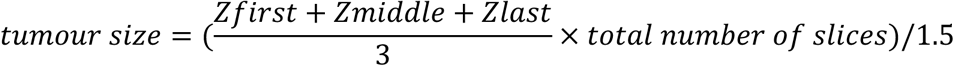

The 1.5 correction number was estimated for human cells that have a nucleus with an average diameter of 10-12 μm. The numbers of Tnfa-positive/negative macrophages were individually quantified in every slice along the tumour. To get the percentage of each population, the obtained number was divided by its corresponding tumour size.

### Statistical Analysis

Statistical analysis was performed using the GraphPad Prism 10.0 software. All data sets were challenged by D’Agostino Samp; Pearson and Shapiro–Wilk normality tests. In general, data sets with a Gaussian distribution were analysed by parametric unpaired t test and data sets that did not pass the normality tests were analysed by nonparametric unpaired Mann–Whitney test. Clearance data sets were analysed using Fisher’s exact test. All were two-sided tests with a confidence interval of 95%. Differences were considered significant at P<0.05 and statistical output was represented as follows: non-significant (ns) ≥0.05, *p<0.05, **p<0.01, ***p<0.001, ****p<0.0001. The bars indicate the results as AVG ± standard deviation of the mean (STDEV).

## Mouse Studies

### Cell culture for MC-38 murine model

C57BL6 murine colon adenocarcinoma cell MC38 was purchased from Kerafast (ENH204-FP). The cell was cultured in Dulbecco’s modified MEM (Gibco) with 10% fetal bovine serum (FBS) (Gibco, Thermo Scientific), 1x GlutaMAX (Gibco, Thermo Scientific), 1x penicillin/streptomycin solution (Gibco, Thermo Scientific). The cell was cultured at 37 °C in humidified condition with 5% CO2.

### In vivo MC-38 colorectal cancer mouse study

All animal experiments were conducted at the Instituto de Medicina Molecular João Lobo Antunes (IMM, Lisbon). Animal work was performed in strict accordance with Portuguese Law (Portaria 1005/92) and the European Guideline 86/609/EEC and follow the Federation of European Laboratory Animal Science Associations guidelines and recommendations concerning laboratory animal welfare. All animal experiments were approved by the Portuguese official veterinary department for welfare licensing – Direção Geral de Alimentação e Veterinária (DGAV) and the IMM Animal Ethics Committee (authorization AWB_2021_03_GB_TargCancerDrugs).

8-10-week-old female C57BL/6J mice (purchased from Charles River) were used in this study, with 1×10^6^ MC38 cells inoculated subcutaneously in the flanks of mice. Tumour growth was monitored over time, by performing bilateral vernier caliper measurements every day and mean tumour volumes were calculated using the formula (length x width^2^) /2. Treatments were initiated when tumours reached approximately 90-100 mm^3^ (approximately 7-8 d after tumour induction), with the mice been randomly assigned to receive longPSE, dnPDL1, Magrolimab, Peptide and PBS as controls. Treatments were administered intratumourally in a total of 3 injections for every 3 d. Animals were observed every day; tumours were measured as described before and mouse weight was evaluated throughout the study. Mice were sacrificed when tumour volume reached 1,000 mm^3^, or body weight loss exceeded 20%, or tumours ulcerated. The data collected was analysed using GraphPad Prism v10.0.

### In vivo GBM mouse study

The experiments involving mice comply with ethical research regulations using animals from the Veterinary Office of the Health Department of Canton Basel-Stadt and were performed under license #2929_31795.

### Orthotopic Glioma cell inoculation

Murine GL-261-L2t glioblastoma cells were orthotopically injected into the right hemisphere of adult (8–12-week-old) C57BL/6 mice. Mice were anesthetised using 2.5% Isoflurane (Attane™ Isoflurane ad us. vet., Piramal Pharma Limited India) in an induction chamber. During the surgery, anesthesia was maintained at 1.5% Isoflurane administered through a nose adaptor. The corneas were protected by applying artificial tear ointment (Lacrinorm®, Bausch&Lomb Swiss AG, 6301 Zug) and the surgery site was disinfected using betadine. The scalp was incised by 1 cm long median-sagittal cut, and the burr hole site was located using the Bregma as a landmark (1mm posterior, 3mm lateral from the Bregma). After performing a burr hole using a handheld drill, 1×10^5 GL-261-L2t cells were injected into the right hemisphere at a depth of 3mm below the dura surface using a blunt-ended syringe (701N, 26s/2”/2,10 μl, Hamilton) into a pocket created by inserting and retracting the needle an additional 0.5mm. The tumour cells were injected at a concentration of 2.5×10^4 cells/μl in a total of 4 μl of Phosphate Buffered Saline (PBS) at a speed of 1 μl/min. After completing the injection, the needle was kept in place for 1 min and then slowly retracted in 50 μm steps. The scalp wound was closed with single-knot sutures and mice were monitored for postoperative complications for 30 min. For perioperative analgesia, 0.1mg/kg buprenorphine (Bupaq® P ad us. vet., VetViva Richter GmbH 4600 Wels, Austria) was administered 30 min prior to the surgery via s.c. injection. Sutures were removed 7d post-surgery.

### In vivo bioluminescent imaging (BLI)

Mice were imaged by BLI starting at 7d post tumour injection. Isoflurane anesthetised mice were injected intraperitoneally (i.p.) with 150 mg/kg D-luciferin (#LUCNA-1G, Goldbio) and imaged after 10 min using a Newton 7.0 BLI system (Vilber). A region of interest (ROI) was drawn around the head and quantitated on mean luminescence (photon count per pixel area). Mice were imaged weekly and checked for neurological symptoms and weighed daily from d 21 onwards.

### Evaluation of systemic toxicity after intratumoral single-shot treatment

GL-261 L2tdTomato tumour-bearing C57BL/6 mice were treated with a single injection of longPSE or PBS (4 μl, 1 μl/min) into the tumour site under Isoflurane anesthesia using a stereotactic frame and blunt-ended syringe (701N, 26s/2”/2,10 μl, Hamilton). Treated mice were monitored for symptoms of acute toxicity for 30 min post injection and then in 2 h intervals for 6 h. Blood samples were collected 24 h after treatment by tail vein blood draw. Mice were placed under a heat lamp for 2 min to achieve blood vessel dilation and subsequently restrained in a restraining tube. The tail vein was incised using a surgical blade and 2 x 10 μl of blood were withdrawn and immediately mixed with 120 μl 1mM EDTA in PBS. The tail vein was compressed until haemostasis was achieved and mice were placed into a surveillance cage and monitored for 10 min. Whole blood samples were immediately analysed using a haemocytometer (XN-1000V, Sysmex). Parameters used to detect haematological toxicity include WBC, RBC, MCV, MCH, and PLT. Results were analysed using the Graph Pad Prism software v10.0 (GraphPad, USA).

### Alzet® mini-osmotic pump implantation

Alzet® mini-osmotic pumps (model 2004, 0.25 μl/h, 28 d, DURECT Corporation, Cupertino, CA 95014. Lot specific performance: Lot no. 10435-22; Mean pumping rate 0.26 μl/h, SD 0.01 μl/h; Mean fill volume 232.9 μl, SD 5.7 μl) were loaded with longPSE, dnPDL1, Atezolizumab and PBS control solutions under sterile conditions using a 1 ml syringe and connected via flow moderator and polyvinylchloride catheter tube to a brain infusion cannula (Alzet® brain infusion kit 3, Lot no. 10445-23, DURECT Corporation, Cupertino, CA 95014) according to the manufacturer’s instructions. Prefilled pumps were placed in 0.9% saline for 40 h at 37°C before implantation for priming. C57BL/6 mice were anesthetised with Isoflurane (Attane™ Isoflurane ad us. vet., Piramal Pharma Limited India) and mounted onto a stereotactic frame. The eyes were protected from drying out by application of artificial tear ointment (Lacrinorm®, Bausch&Lomb Swiss AG, 6301 Zug) and the surgery site was extensively disinfected with betadine. Unsterile areas were covered with sterile surgical drapes. All instruments were autoclaved before use and the surgery was performed using sterile surgical gloves. The scalp was incised by 1 cm long median-sagittal cut. The skull was dried and cleaned of periost connective tissue with a sterile cotton swab. Entering through the skin incision, an arterial clamp was used to form a pocket for the pump at the height of the scapulae. After inserting the pump into the pocket, the brain infusion cannula was introduced into the tumour through the same burr hole previously used for tumour cell inoculation and fixed to the skull bone with dental cement (Hoffmann’s quick setting Zinc Phosphate Cement, Hoffmann Dental Manufaktur GmbH, 12099 Berlin). The skin incision was closed with single-knot sutures (5-0 Prolene™ Polypropylene non-absorbable monofil sutures, Johnson&Johnson Medical GmbH, 22851 Norderstedt, Germany). Mice were transferred into a wake-up cage placed on a 37°C electrical heating mat and monitored continuously for the first 30 min after surgery. Fully awake animals were put back to the housing cage and checked for postoperative complications after 2 h. Mice with implanted Alzet® pumps were housed in single cages and stitches were removed after 7 d. For perioperative analgesia, 0.1mg/kg buprenorphine (Bupaq® P ad us. vet., VetViva Richter GmbH 4600 Wels, Austria) was administered 30 min prior to the surgery via s.c. injection.

### Mouse tumour dissociation

Mice were euthanised by CO_2_-suffocation at endpoint or indicated time points and tumour-bearing cerebral hemisphere without cerebellum was harvested into ice-cold HBSS. On ice, brain tissue was manually minced using razor blades and enzymatically dissociated at 37 °C for 30 min using the same dissociation buffer as described above. The suspension was filtered through a 70 μm strainer and subjected to a density gradient centrifugation using debris removal solution (#130-109-398, Miltenyi Biotec) according to manufacturer’s protocol to remove myelin and cell debris. Following ACK-lysis, the remaining myelin- and erythrocytes-depleted tumour cell suspension was used in downstream applications.

### Transduction of mouse GBM cell lines

For in vivo tumour monitoring by bioluminescence imaging (BLI) and fluorescent labelling of tumour cells in flow cytometry-based phagocytosis assay, GL-261 were lentivirally transduced to express luciferase 2 (L2) and fluorescent reporter protein tdTomato (GL-261 L2-tdTomato). The lentiviral L2-tdTomato construct was a gift from Prof. Bentires (University of Basel). For lentiviral transduction, tumour cells were plated at 5 x 10^4^ cells per well of a 24-well plate 16 h prior to transduction. Media was replaced with 0.5 mL antibiotic-free media containing 8 μg/mL Polybrene (Sigma-Aldrich). 5 μL of lentivirus suspension was added to the cells and incubated for 6 h at 37°C. Afterwards, transduction medium was replaced with normal growth medium and cells were expanded for 4 d and positively sorted for tdTomato expression. Luciferase expression was checked by using *in vitro* bioluminescence imaging by adding 100 μL of 15 mg/mL D-luciferin solution (#LUCNA-1G, Goldbio) to the cells and imaged after 2 min using a Fusion FX system (Vilber).

## Spectral flow cytometry analysis

### Staining and acquisition

A 200 μL fraction of one sample per treatment group was used as an unstained control to detect and correct for condition-specific autofluorescence. In all cases, input volume for full-stained samples were kept at 200 μL to ensure equivalent staining conditions across all samples remaining 200 μL of SCS volume was mixed and used to generate fluorescence minus one (FMO) stainings, allowing for condition-specific FMO gating. All centrifugation and incubation steps were performed at 300 × g at 4°C, protected from light, unless otherwise stated. Viability staining was performed by incubating cells with Zombie Aqua Viability kit for 20 min. Subsequently, Fc-block was performed by incubating cells in a dilution of mouse TruStain FcX (#101320, BioLegend, USA) for 10 min. Antibody mastermixes (full-stains and FMOs) were freshly prepared on the day of staining in Brilliant Stain Buffer (#00-4409-75, ThermoFisher Scientific, USA). Cell surface markers were stained with surface antibody mastermixes for 25 min and then washed twice in autoMACS Running Buffer. To allow the subsequent staining of intracellular antigens, a fixation/permeabilization step was performed by incubating the cells for 20 min at RT using a Cyto-Fast Fix/Perm Buffer set (#426803, BioLegend, USA). Intracellular antibody staining was then performed for 25 min at RT. The spectral FC staining panel is summarised in **Table S1-2.** Finally, the samples were washed twice, resuspended in a final volume of 200 μL of PBS and acquired on a Cytek Aurora 5-Laser Spectral Analyzer (Cytek Biosciences, USA) using standard, daily quality-controlled, Cytek-Assay-Settings.

### FlowSOM statistical analysis

Data were manually pre-gated to remove the debris and select for CD45+, live, single cells using FlowJo v10 Software. The analysis was subsequently performed in R (version 4.4.1). Data was transformed by asinh transformation using variance stabilizing cofactors for each channel (estParamFlowVS and transFlowVS functions from the FlowVS package), except for the dTomato, Alexa Fluor 488, APC and BUV661 channel in the myeloid panel where the cofactor were manually set to 1500, 800, 800 and 800 respectively. In the lymphoid panel cofactors for tdTomato and PE were set to 1500 and 5000 respectively. Pre-processing QC (using PeacoQC function from the PeacoQC package) was performed to remove outliers and unstable events (IT_limit was set at 0.55 and MAD at 6). Clustering was performed using FlowSOM and ConsensusClusterPlus using the wrapper function cluster from the CATALYS package. The resulting clusters were manually annotated. Differential testing was performed using the diffcyt package (diffcyt-DA-edgeR for differential abundance and diffcyt-DS-limma for differential state).

### Sample size determination

Sample size was determined according to the prior work executed in the laboratory, or from published reports of similar scope within the appropriate fields.

### Replication

The number of biological replicates (typically n ≥ 3) is indicated in the figure legends. Key findings were replicated in at least two independent experiments except for the in vivo GBM model, which was carried out as single experiment.

## RESULTS

### ENPHASYS as a modular approach for targeted cancer therapy

We developed a modular approach to assemble PSEs against the desired target antigen. PSEs consist of three modules: the binder, the linker, and the phagocytosis-promoting peptide (**Fig. 1a–c**). The binder can be any molecule able to target a cancer antigen. For proof-of-concept, we focused on a de novo PD-L1 binder^38^ (dnPDL1) bearing a single cysteine available for bioconjugation (**Fig. 1a**). The low molecular weight of the PSE (ca. 14 kDa) improves its tissue penetration while directing the bispecific molecule to the TME. For bioconjugation (**Fig. 1b**) we employed maleimide-PEG linkers with three or one PEG_3_ units (longPEG and shortPEG) to provide flexibility to the bifunctional molecule. As phagocytosis-promoting peptide, we selected tuftsin, a tetrapeptide (TKPR) known to enhance phagocytosis (**Fig. 1c**) ^39–48^. The tuftsin sequence is conserved in the Fc region of IgGs and is liberated by enzymatic cleavage during IgG catabolism. Reports suggest that tuftsin acts as an Fc mimic binding to FcR, while other biological functions are attributed to its interaction with Neuropilin-1^41,42^. We validated tuftsin Mϕ-stimulating activity using Neutravidin Oregon Green 488 (NOG_488_) complexed to a biotinylated TKPR, leading to higher fluorescence uptake in RAW 264.7 Mϕ (**Extended data figure 1c-e**). This effect was reversed by the tuftsin antagonist TKPPR, showing that the uptake is tuftsin-mediated (**Extended data figure 1g-i).**

**Figure 1.**
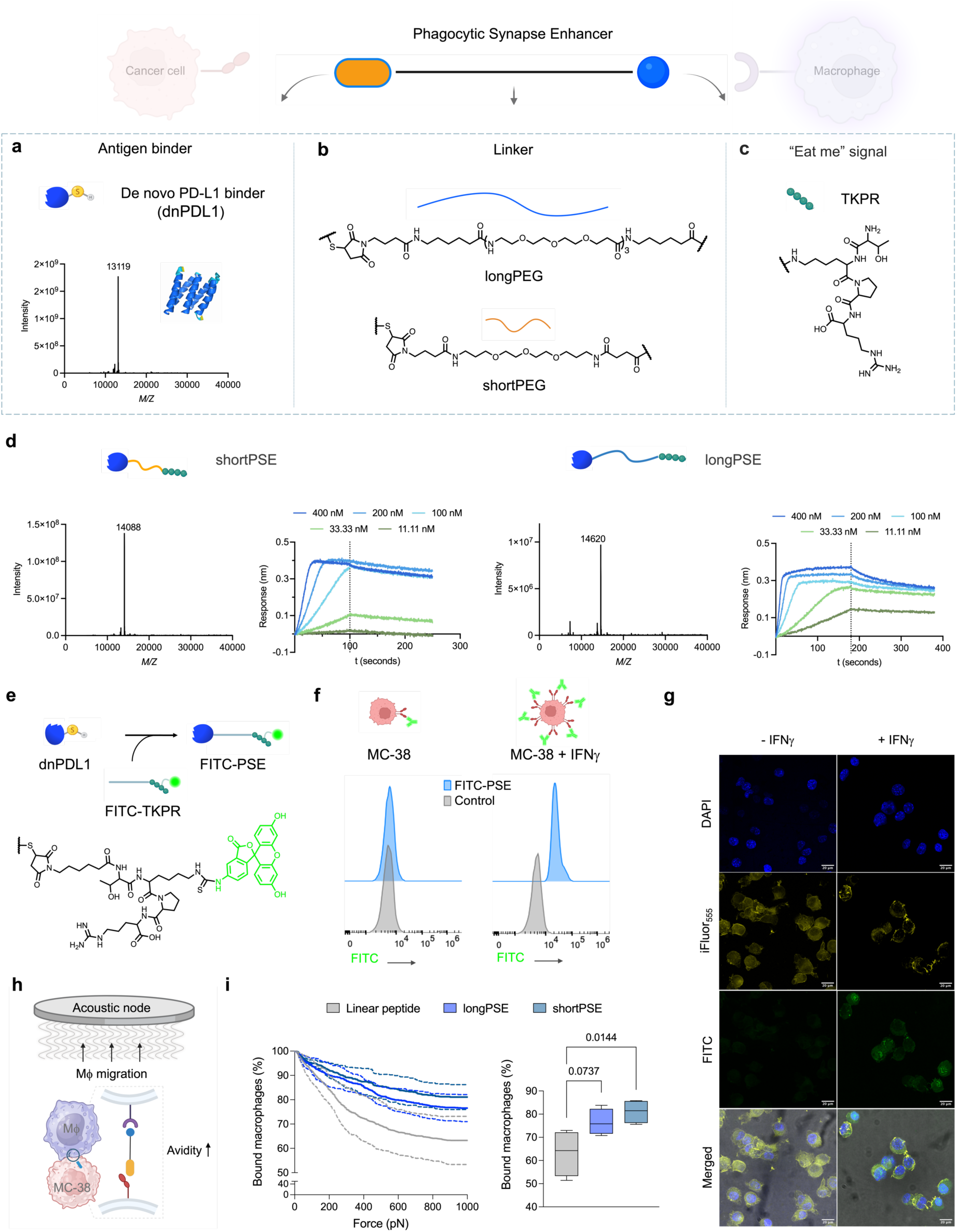
Modular approach to the synthesis of PSEs and functional characterization of the PSEs. **a,** The antigen binder is a single cysteine-bearing de novo mini protein of ca. 13 kDa with high affinity for PD-L1 (dnPDL1). The model was generated with AlphaFold 2.0. **b,** Maleimide PEG linkers of different length are used to tether the bifunctional PSEs. **c,** The phagocytosis-promoting peptide (TKPR) is coupled through the lysine side chain. **d,** Characterization of shortPSE and longPSE by LC-MS and biolayer interferometry analysis. Dissociation constants are calculated by non-linear regression analysis (shortPSE: K_D_ = 5.8 nM; longPSE K_D_ = 5.7 nM). **e,** Molecular structure of a FITC-based maleimide probe displaying the TPKR promoter. **f,** PD-L1 expression on MC-38 cells in the presence or absence of IFNγ. Histograms of the probe FITC-PSE bound to MC-38 cells determined by FC. Stimulation with IFNγ leads to overexpression of PD-L1 and higher shift in fluorescence intensity of the bound probes. **g,** Imaging of MC-38 cells labelled with the fluorescent probe FITC-PSE after 2h incubation. Panels from top to bottom: blue channel – nuclei stained with DAPI; yellow channel – cytoskeleton stained with Phalloidin iFLuor_555_; FITC channel – PD-L1 stained with FITC-PSE; merged channels with brightfield. Scale bar: 20 μm. **h,** Schematic representation of effector-target cell interaction and increase in avidity triggered by PSEs. **i,** PSEs (125 nM) with different linker length were incubated for 3 min in co-culture before application of acoustic force (0 to 1000 pN). Avidity assays show overall increased avidity with the PSEs compared to the control linear peptide (n=4). Statistical analysis was performed using ordinary one-way ANOVAs, followed by Tukey’s multiple comparison test to compare treatment groups. Illustrations created with Biorender.

Tuftsin is tethered to the linkers via functionalization of the lysine ε-amino group to yield the bifunctional molecules shortPSE and longPSE (**Fig. 1d**), characterized by LC-MS and octet biolayer interferometry. We confirmed that the bioconjugation step did not have a detrimental effect on the binding affinities (**Fig. 1d**), with K_D_ values in the low nanomolar range (shortPSE K_D_=5.7 nM, longPSE K_D_=5.8nM).

Next, we tested the binding of PSEs to cells. We synthesised a TKPR-linker displaying FITC (FITC-TKPR, **Fig. 1e, Extended data figure 2c**) to generate FITC-PSE (**Fig. 1e**) for FC and imaging studies with the murine colon carcinoma cell line MC-38 (**Fig. 1f-g**). The basal PD-L1 expression of MC-38 cells was not sufficient to appreciate a shift in fluorescence relative to the control, while stimulation with IFNγ led to PD-L1 overexpression and major shift in fluorescence (**Fig. 1f-g**). To localize the binding, we performed confocal microscopy (**Fig. 1g**). After 2 h incubation with the probe, the IFNγ-stimulated cells displayed higher fluorescence (**Fig. 1g**). This data, in combination with the FC analysis, confirmed FITC-PSE binding to PD-L1 on the cancer cell surface.

### PSEs increase the avidity between effector and target cells

Due to their bifunctional nature, we considered whether the PSEs could bridge Mϕ and cancer cells and enhance avidity by induced proximity, increasing the probability to initiate phagocytosis. The cell-binding strength of PSEs was assessed in co-culture with the *Z*-movi cell avidity analyser, an instrument that quantifies avidity by using acoustic forces (**Fig. 1h**). The assays were performed in a co-culture system with MC-38 cells and RAW 264.7 Mϕ. We performed an initial titration to identify the optimal concentration window for avidity (**Extended data figure 2d**). The binding strength increased from 12.5 nM to 1.25 μM to eventually drop – due to the Hook effect – once the concentration of the PSE reached 12.5 μM.^49^ We identified 125 nM as the optimal concentration to test the avidity of the PSEs. Both shortPSE and longPSE showed higher avidity compared to the linear peptide alone (**Fig. 1i and Extended data figure 2e**), suggesting that binding to the cancer antigen is crucial to augment avidity between the two cells. This represents a first methodology for avidity assays employing Mϕ as effector cells. Further optimization will lead towards the systematic screening of Mϕ engagers.

### PSEs enhance phagocytosis of colorectal cancer cells *in vitro* and extend the overall survival in a mouse CRC syngeneic model

We proceeded to evaluate the phagocytosis-enhancing properties of the PSEs. The murine cell line MC-38 (pre-treated with IFNγ to overexpress PD-L1) was used in co-culture with RAW 264.7 or J774A.1 Mϕ. PSEs enhanced phagocytosis by combining PD-L1 blockade^22^ and display of the “eat me” signal peptide (**Fig. 2a).** Co-culture assays with RAW 264.7 Mϕ indicated a higher phagocytic efficiency of the PSEs compared to controls (**Fig. 2b-f**). By confocal microscopy analysis (**Fig. 2b**), we observed that the control groups (**Fig. 2b**, panels A-C, respectively media, linear peptide, dnPDL1) induced moderate phagocytic activity. In contrast, PSE-treated co-cultures displayed a higher number of engulfed cancer cells and more pronounced phagosomes (**Fig. 2b**, panels D-E, respectively shortPSE and longPSE). We observed a similar phagocytosis trend by FC (**Fig. 2c-d**). While shortPSE had a modest effect compared to the single modules (Peptide, dnPDL1), longPSE mediated a >2-fold phagocytic increase compared to control. Increasing the concentration of longPSE to 1μM did not lead to a further increase in phagocytosis, suggesting the molecule reached its maximal effect (**Fig. 2e**). The phagocytic enhancement of longPSE was comparable to Magrolimab (**Fig. 2f**). Further in vitro assays were carried out with a larger library of PSE (**Extended data figure 2a-b**) displaying a variety of linkers, a cyclic version of tuftsin and a PD-L1 targeting nanobody (nbPDL1). We selected the best performing PSE, longPSE, to demonstrate the therapeutic efficacy of the ENPHASYS technology in a syngeneic mouse model of colorectal carcinoma (**Fig. 2g**). A subcutaneous flank model was engrafted with the same cell line studied in vitro, the colon cancer cell line MC-38. C57BL/6J mice were inoculated subcutaneously with MC-38 tumour cells to induce tumour and were treated intratumorally (i.t.) with longPSE, its unconjugated components (Peptide, dnPDL1), Magrolimab and PBS (**Fig. 2h**). The mice tolerated the molecules and did not show any body weight loss or drug-related toxicity (**Figure 2h**, **Extended data figure 6b**). Treatment with longPSE significantly reduced tumour growth compared to PBS and at the same time extended the overall survival, even when compared to Magrolimab (**Fig. 2h)**.

**Figure 2.**
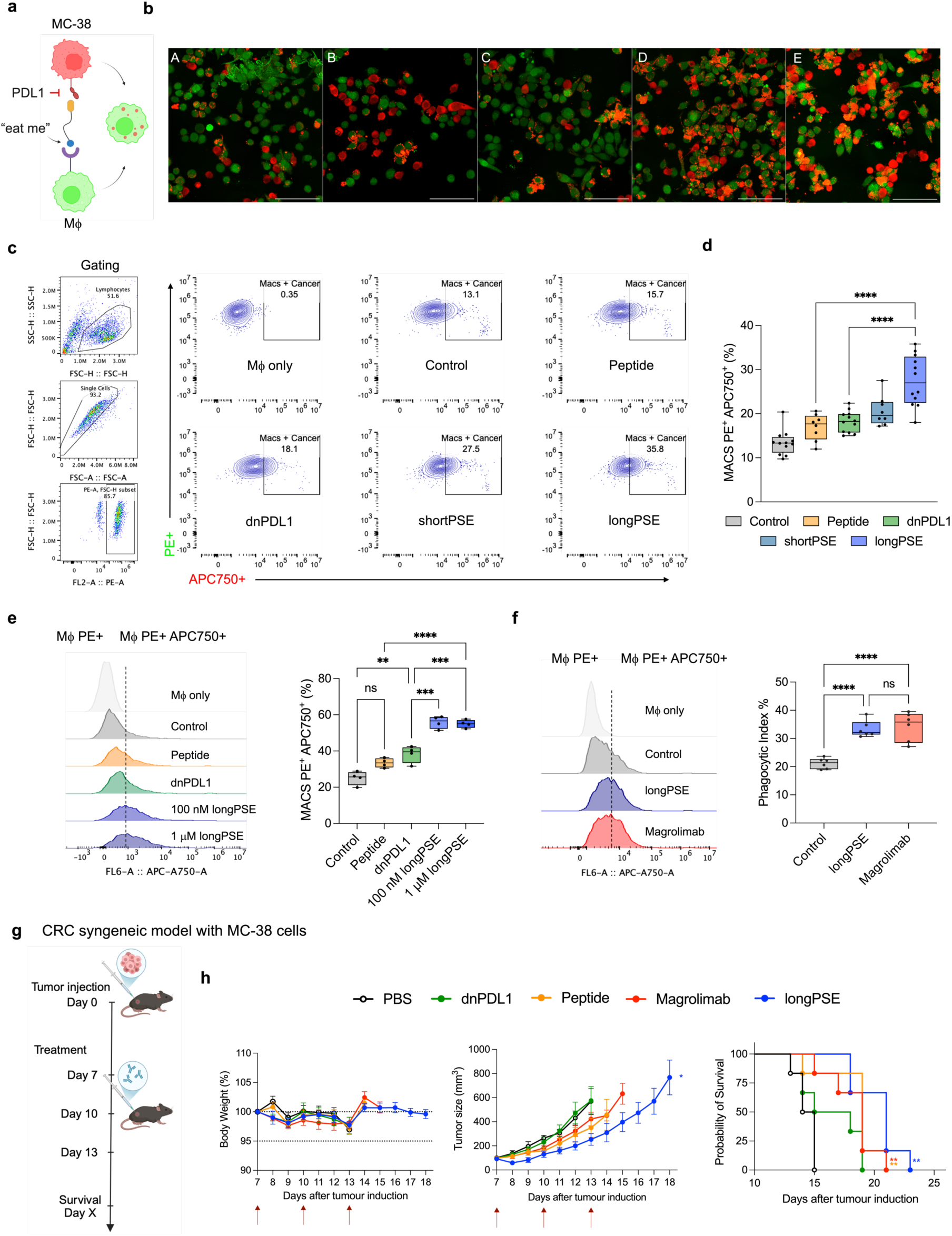
PSEs enhance phagocytosis in experimental CRC murine model *in vitro* and extend overall survival in syngeneic CRC mouse model. **a,** Schematic representation of a PSE targeting PD-L1 on the surface of tumor cells and displaying a phagocytosis-promoting peptide. **b,** Confocal imaging of co-culture assays. RAW 264.7 Mϕ (stained with CMFDA green) were co-cultured with MC-38 cancer cells (stained with CellTrace Far-Red proliferation kit) and treated for 4h with the respective molecules. Panels (A-E) correspond to co-cultures treated respectively with: control media, linear peptide, dnPDL1, shortPSE, longPSE. Scale bar: 100 μm. **c,** FC analysis after 4h incubation with different PSEs and controls showed an increase in the amount of double-positive PE^+^ APC750^+^ M. Cells were co-cultured in 1:1 ratio in tissue culture-treated 96-well plates. (Control vs shortPSE: p=0.0087; linear peptide vs longPSE: p<0.0001, n = 8 or 12). **d,** Phagocytic index of RAW 264.7 Mϕ in co-culture with MC-38 cells treated with the PSEs. The molecules were tested in co-culture (1:1 ratio) during 4h incubation. To determine the phagocytic index, double-positive Mϕ were pre-gated as PE^+^ Mϕ. **e,** FC analysis after 4h incubation with different concentration of PSE showed an increase in the amount of double-positive PE^+^ APC750^+^ Mϕ compared to relative to controls but no difference between the two different concentrations of PSE. Cells were co-cultured in 1:1 ratio in tissue culture-treated 96-well plates. (dnPDL1 vs 1μM longPSE: p=0.0008; dnPDL1 vs 100 nM PSE: p=0.0004, dnPDL1 vs Control: p=0.0082; peptide vs 1μM longPSE: p<0.0001, n = 4). Gating strategy and representative histograms for each condition are shown. **f,** FC analysis after 2h incubation of longPSE using Magrolimab as benchamark. Phagocytic index (%) is shown as the amount of double-positive PE^+^ APC750^+^ Mϕ pre-gated on PE^+^ Mϕ. Assay medium is used as control. Cells were co-cultured in 1:1 ratio in tissue culture-treated 96-well plates. (Control vs PSE: p<0.0001; Control vs Magrolimab: p<0.0001; n = 8 or 12). Representative histograms for each condition are shown. **g**, Experimental design of the in vivo study with syngeneic model of CRC. 1×106 MC-38 tumour cells were inoculated subcutaneously into C57BL/6J mice on d 0. Then, different treatments (longPSE, dnPDL1, peptide, Magrolimab or PBS) (n=6 per group) were administered i.t. on d7, 10 and 13. Mice were sacrificed when tumour volume reached ∼1000 mm^3^. **h**, from left to right, i) body weight changes during treatment with longPSE, peptide, dnPDL1, Magrolimab and vehicle (PBS). The body weight of each group of mice measured every day; ii) tumour growth curve. Data represents mean SEM of n=6. Multiple unpaired t test. *p<0.05; iii) overall survival of treated mice. Log-rank (Mantel-Cox) test indicates a statistically significant difference between treated mice and the control group. **p<0.01 (PBS vs longPSE: p=0.001; peptide: p=0.0095; Magrolimab: p=0.0036). Illustrations created with Biorender.

### PSEs enhance phagocytosis of mouse and human glioblastoma (GBM) cells in vitro and slow down the progression of glioblastoma in syngeneic models

To showcase their translational potential across different solid tumour entities, we tested the PSEs in GBM models. We used fluorescent reporter glioma cell lines (U-87 L2B, GL-261 L2tdTomato) expressing blue fluorescent protein (BFP) or tdTomato (tdTom) as a readout for phagocytosis. We initially tested the PSEs in co-culture with differentiated THP-I Mϕ (**Fig. 3a**). THP-I Mϕ were labelled with CellTrace Far Red and polarised for 24 h with IFNγ/LPS (pro-inflammatory) or IL-4 (anti-inflammatory). Pro-inflammatory Mϕ displayed a higher tendency to engulf cancer cells, although in both conditions the longPSE enhanced phagocytosis (M1-like: 26% vs 11% Control; M2-like: 15% vs 4% Control, **Fig. 3b-c**). Moving towards a less artificial co-culture system, we performed phagocytosis assays using bone marrow-derived Mϕ (BMDMs) and the glioma cell line GL-261 L2tdTomato (**Fig. 3d-e**). Again, longPSE had the highest phagocytic activity, this time comparable to the unconjugated dnPDL1 (**Fig. 3e**). In co-cultures of human peripheral blood mononuclear cells (PBMCs)-derived Mϕ with U-87 glioma cells (**Fig. 3f**), only longPSE showed significant phagocytic activity compared to control (**Fig. 3g**).

**Figure 3.**
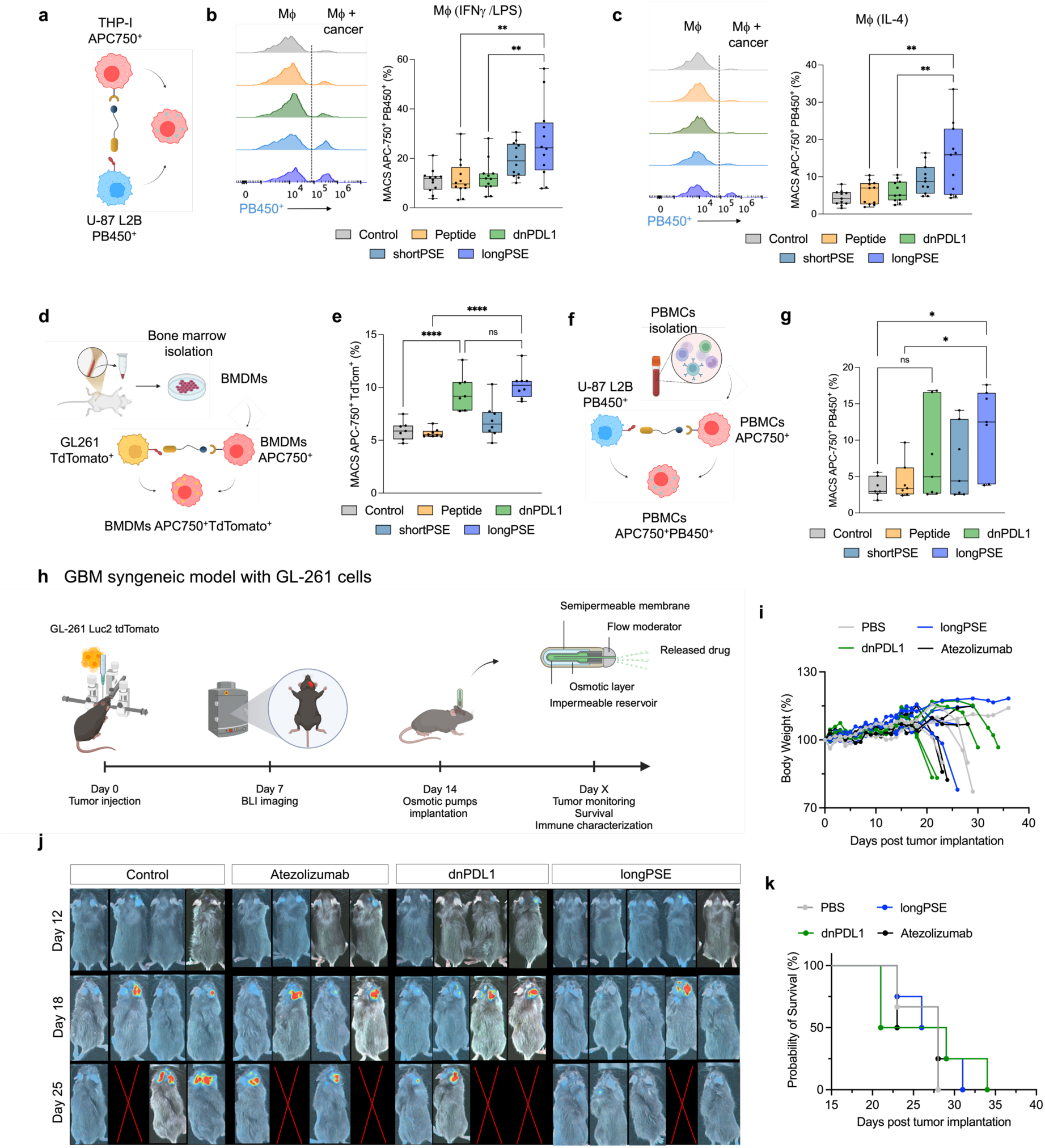
PSEs enhance phagocytosis of glioma cells in vitro and suppress tumour growth in glioblastoma syngeneic models. **a,** Schematic of phagocytosis assay of polarised THP-I Mϕ co-cultured with BFP-labelled U-87 GBM cells +/- PSEs. **b,** Bar graphs represent the phagocytic activity given by the amount of double-positive Mϕ/total amount of pro-inflammatory (stimulated with IFN-γ/LPS) Mϕ. FC analysis after treatment of polarised THP-I Mϕ/U-87 co-cultures (n=12). Histograms relative to PSE-treated cells show an increase in double-positive APC750^+^ PB450^+^ THP-I Mϕ (Mϕ + Cancer). (Linear peptide vs longPSE: p=0.0013; dnPDL1 vs longPSE: p=0.0022). **c,** Bar graphs represent the phagocytosis activity given by the amount of double-positive Mϕ /total amount of anti-inflammatory (stimulated with IL-4) Mϕ (n=11). FC analysis after treatment of polarised THP-I Mϕ /U-87 co-cultures. Histograms relative to PSE-treated cells show an increase in double-positive APC750^+^ PB450^+^ THP-I Mϕ (Mϕ + Cancer). (Linear peptide vs longPSE: p=0.0011; dnPDL1 vs longPSE: p=0.0018). **d,** Schematic of phagocytosis assay of isolated murine BMDMs (stained with Cell Trace far-red dye) in co-culture (1:1) with mouse tdTomato-labelled GL-261 cells +/- PSEs. **e,** Bar graphs representing the phagocytosis activity given by the amount of double-positive BMDMs/total amount of BMDMs (n=8. FC analysis after treatment of BMDM/U-87 co-cultures. Y axis represents percentage of double-positive APC750^+^ tdTom^+^ THP-I cells. (Control vs dnPDL1: p<0.0001; Linear peptide vs longPSE: p<0.0001). **f,** Schematic of phagocytosis assay of isolated human primary Mϕ (stained with CellTrace Far-Red proliferation kit) in co-culture (1:1) with BFP-labelled U-87 cells +/- PSEs. **g,** Bar graphs representing the phagocytosis activity given by the amount of double-positive primary Mϕ/total amount of primary Mϕ. FC analysis after treatment of primary Mϕ /U-87 co-cultures (n=6). y-Axis represents percentage of double positive THP-I APC750^+^ PB450^+^. (Control vs longPSE: p=0.0196; Linear peptide vs longPSE: p=0.0466). Statistical analysis was performed using ordinary one-way ANOVAs, followed by Tukey’s multiple comparison test to compare treatment groups. In every instance the asterisk p>0.05 = ns, *p<0.05, **p<0.01, ***p<0.001, ****p<0.0001. **h,** Experimental design of the in vivo study in a syngeneic GBM model. 5×10^5^ GL-261 L2tdTomato cells were inoculated stereotactically into C57BL/6J mice on d 0 and monitored by BLI for tumour engraftment, then different treatments (longPSE, dnPDL1, peptide or PBS) (n=4-5 per group) were administered i.t. on d14 via osmotic pumps. Mice were sacrificed when they reached humane endpoint. **i,** body weight changes during treatment with the longPSE, dnPDL1, Atezolizumab and vehicle (PBS). The body weight of each group of mice measured everyday. **j,** BLI of the mice in the different treatment groups at d 12, 18, 25. Tumour, growth is delayed in the group treated with longPSE. **k,** Overall survival curve shows no benefit for the PSE treatment group that displays no tumour growth until the endpoint. Illustrations were created with Biorender.

Across the different co-culture systems, longPSE consistently induced the highest phagocytosis activity, therefore we tested its efficacy in a GBM syngeneic model using the cell line GL-261 L2tdTomato orthotopically injected in the pre-frontal cortex of C57BL/6 mice. Tumour cell growth was followed by serial bioluminescent imaging (BLI) (**Fig. 3h**). Local administration of brain tumour treatments is limited by the amount of drug that can be safely administered at once without causing a symptomatic increase in intracranial pressure. A single application provides a rapid peak in drug concentration but very short duration of drug exposure. To circumvent this, we implanted osmotic minipumps for continuous i.t. delivery at d14 post glioma cell inoculation (0.25 μl/h). Four treatment groups were selected: longPSE, dnPDL1, the PD-L1 inhibitor Atezolizumab and PBS. During the days following treatment, all the cohorts suffered from steep tumour growth and reduction in body weight (**Fig. 4i-j**), apart from the longPSE treatment arm that displayed a stationary tumour situation. We did not observe significant overall survival benefit (**Fig. 4k**), although the key difference was the absence of tumour in the longPSE group, while in all other groups the mice were sacrificed due to the tumour burden.

**Figure 4.**
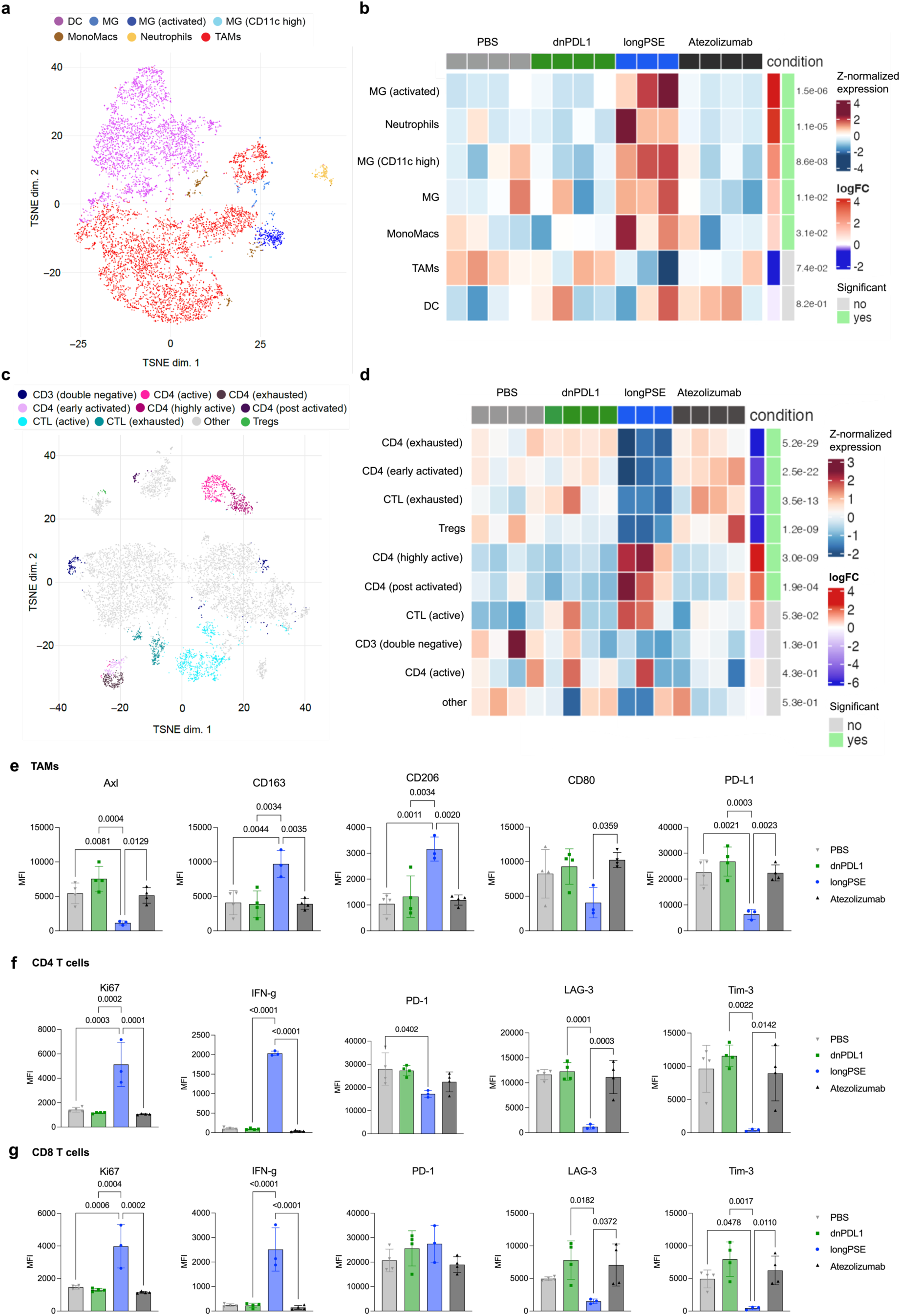
Immune characterization of tumour-bearing hemispheres reveals T cell activation and downregulation of immunosuppressive TAMs in mice treated with PSEs. **a,** t-distributed stochastic neighbour embedding (TSNE) plots depicting 7 main myeloid populations after initial merging and manual annotation from 12 populations originating from unbiased clustering. **b,** Box plots representing myeloid cell population abundance per therapeutic condition according to the clusters identified. **c,** t-distributed stochastic neighbor embedding (TSNE) plots depicting 10 main lymphoid populations after initial merging and manual annotation from 20 populations originating from unbiased clustering. **d,** Box plots representing lymphoid cell population abundance per therapeutic condition according to the clusters identified. **e,** TAMs expression marker analysis by flow cytometry. TAMs were gated as single, live, CD45+, CD11b+, F4/80+, CD49d+, PR2Y12-cells. **f,** CD4 T cells expression marker analysis by flow cytometry. CD4 T cells were gated as single, live, CD45+, CD3+, CD4+ cells. **g,** CD8 T cells expression marker analysis by flow cytometry. CD8 T cells were gated as single, live, CD45+, CD3+, CD8+ cells.

### Characterization of the brain tumour microenvironment highlights key phenotypic differences in mice treated with PSEs

To pinpoint the PSE-related immune-modulating effects in the complex GBM-TME, we examined the tumour-bearing hemisphere per each condition (**Fig. 4**). Single-cell suspensions were analysed by flow cytometry (FC) using two panels for the myeloid and lymphoid compartments (**Table S1-2**). Unsupervised cluster analysis was performed on single-cell FC data, and the protein marker expression was used for cell type annotation (**Fig. 4a-c**). The myeloid compartment of longPSE-treated brains displayed a relative increase in microglia (MG), transitory monocytes and neutrophils, while TAMs were less abundant (**Fig. 4b, Extended data figure 3b-c**). Differential expression analysis highlighted important differences between PSE-treated and the other groups, such as downregulation of PD-L1 and TAMs shifting towards a microglia-like phenotype (**Fig. 4e** and **Extended data figure 3c,e**).

Furthermore, we observed important differences between groups in the lymphoid compartment (**Fig. 4c-d**). The longPSE-treated brain tumours displayed highly activated and proliferating CD4 and CD8 T cells (**Fig. 4d**), indicating a state of expansion and proliferation in response to antigen stimulation. Moreover, the downregulation of inhibitory surface markers such as LAG-3 and TIM-3 pointed towards a functional, non-exhausted state (**Fig. 4f-g**). The high effector function driven by IFNγ was likely involved in the anti-tumour response and only occurred in the PSE treatment, suggesting the display of the phagocytosis-promoting peptide profoundly impacted immune activation. However, the overall activation of CD4 and CD8 T cells accompanied by IFNγ increase might be the cause of adverse-related immune events and premature death in the PSE-treated mice, which were devoid of detectable tumours at endpoint. We did not observe systemic toxicity by analysing the contralateral hemisphere and hematological parameters, which did not differ across groups (**Extended data figure 6b-7**).

In summary, the continuous PD-L1 blockade, in the context of high IFNγ release from T cells, led to a more effective anti-tumour response without the typical compensatory immune evasion arising from PD-L1 upregulation. This combination could thereby sustain a prolonged activated T-cell response.

### PSEs mediate the internalization of membrane PD-L1 and extracellular targets in Mϕ

Our studies confirmed that PD-L1 was widely expressed in the TME by tumour cells and immunosuppressive myeloid cells^59–62^ (**Fig. 5**). To follow-up on the pronounced downregulation of PD-L1 upon longPSE treatment, we used PD-L1-expressing TAMs to study the effect of longPSE at cellular level. We performed cell assays to track PSE and PD-L1 (**Fig. 5a and Extended data figure 4a-b**) by incubation of PD-L1-expresing Mϕ with FITC-PSE and observed that the PSE (**Fig. 5a**, green channel) and PD-L1 (**Fig. 5a**, red channel) co-localised with the lysosomes (**Fig. 5a**, yellow channel). In addition, the treatment led to a strong decrease in surface PD-L1 staining (**Extended data figure 4a-c**). Overall, we observed a 68% decrease in PD-L1 surface staining after treatment with longPSE and 47% decrease with the dnPDL1 binder compared to control (**Fig. 5b**). Imagestream analysis revealed that longPSE leads to a 2-fold higher PD-L1 co-localisation in the lysosomes compared to dnPDL1, while the control (media only) did not show PD-L1 internalization but a strong staining of surface PD-L1 (**Extended data figure 4a-c**). On the other hand, the same treatment applied to tumour cells did not lead to internalization of the probe FITC-PSE (**Fig. 5c**), although PD-L1 blockade of the longPSE on tumor cells had the same inhibition curve as the unconjugated binder dnPDL1 and was comparable to Atezolizumab (Atezolizumab IC_50_ = 1 nM; longPSE IC_50_= 3nM, **Fig. 5d**). The results described above suggest that target internalization was Mϕ-specific and that the bifunctional molecule favoured the endolysosomal pathway.^63^

**Figure 5.**
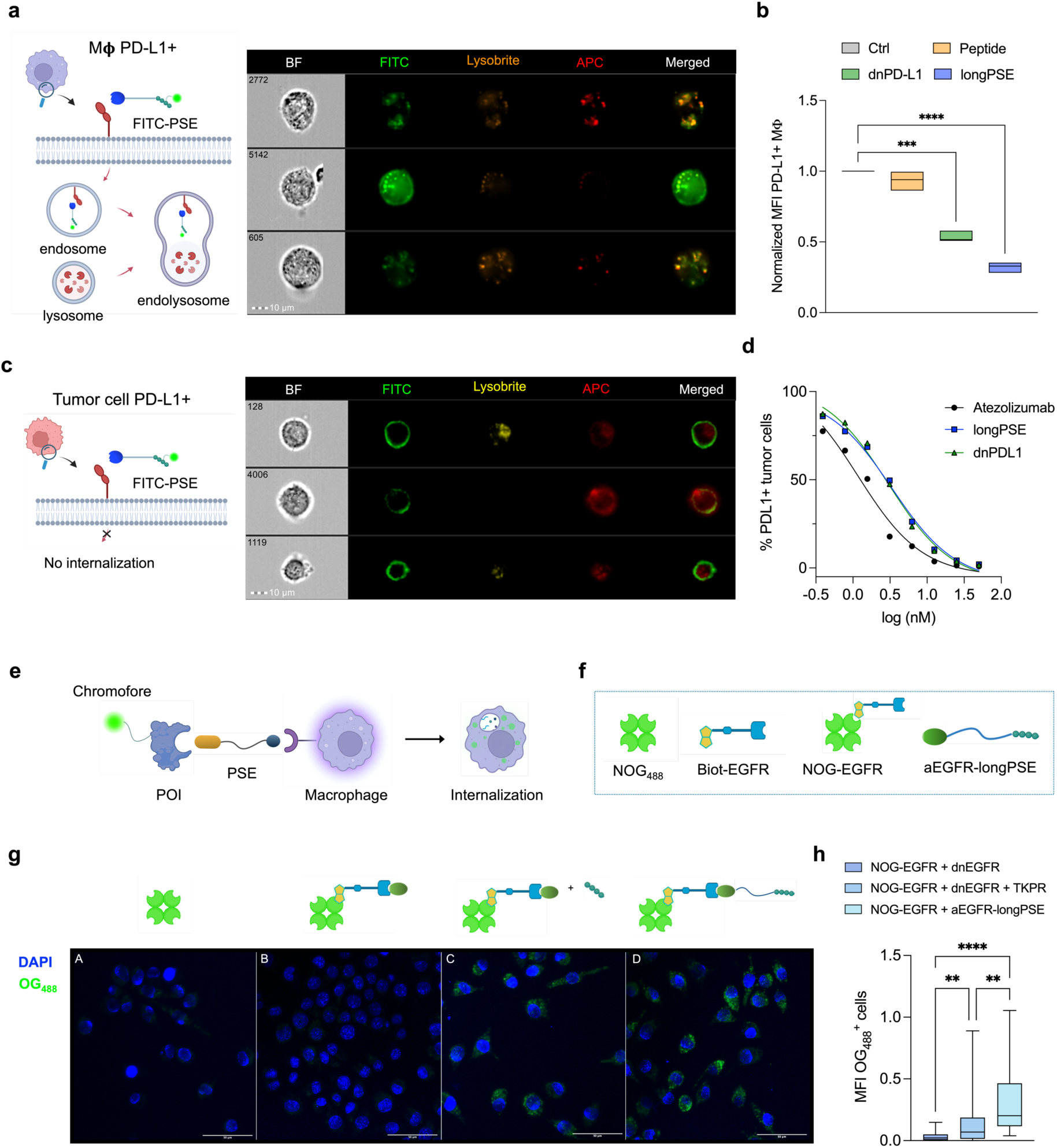
PSEs enhance the uptake of extracellular or membrane protein target via the endolysosomal pathway. **a,** Single-cell image analysis and co-localization studies of PD-L1 expressing THP-I cells treated with FITC-PSE. Single cell imaging analysis was performed 4h after incubation. Cells were pre-treated with Lysobrite Red (lysosomal marker). Cells treated with FITC-PSE were stained with anti PD-L1-APC after incubation. From left to right, the following channels were used: BF to localize the cells (grey channel), FITC to localize the probe FITC-PSE (green channel), Lysobrite to localize the lysosomes (yellow channel), APC to localize the PD-L1 (red channel). Merged comprises green, yellow and red channels. **b,** FC analysis of THP-I Mϕ. PD-L1 staining was measured by the mean fluorescence intensity of APC+ Mϕ and normalised to the fluorescence of untreated APC+ cells (control). (Control vs linear peptide: ns, Control vs dnPDL1: p=0.0003, Control vs longPSE: p<0.0001, n=3). **c,** Single-cell image analysis and co-localization studies of GL-261 tumor cells treated with FITC-PSE. Single cell imaging analysis was performed 4h after incubation. Cells were pre-treated with Lysobrite Red (lysosomal marker). Cells treated with FITC-PSE were stained with anti PD-L1-APC after incubation. From left to right, the following channels were used: BF to localize the cells (grey channel), FITC to localize the probe FITC-PSE (green channel), Lysobrite to localize the lysosomes (yellow channel), APC to localize the PD-L1 (red channel). Merged comprises green, yellow and red channels. **d,** PD-L1 blocking assay with the PD-L1 expressing tumor cell line MDA-MB-231. Range used: 0.39-50 nM by serial dilutions. A fixed concentration of antibody was used (aPDL1-APC, 1:500 dilution, 133 ng/ml). Non-linear fit log (inhibitor) vs. response (three parameters) on GraphPad Prism v10; R^2^ values: 0.9786 (Atezolizumab); 0.9926 (dnPDL1); 0.9898 (longPSE). IC_50_ values: 1.17 (Atezolizumab); 3.343 (dnPDL1); 2.987 (longPSE). **e,** Schematic representation for the enhanced uptake of extracellular targets PSE-mediated. A protein of interest (POI) carrying a chromophore is bound by the PSE, which triggers uptake of the POI in Mϕ determined by increased fluorescence intensity of the specific chromophore. **f,** Illustration of the molecules used for the uptake of a soluble extracellular protein. **g,** Confocal analysis of the enhanced uptake. Panels A-D display RAW 264.7 Mϕ treated with A) NOG_488_; B) NOG_488_-EGFR complex + dnEGFR; C) NOG_488_-EGFR complex + dnEGFR + tuftsin; D) NOG_488_-EGFR complex + aEGFR-longPSE. Images contain the merged channels of i) blue channel - nuclei stained with DAPI; ii) green channel – NOG_488_-EGFR complex. **h,** Quantitative fluorescence measurements of the intracellular uptake of the complex NOG_488_-EGFR. (NOG-EGFR + dnEGFR vs aEGFR-longPSE: p<0.0001, NOG-EGFR + dnEGFR vs NOG-EGFR + dnEGFR + TKPR: p=0.0089, NOG-EGFR + dnEGFR + TKPR vs aEGFR-longPSE: p=0.0014). The average MFI was calculated by processing the images on QuPath Statistical analysis was performed on Prism v10 using ordinary one-way ANOVAs, followed by Tukey’s multiple comparison test to compare treatment groups. In every instance the asterisk * indicates a p<0.05, ** indicates p<0.01, *** indicates p<0.001, and **** indicates p<0.0001. Illustrations created with Biorender.

Mϕ lysates showed a decrease in PD-L1 after longPSE treatment at different timepoints, however absolute PD-L1 degradation remained difficult to prove due to the Mϕ transient PD-L1 expression influenced by their polarization state. In fact, Mϕ treated with longPSE after 24h secreted up to 7-fold higher amount of TNF-α in the supernantant (**Extended data figure 4d**). We assessed in vivo TNF-α secretion with a Mϕ reporter double transgenic xenograft Tg(mpeg1:mCherry-F; tnfa:eGFP-F) allowing the identification of Mϕ populations (M_1_-like = mpeg^+^ / tnf^+^; M_2_-like = mpeg^+^ / tnf^-^, **Extended data figure 8**). The PSEs shifted TAMs towards a pro-inflammatory phenotype (mpeg^+^ / tnf^+^, **Extended data figure 8b-c**). Since the zebrafish embryos have not developed adaptive system at the time of the treatment (2 dpf), the treatment response is directly related to the innate immunity and demonstrates the PSE-induced Mϕ reprogramming.

We further explored whether the PSE-enhanced internalization could be applied to a an extracellular target (**Fig. 5e**). We used a de novo EGFR binder (dnEGFR) to assemble the bifunctional EGFR-based PSEs (**Extended Figure 5a-d**). Biot-EGFR was pre-incubated with NOG_488_ to yield the complex NOG_488_-EGFR which was incubated with aEGFR-longPSE or controls in the presence of Mϕ (**Fig. 5g**). aEGFR-longPSE induced the strongest internalization of EGFR (panel D, **Fig. 5g-h**). This data suggests that the PSE-targeted antigens are internalised by Mϕ. In recent years, several modalities have been developed to direct a protein of interest to cellular compartments such as lysosomes dedicated to protein degradation.^64–75^ This PSE-mediated mechanism might favor the efficacy of PSEs, as it is believed that Ab-mediated internalization of immune checkpoints can further potentiate the immune response.^76^

In summary, the PSEs induced T-cell’s antitumour activity by PD-L1 blockade and enhanced the phagocytic activity of Mϕ in the TME. In addition, the PSEs promoted PD-L1 internalization on PD-L1^+^ Mϕ and enhanced the internalization of an extracellular target.

## Discussion

The complex networks between the immune system and the TME, combined with tumour heterogeneity, pose major hurdles in the development of therapeutic modalities able to eradicate the tumour with high specificity and efficacy. We developed a modular strategy using an exemplary anti PD-L1 binder conjugated to the phagocytosis-promoting peptide tuftsin. Through acoustic force spectroscopy, we demonstrated the PSEs increased the binding strength between effector and target cells. The PSEs enhanced phagocytosis of colorectal tumour cells and glioma cells compared to their respective single modules or control media.

In a preclinical syngeneic model of CRC, treatment with the bifunctional longPSE showed delayed tumour progression and improved overall survival, highlighting its superior efficacy compared to its respective modules and CD47 blockade. In a more challenging syngeneic model of GBM, the PSE did not improve the overall survival, although we observed striking effects on the immune compartment and delayed tumour progression. Differential expression analysis highlighted key phenotypic changes, primarily PD-L1 downregulation in TAMs and IFNγ release from activated T cells.

Compared to checkpoint blockade therapies such as anti-PD-1/PD-L1 antibodies, PSEs provide an advantage by incorporating both immune checkpoint inhibition and immune-stimulatory elements, potentially overcoming resistance mechanisms that arise from monotherapy approaches. Similarly, while CAR-T cell therapy has demonstrated efficacy in hematologic malignancies, its success in solid tumors remains limited due to issues such as poor TME infiltration and immunosuppressive signaling. PSEs, through their modular design, can counteract these barriers by engaging innate immune cells (e.g., macrophages) and enhancing their tumoricidal activity, complementing adaptive immune responses. Furthermore, the modularity of PSEs allows for rapid adaptation to different tumor types by tuning their functional domains, making them a versatile tool in the evolving immunotherapy landscape.

While the results herein described are very promising, there are limitations that will require further investigations to unravel the full mechanism of action. In the GBM model, while the potent and continuous immune response effectively clears the tumour, it might also be the cause of adverse-related events that need to be addressed in translational settings. Furthermore, the role of the linkers and why longPSE performed better than others is not fully understood. We anticipate the choice of the linker plays a pivotal role towards TME targeting and the molecular mechanism of action. Screening of larger PSE libraries are needed to further validate these hypotheses, directing future studies towards fine-tuning the phenotypic changes occurring in the TME.

Overall, these findings provide a first development towards the therapeutic application of ENPHASYS in cancer immunotherapy. The phagocytosis-enhancing properties of tuftsin combined with immune checkpoint inhibition is a promising approach that potentiates the anti-tumour immune response. Alternatively, PSEs offer a distinct approach that can be exploited to internalize extracellular or membrane targets. As we explore these avenues in cancer immunotherapy, we seek to expand this technology towards new targets to modulate tumour immunity and restore phagocytes to their native function.

## Extended Data Figure

**Extended data figure 1.**
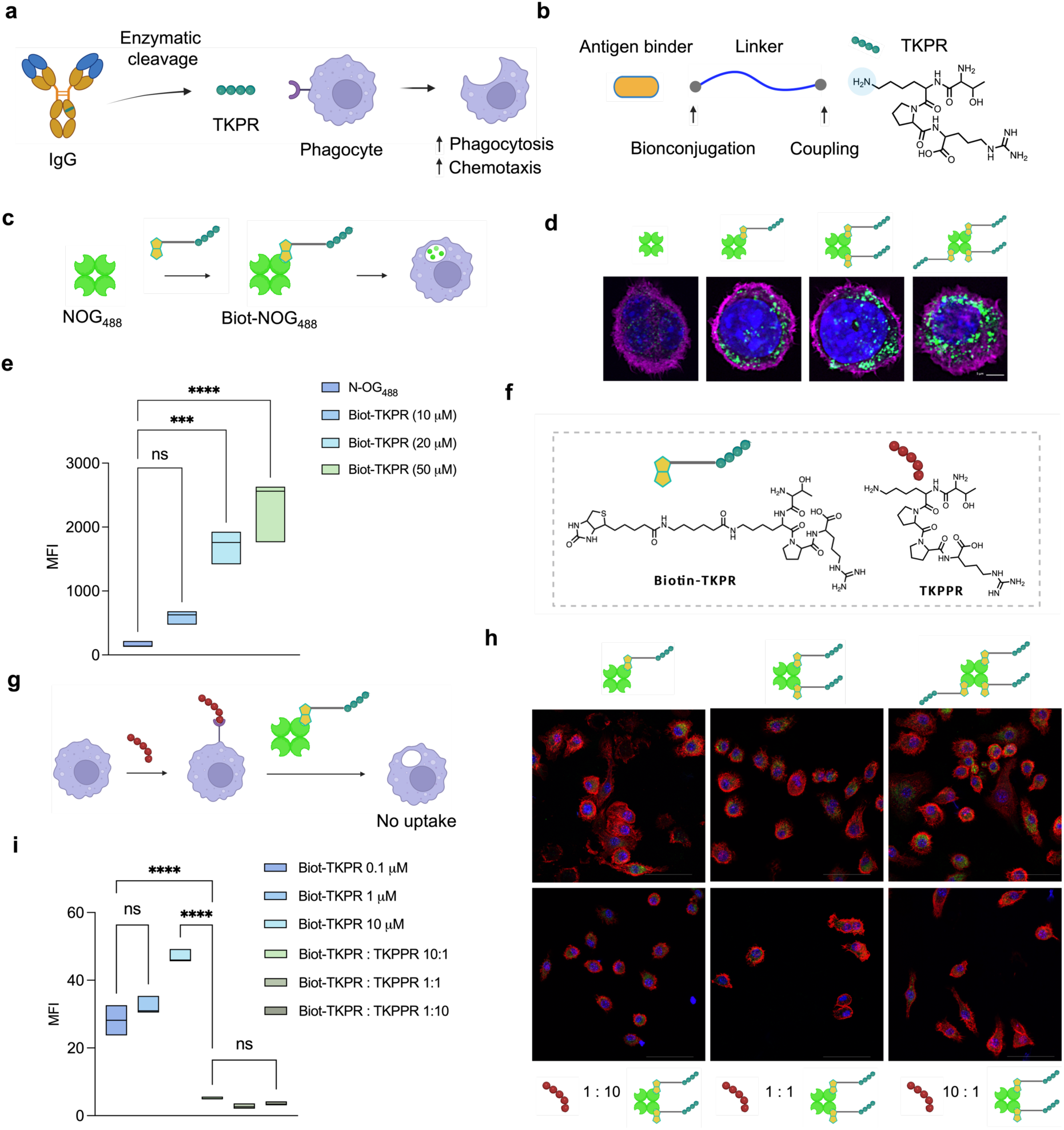
Tuftsin-mediated uptake assays with RAW 264.7 Mϕ. **a,** Tuftsin (TKPR) is a natural peptide derived from the catabolism of IgG. **b,** Assembling of the PSE through the lysine side chain of tuftsin. **c,** Schematic representation of enhanced uptake of a fluorescent protein. The fluorescent protein Neutravidin Oregon Green_488_ (NOG_488_) is complexed to the biotinylated promoter Biot-TKPR leading to enhanced uptake in RAW 264.7 Mϕ. **d,** Microscopy images of fluorescence uptake. From left to right: NOG_488_, NOG_488_-Biot-TKPR (10, 20, 50 μM). Scale bar: 5 μm. **e,** Relative quantitative analysis of the mean fluorescence intensity of intracellular NOG_488_. (NOG_488_ vs Biot-TKPR 10 μM: p=0.3016; NOG_488_ vs Biot-TKPR 20 μM: p=0.0007; NOG_488_ vs Biot-TKPR 50 μM: p<0.0001). **f,** Molecular structure of Biot-TKPR and TKPPR. **g,** Schematic representation of uptake inhibition in the presence of the tuftsin antagonist TKPPR. Pre-incubation with TKPPR inhibits the uptake of Biot-NOG_488_. **h,** Imaging of Biot-NOG_488_ uptake in the absence (upper panels) or presence (lower panels) of TKPPR. Scale bar: 50 μm. **i,** quantitative analysis of mean fluorescence intensity relative to h. (Biot-TKPR 0.1 μM vs Biot-TKPR: TKPPR 10:1 p<0.0001; Biot-TKPR 10 μM vs Biot-TKPR: TKPPR 10:1 p<0.0001). The intracellular fluorescence intensity was calculated by processing the images on QuPath. Statistical analysis was performed using ordinary one-way ANOVAs, followed by Tukey’s multiple comparison test to compare treatment groups. In every instance *p<0.05, **p<0.01, ***p<0.001, and ****p<0.0001. Illustrations created with Biorender.

**Extended data figure 2.**
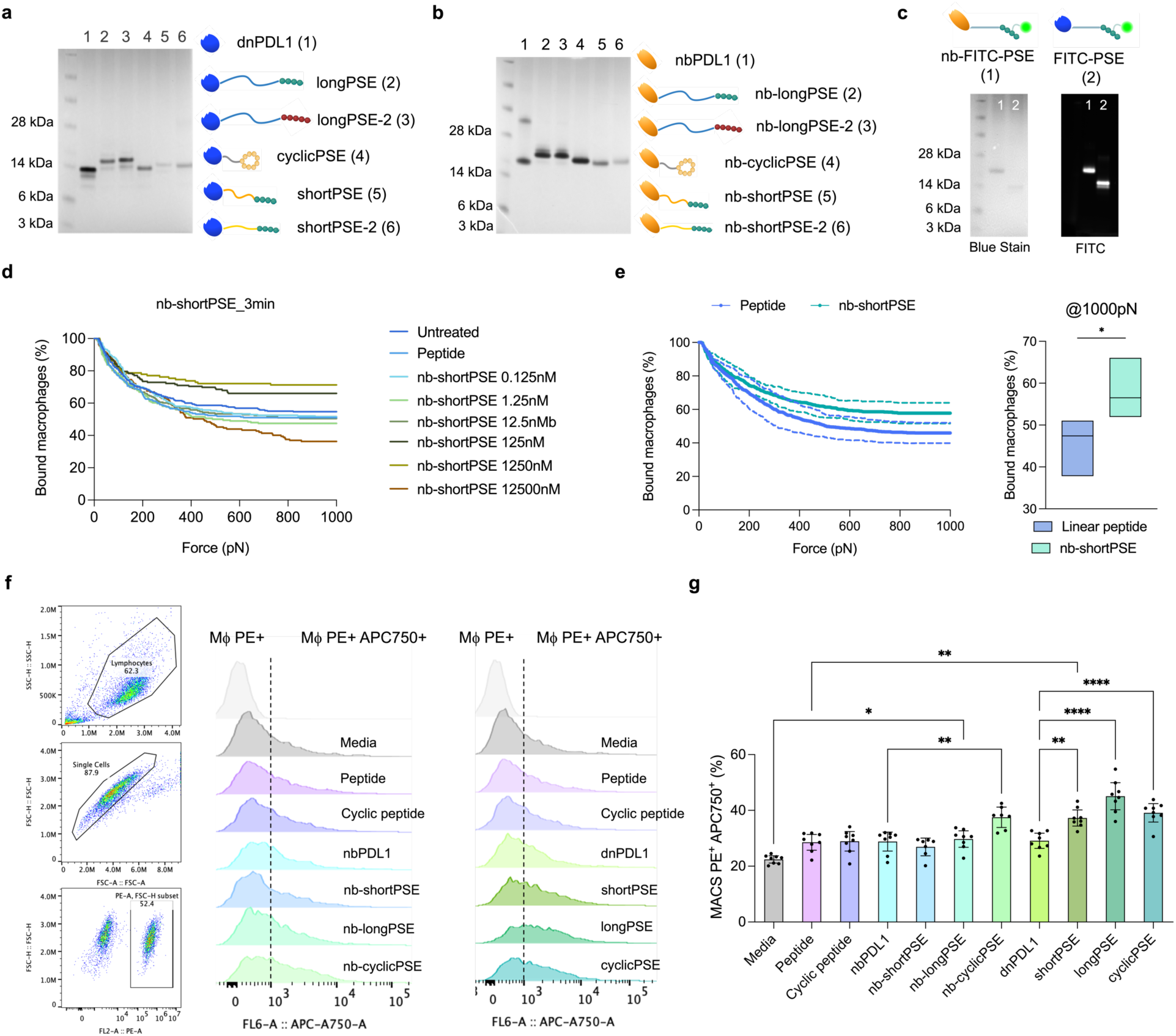
Synthesis and in vitro characterization of a nanobody-based PSEs targeting PD-L1. **a,** SDS-PAGE of nanobody-based (nb) PSEs targeting PD-L1. **b,** SDS-PAGE of de novo PSEs targeting PD-L1. **c,** SDS-PAGE and FITC blot analysis of PSE fluorescent probes nb-FITC-PSE and FITC-PSE. Flow cytometry analysis of PD-L1 binding assays. PSE probes nb-FITC-PSE and FITC-PSE were incubated with MC-38 cells untreated or pre-treated for 24h with IFNγ (20 ng/ml). **d,** histograms show the fluorescence intensity relative to the FITC channel. **e,** Avidity assays with nb-shortPSE. Analysis of bound Mϕ after 3 min incubation with the PSE in co-culture before application of acoustic force (0 to 1000 pN). PSE nb-shortPSE shows increased avidity compared to the linear peptide (n=4). **f,** Flow cytometry analysis of double-positive Mϕ (PE^+^ APC750^+^) after treatment with PSEs in co-culture with MC-38 cells. Histograms show the increase in APC750^+^ Mϕ treated with the PSEs. **g,** Phagocytic index of pre-gated Macs PE^+^. Mϕ and tumor cells were co-culture for 4h in the presence of PSEs or relative controls. (Media vs nb-longPSE, p=0.0151; Peptide vs shortPSE, p=0.0013; ndPDL1 vs nb-cyclicPSE, p=0.0022; dnPDL1 vs shortPSE, p=0.0034; dnPDL1 vs longPSE, p<0.0001; dnPDL1 vs cyclicPSE, p<0.0001). Illustrations created with Biorender.

**Extended data figure 3.**
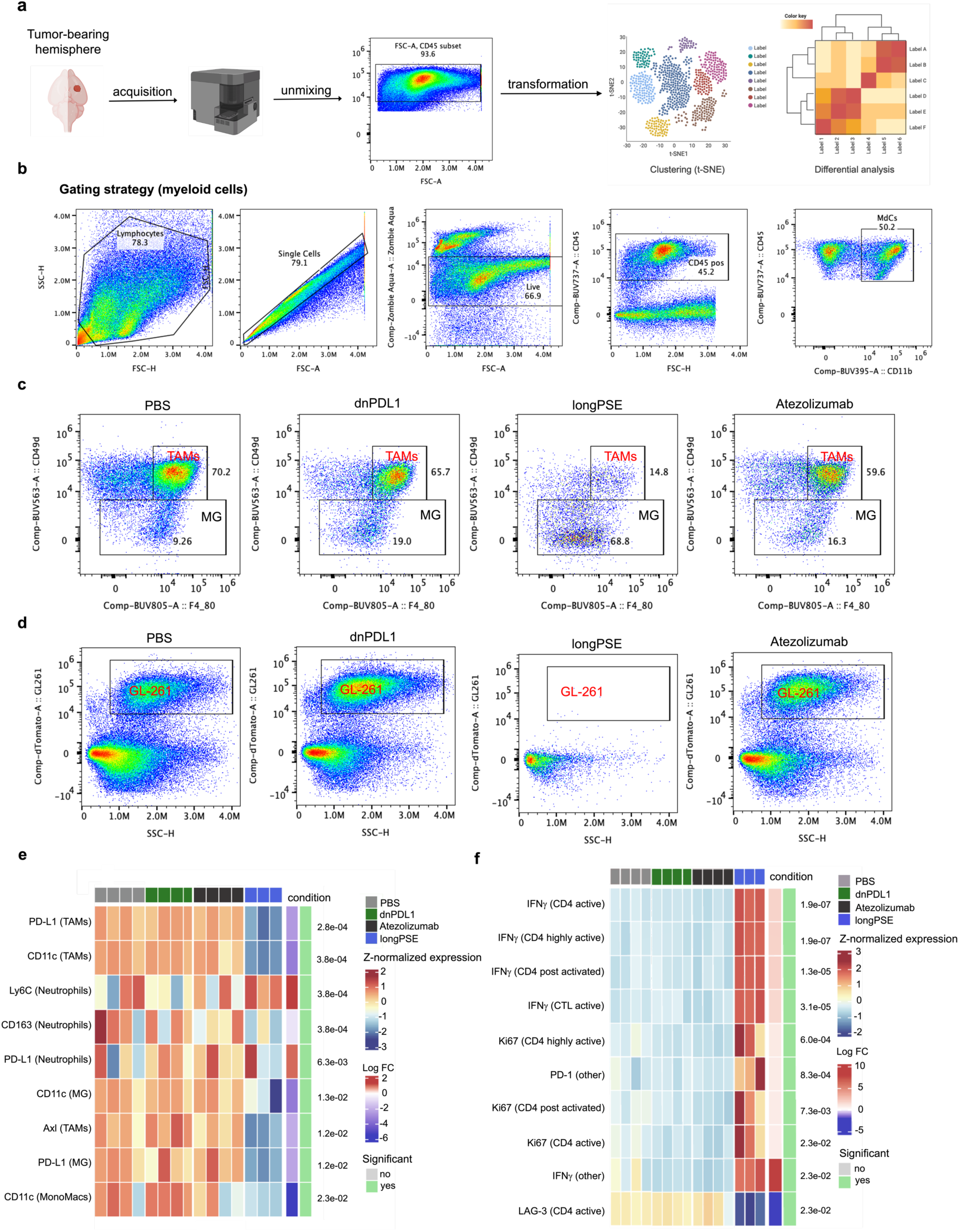
Myeloid and lymphoid populations of PSE-treated brain tumors display the phenotypic changes of potent antitumour response. **a,** Flowchart visualizing the characterization of the brain tumour-bearing hemisphere. After acquisition and unmixing, pre-gated live CD45+ events were processed via FlowSOM analysis and differential data analysis. **b,** Gating strategy for the myeloid cell panel. Myeloid-derived cells were gated as live/single/CD45+Cd11b+ cells. **c,** Representative diagrams of TAMs and MG abundance in the tumour-bearing hemisphere for each condition. TAMs are gated as live/single/CD45+Cd11b+CD49d+ cells and MG live/single/CD45+Cd11b+CD49d-cells. **d,** GL-261 (tdTomato+) cancer cells abundance in the tumour-bearing hemisphere after each treatment. **e,** Heatmap of z-scores for aggregated marker expression per individual sample highlighting the significant changes in myeloid markers expression in PBS (n=4), longPSE (n=3), dnPDL1 (n=4), Atezolizumab (n=4). Reference condition: PBS, contrast condition: PSE. **f,** Heatmap of z-scores for aggregated state marker expression (removing lineage markers: CD3, CD4, CD8, CD45) per individual sample highlighting the significant changes in lymphoid state markers expression (removing lineage markers: CD45, P2ry12, Ly6G, F4/80, CD11b, CD49d) in PBS (n=4), longPSE (n=3), dnPDL1 (n=4), Atezolizumab (n=4). Reference condition: PBS, contrast condition: longPSE. The statistical analyses were conducted in R using the diffcyt package, specifically with the diffcyt-DS-limma method, to identify differentially abundant cell populations between the reference and contrast conditions. Adjusted p-values (using FDR) were computed to control for multiple testing, with significant results filtered at a false discovery rate of 0.05. For visualization, a log fold change threshold of 0.5 was applied in the heatmap to highlight meaningful expression changes in specific markers by condition. Illustrations created with Biorender.

**Extended data figure 4.**
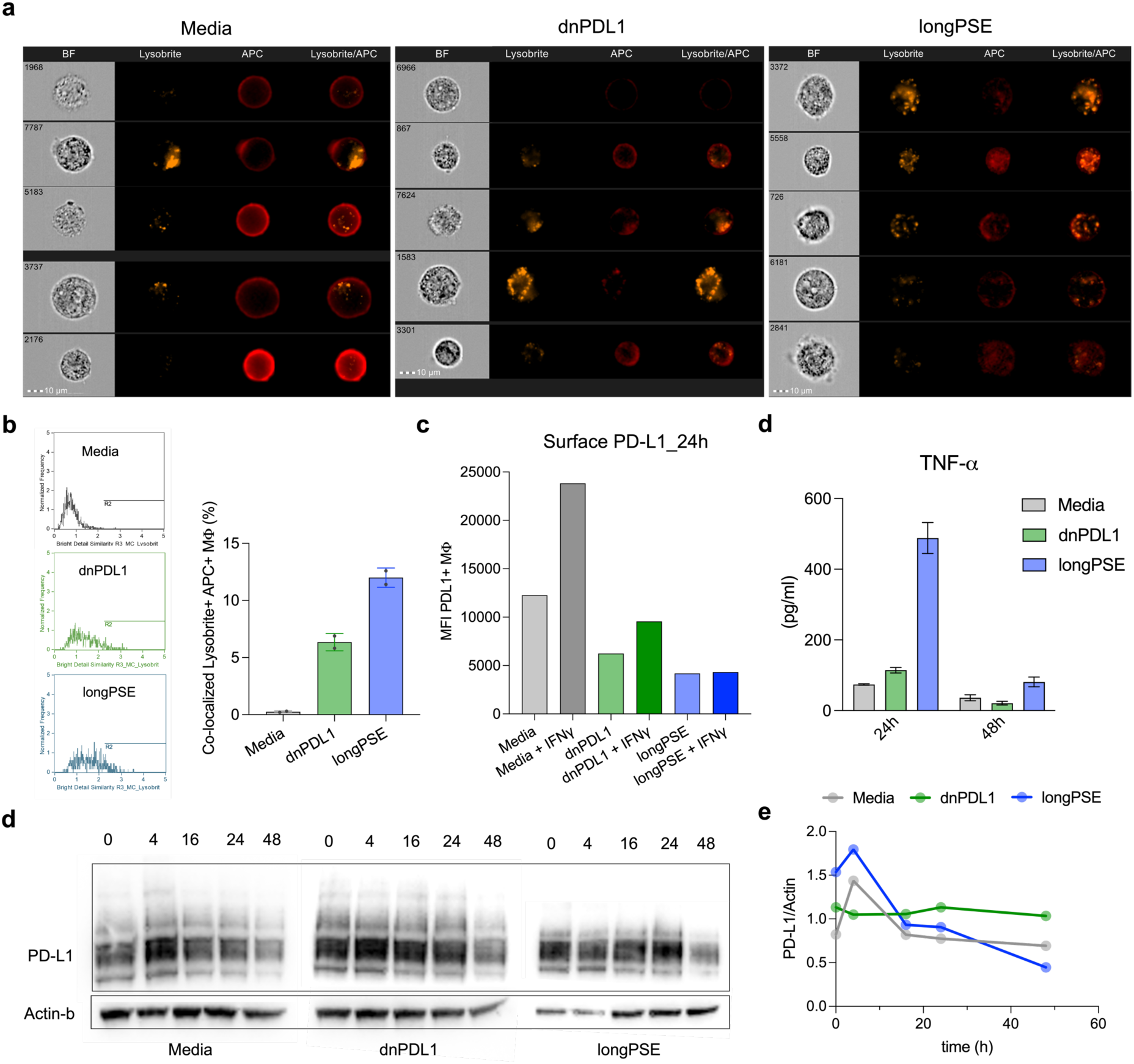
PD-L1 analysis of THP-I Mϕ treated at 4h and 24h suggest target internalization and degradation PSE-mediated. **a,** Imagestream analysis of THP-I Mϕ after 4h incubation with: Media only, dnPDL1, longPSE. Concentration of compounds is 100 nM. After treatment, THP-I Mϕ were stained with aPDL1-APC. Cells were pre-gated as live single cells. From left to right, each image displays channels for Brightfield (BF), Lysobrite Red (lysosome stain), APC (aPD-L1-APC), Lysobrite/APC (merged). Scale bar: 10 μm. Image analysis was performed on IDEAS 6.2 software. Scatter plot and statistics available on Supplementary Information, Fig. S31. **b,** Co-localization analysis of THP-I Mϕ treated with: Media, dnPDL1, longPSE. Co-localization was measured by the bright detail similarity score of double positive Mϕ (Lysobrite+ APC+). Quantitative analysis was performed on IDEAS 6.2 software. **c,** Flow cytometry analysis of PD-L1 on THP-I Mϕ at 24h treatment with Media, dnPDL1 and longPSE in the presence or absence of IFNγ (20 ng/ml). **d,** Analysis of Mϕ culture supernatants at 24-48h after treatments with Media, dnPDL1, longPSE. The amount of TNF-α was determined by ELISA MAX Standard Set Human TNF-α according to the manufacturer protocol (Biolegend, 430201, Lot. No: B393898). **e,** WB analysis of PD-L1 in THP-I Mϕ lysates at timepoints 0-48h: Media only, dnPDL1, longPSE. **f,** quantitative analysis of PD-L1 in THP-I Mϕ lysates after treatment at different timepoints (0-48h) with Media, dnPDL1 and longPSE. PD-L1 was quantified as the ratio of mean gray values between PD-L1 and Actinϕ blots. Analysis was performed on ImageJ software. Statistical analysis was performed using one-way ANOVAs with Tukey’s multiple comparison and Mann Whitney unpaired t test to compare treatment groups.

**Extended data figure 5.**
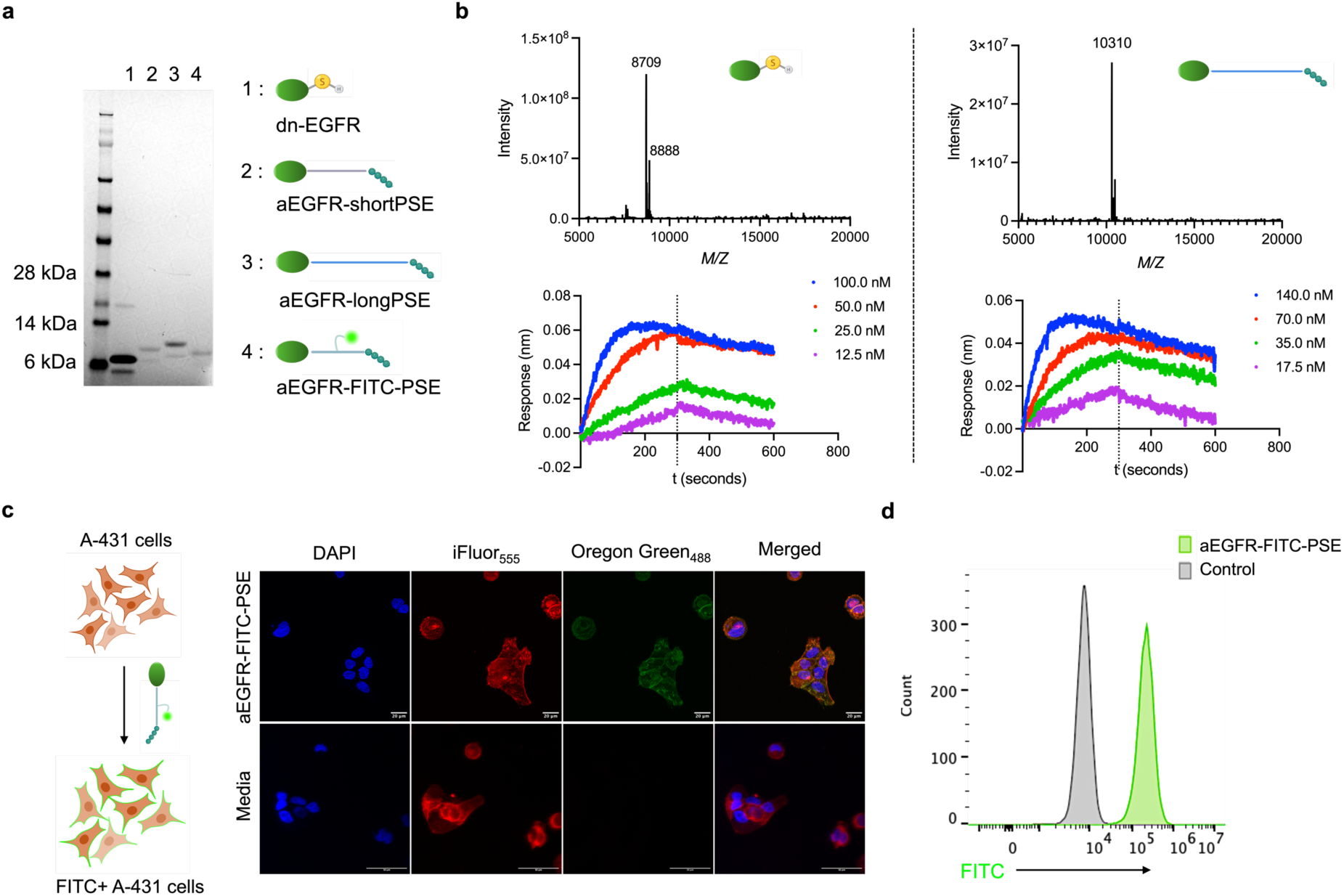
Synthesis and characterization of EGFR-targeting PSEs. **a,** SDS-PAGE of anti-EGFR de novo binder (dnEGFR)-based PSEs targeting PD-L1. **b,** Analysis of dnEGFR and aEGFR-longPSE by LC-MS and biolayer interferometry analysis. Dissociation constants are calculated by non-linear regression analysis using the global analysis model for association kinetics within the Prismv10 software. (dnEGFR: K_D_ = 4.2 nM; aEGFR-longPSE: K_D_ = 5.0 nM). **c,** Confocal microscopy analysis of A-431 cells labelled with the fluorescent probe aEGFR-FITC-PSE after 2h incubation at 37 °C. Panels from left to right: blue channel – nuclei stained with DAPI; red channel – cytoskeleton stained with Phalloidin iFLuor_555_; OG_488_ channel – EGFR stained with aEGFR-FITC-PSE. Scale bar: 20 µm (top panels), 50 µm (bottom panels). **d,** Flow cytometry analysis of EGFR-expressing A-431 cells after 30 min incubation on ice with aEGFR-FITC-PSE.

**Extended data figure 6.**
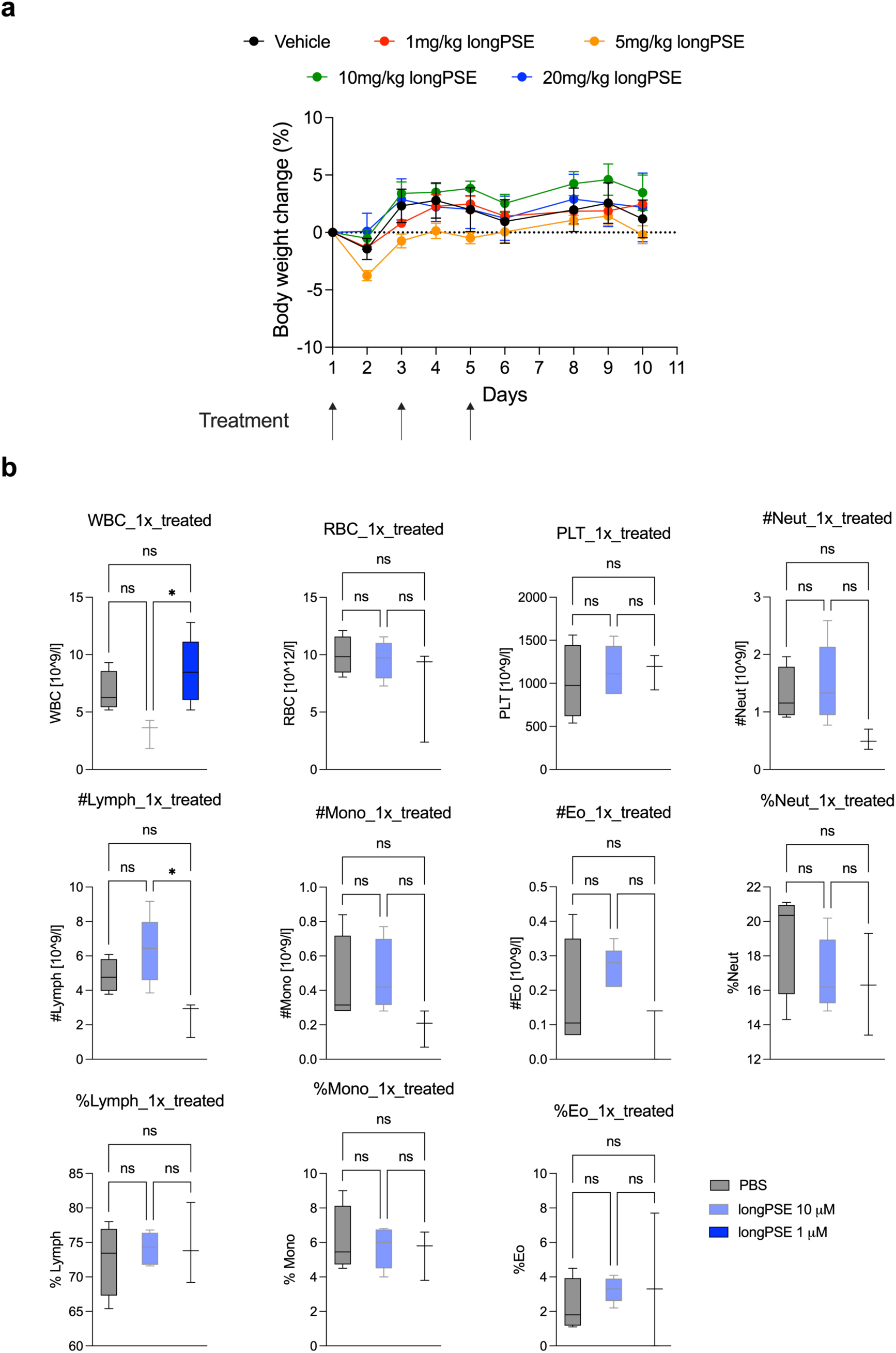
Toxicity studies with syngeneic models of CRC and GBM. **a**, Maximum tolerated dose studies with 1 / 5 / 10 or 20 mg/kg of longPSE treatment (4 mice/group), every other day, in a total of 3 doses, by intraperitoneal injection. Body weight of mice was monitored throughout the study and mice were sacrificed 5 days after last treatment. **b**, Toxicity studies of GL261-L2t tumor-bearing C57BL/6 mice treated with a single injection of longPSE (1 μM/10 μM) or PBS (4 μl, 1 μl/min). Blood samples were collected 24 hours after treatment by tail vein blood draw. Whole blood samples were immediately analysed using a haemocytometer (XN-1000V, Sysmex). Parameters used to detect haematological toxicity include WBC, RBC, MCV, MCH, and PLT. Statistical analysis was performed in Prism (version 10). Ordinary one-way ANOVAs were performed with Tukey’s multiple comparison’s test to compare treatment groups. In every instance the asterisk * indicates a p<0.05, ** indicates p<0.01, *** indicates p<0.001, and **** indicates p<0.0001.

**Extended data figure 7.**
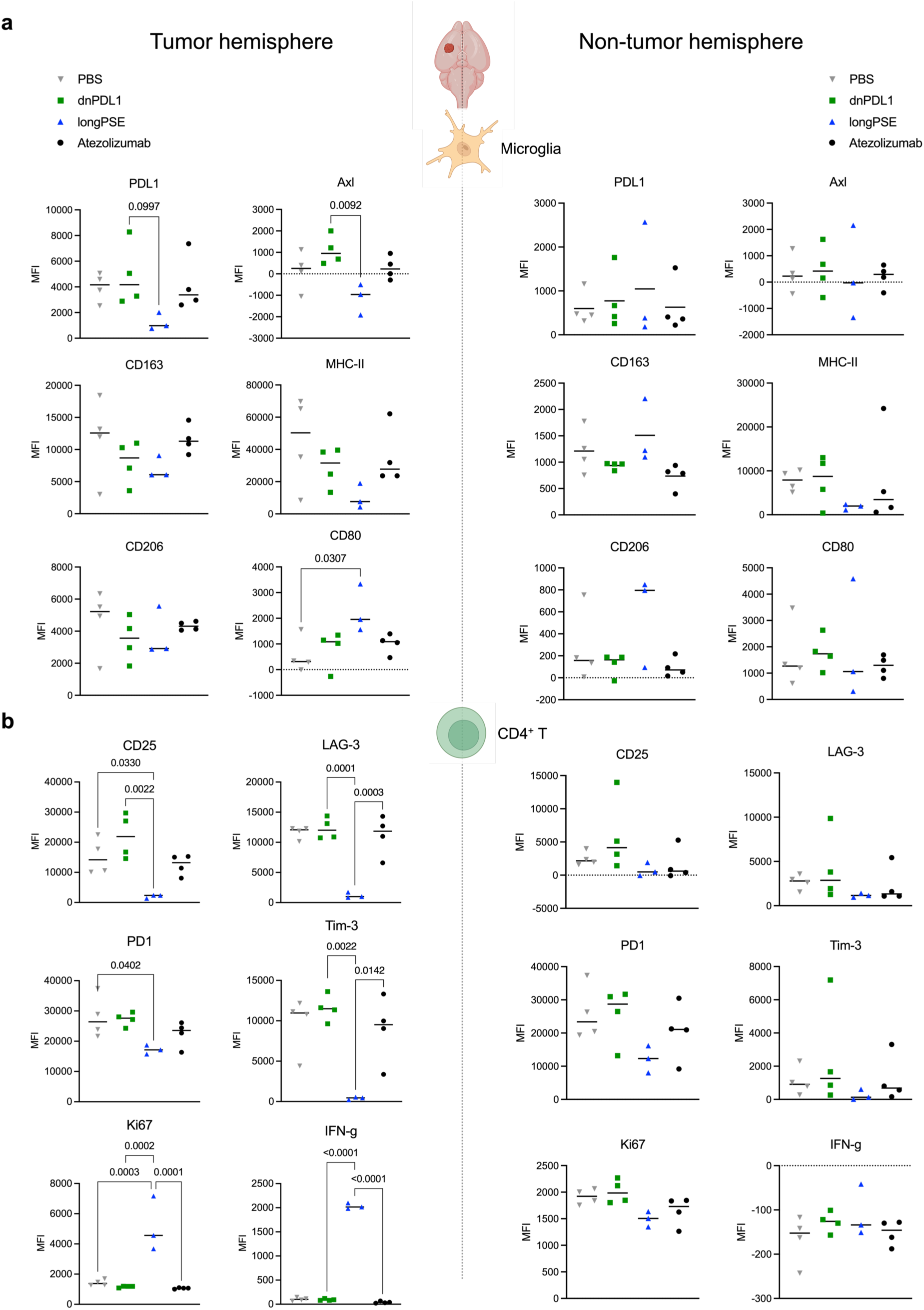
Analysis of the contralateral hemisphere in mice treated with PBS, dnPDL1, longPSE and Atezolizumab do not reveal significant phenotypic differences. **a,** Analysis of microglia markers expression in tumor-bearing and healthy hemisphere. **b,** Analysis of CD4+ T cells markers expression in tumor-bearing and healthy hemisphere. Statistical analysis was performed in Prism (version 10). Ordinary one-way ANOVAs were performed with Tukey’s multiple comparison’s test to compare treatment groups. In every instance the asterisk * indicates a p<0.05, ** indicates p<0.01, *** indicates p<0.001, and **** indicates p<0.0001.

**Extended data figure 8.**
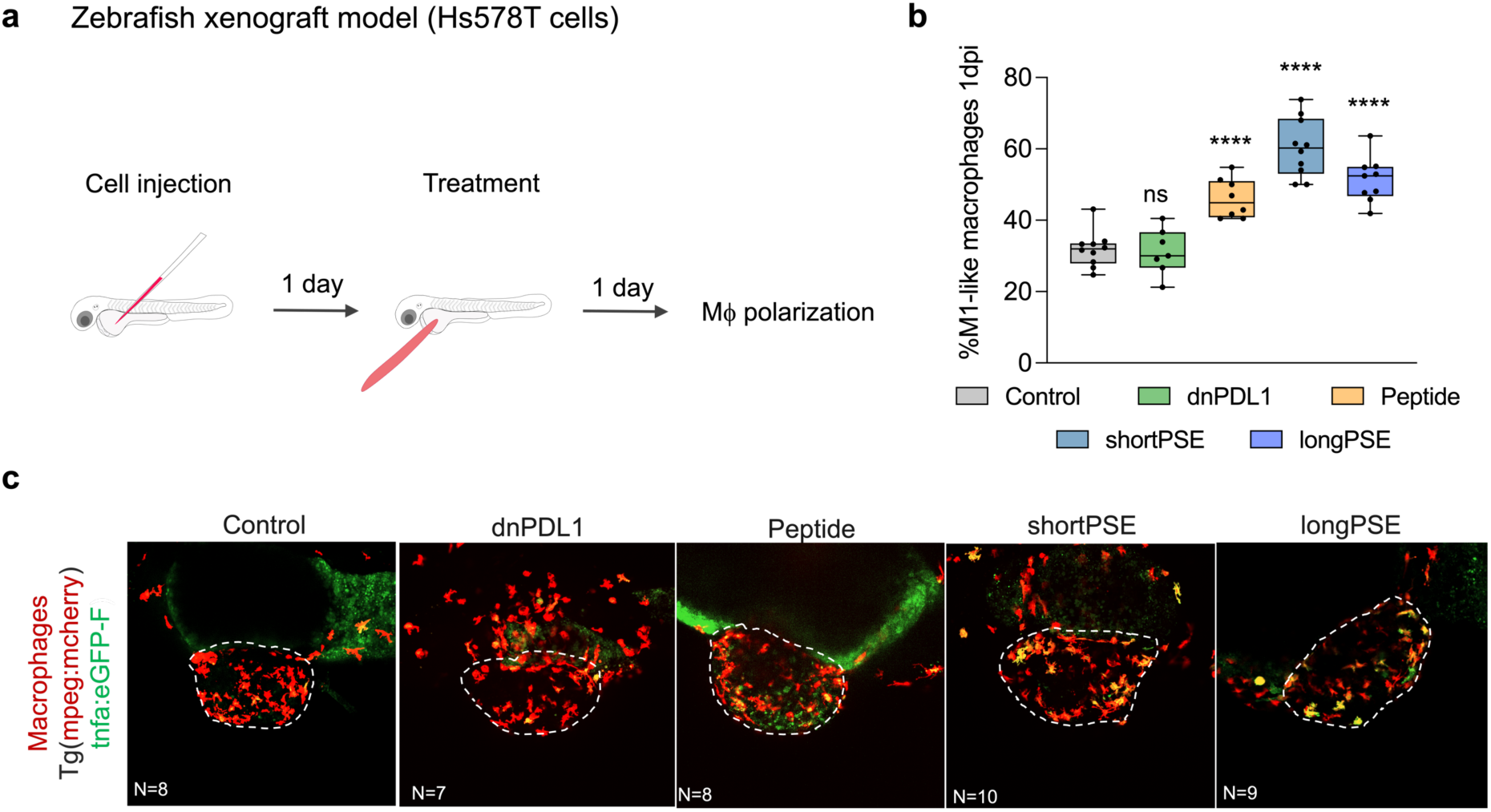
PSEs induce pro-inflammatory TAMs in zebrafish xenograft model. **a,** Experimental design: Hs578T TNBC cells were injected into the PVS of 2dpf zebrafish embryos and resuspended in different PSE compounds. Each compound was added to the cell suspension prior to injection. At 1dpi, xenografts were distributed into 5 treatment groups: Control, dnPDL1, Peptide, shortPSE and longPSE. The compounds were added to the water of the fish. **b,** percentage of M1-like macrophages at 1dpi (Control vs linear peptide: p<0.0001, Control vs shortPSE: p<0.0001, Control vs longPSE: p<0.0001). Graphs are presented as average ± SEM. Results are from two independent experiments. Statistical analysis was performed using an unpaired two-sided Mann–Whitney test. Statistical results: ns > 0.05, *P ≤ 0.05, **P ≤ 0.01, ***P ≤ 0.001, and ****P ≤ 0.0001. **c,** Representative confocal images of Mϕ in Hs578T xenografts in Tg(mpeg1:mcherryF, tnfa:eGFP-F) at 1dpi. Mϕ activation was determined by expression of TNFα by reporter Mϕ co-expressing GFP. Scale bars: 50 μm. Dashed lines outline the tumours. All images are anterior to the left, posterior to right, dorsal up, and ventral down. The number of xenografts analysed is indicated in the representative images and each dot in the graphs represents one zebrafish xenograft.

## Acknowledgements

V.S. thanks the Swiss National Science Foundation (P500PN_202711, P5R5PB_217904), the Cambridge Academy of Therapeutic Sciences, Novartis Medical-Biological Foundation (24B120) and Swiss Life Jubiläumsstiftung for fundings. This work was supported by a Swiss National Science Foundation Professorial Fellowship (PP00P3_176974); the ProPatient Forschungsstiftung, University Hospital Basel (Annemarie Karrasch Award 2019, additional research support); Swiss Cancer Research Grant (KFS-4382-02-2018 and KFS 5789-02-2023) to G.H.; the Department of Surgery, University Hospital Basel, to G.H. and Xi’an Fengcheng Hospital, to G.J.L.B. The authors thank Dr. Marie-Françoise Ritz for helpful discussions for in vivo GMB experiments and for amendment of the animal licences relative to this work. F.M.M. thanks the Herchel Smith Fund. C.T. thanks Fundação para a Ciência e a Tecnologia Portugal (2022.08101.CEECIND).

## Author Contributions

V.S. and G.J.L.B conceived the study. V.S. designed, synthesised and characterised the PSEs, designed and performed biological assays, visualised and analysed the data, wrote the manuscript. C.T., C.L-A., A.R.C. performed in vivo studies and analysis with CRC model. S.F. performed in vivo studies with GBM model. D.H. and W.Y. produced the mini binders dnPDL1 and dnEGFR. F.M. and W.T.K. synthesised the PSEs in larger scale for the in vivo studies. S.H. developed the methodology for unsupervised clustering and differential expression analysis. I.S., D.S and R.R. designed, performed, and analysed avidity assays with the support of V.S. A.V. synthesised the peptide cTKPRG. M.C-C. produced the PD-L1 targeting nanobody nbPDL1 and helped with microscopy experiments. T.R. helped with the phagocytosis assays and flow cytometry analysis. D.K. designed the panels for immune characterization. F.G. performed PBMC isolation and primary monocyte extraction. M.F. and R.F designed and performed zebrafish experiments. G.H. co-supervised the project, allocated resources and funding, co-wrote the manuscript. G.J.L.B coordinated and supervised the project, allocated resources, co-wrote the manuscript. The manuscript was finalised with contributions from all authors.

## Competing Interests statement

V.S. and G.J.L.B. are inventors on a patent application that incorporates the discoveries described in this manuscript [United Kingdom (GB) Patent Application No: 2318879.0]. D.H., W.Y., and D.B. are inventors on a patent application for the de novo PDL1 binder. D.B. is an inventor on a patent application for the de novo EGFR binder. G.H. has equity in, and is a cofounder of Incephalo Inc. C.T., D.H., A.R.C., W.Y., D.B., and G.J.L.B. are cofounders and shareholders of Vesto Therapeutics Inc. All other authors declare no conflict of interests.

## Data availability

All data are available within the Article, Supplementary Information. Raw data are available upon request.

## Code availability

All code used for data analysis is referenced in the relevant Methods subsections. R scripts used in the study are adopted from our previous work and are available in a GitHub repository under https://doi.org/10.5281/zenodo.1381990.

## Notes

https://doi.org/10.5281/zenodo.1381990

