## Supplementary Information for "Bifunctional Phagocytic Synapse Enhancers for Cancer Immunotherapy"

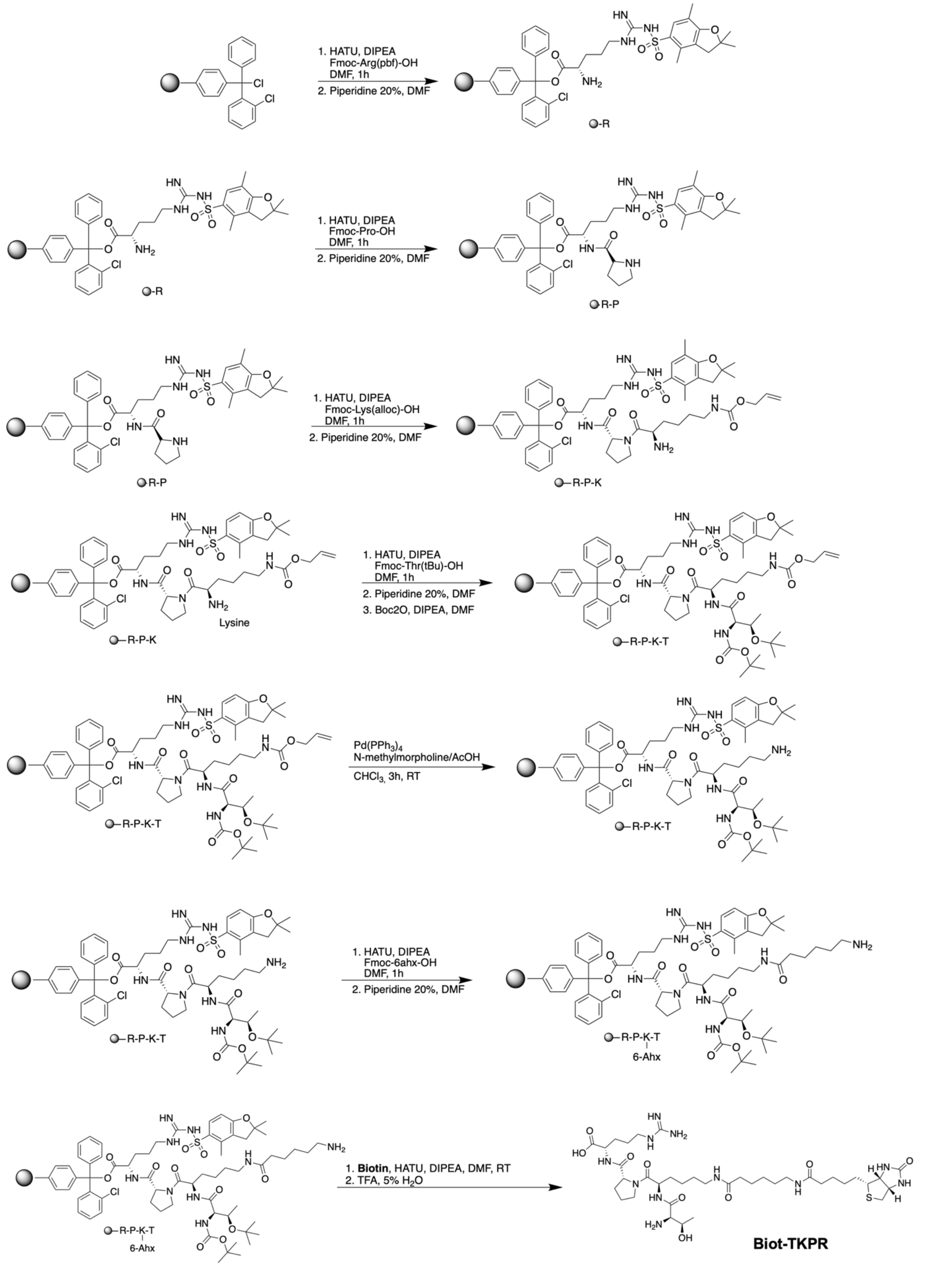

**Figure S1. Synthesis of Biot-TKPR by standard solid-phase peptide synthesis.**

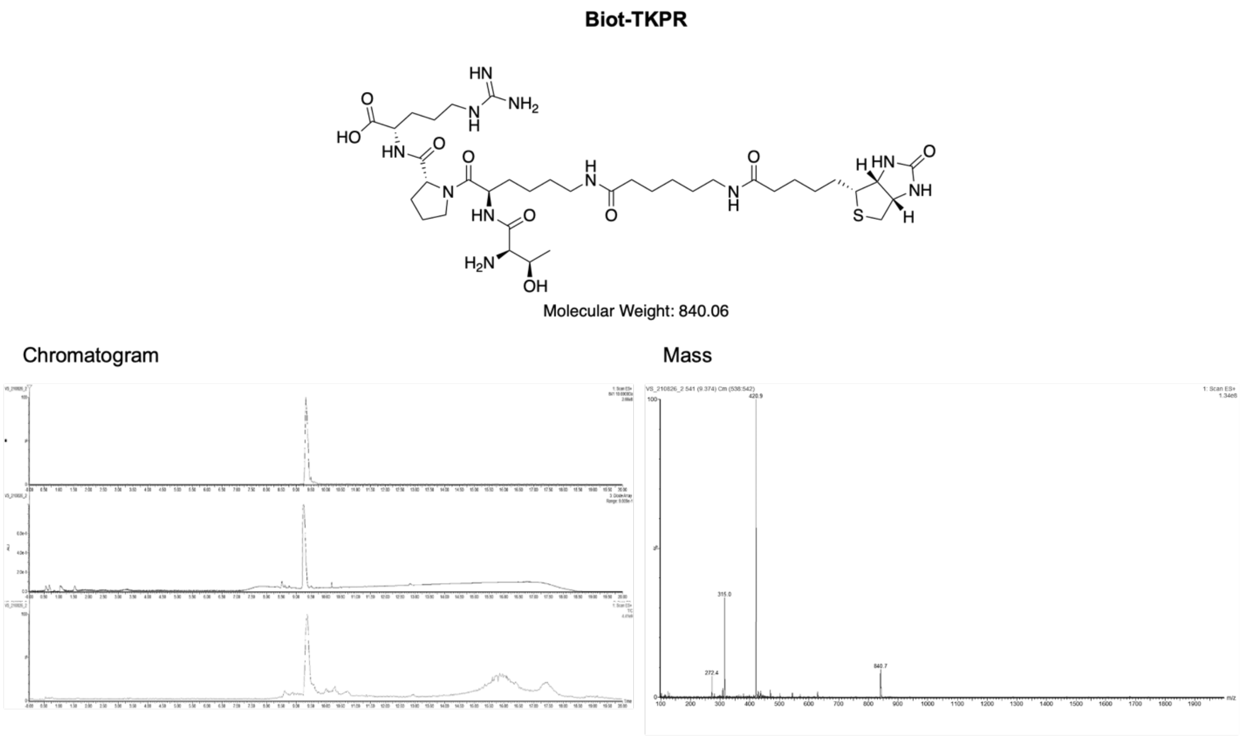

**Figure S2. Characterization of Biot-TKPR.**

**Maleimidocaproyl-cTKPRG**

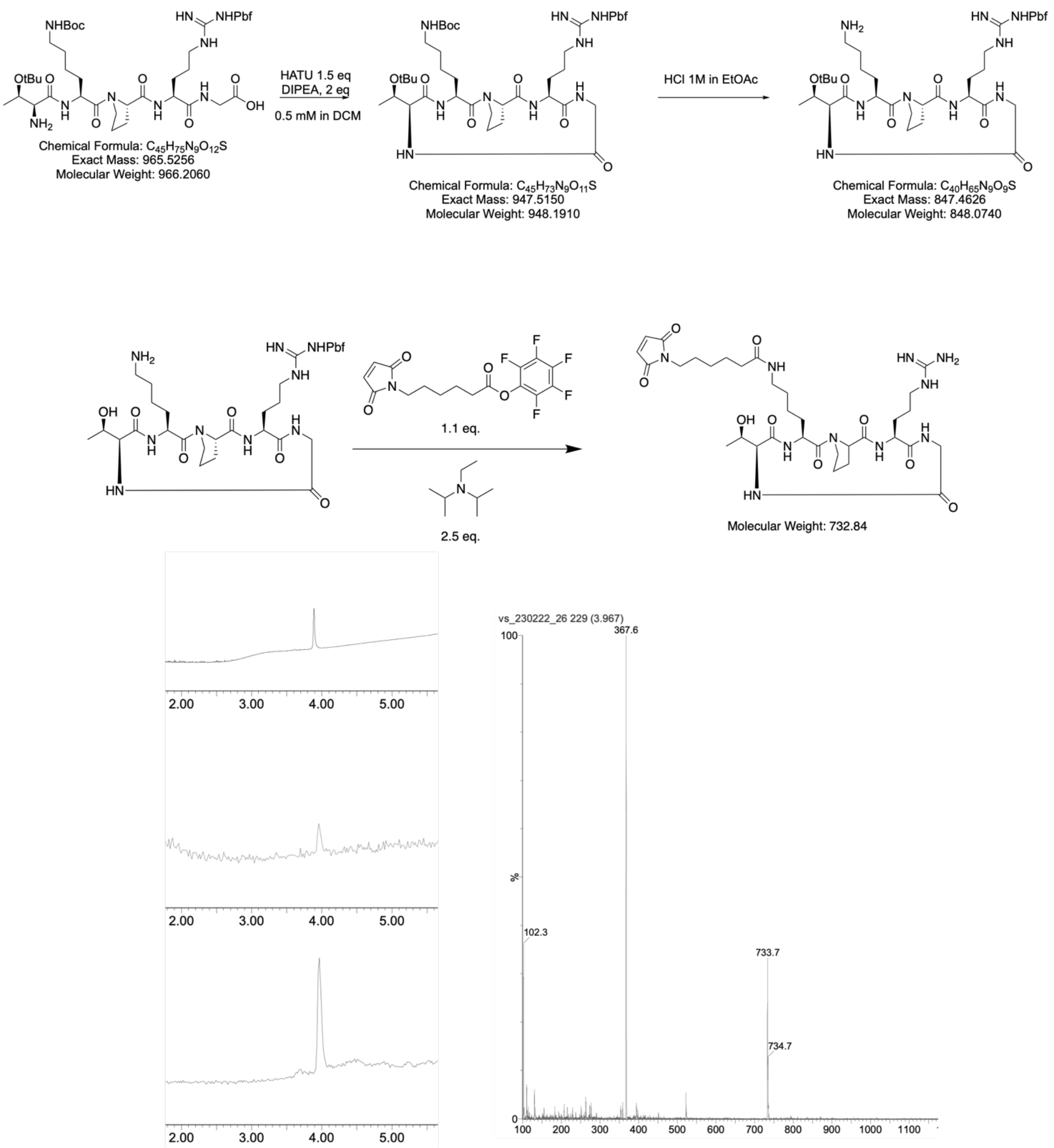

**Figure S3. Synthesis and characterization of the cyclic analogue cTKPRG and tethering to maleimide.**

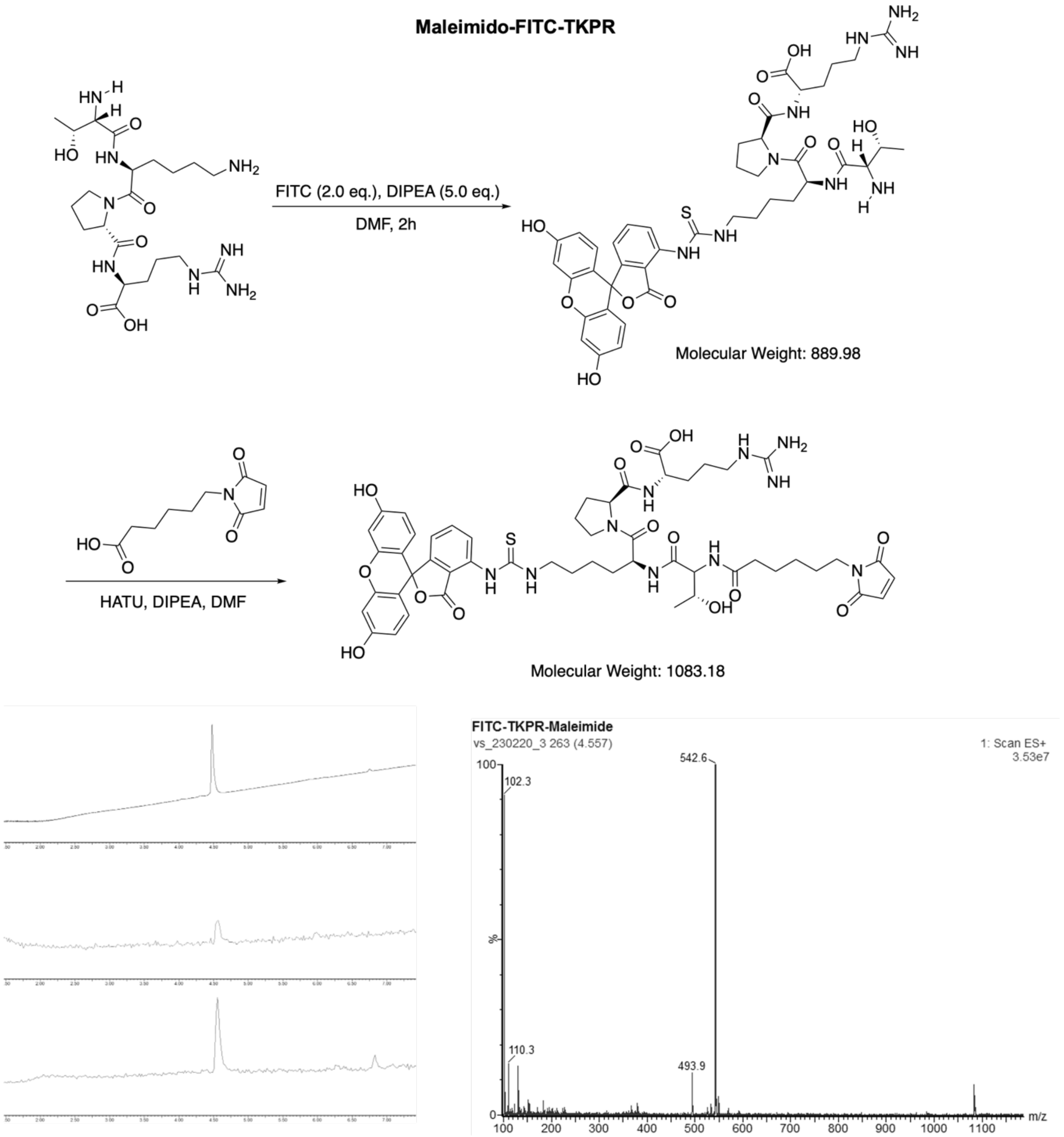

**Figure S4. Synthesis and characterization of the fluorescent probe precursor maleimido-FITC-TKPR.**

**Maleimido-shortPEG-TKPR**

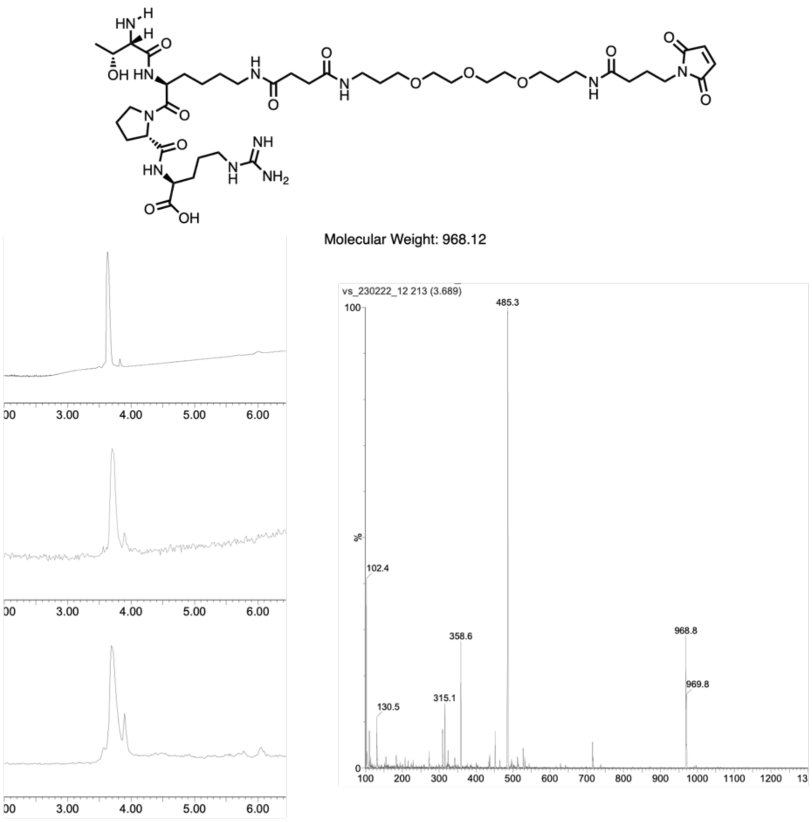

**Maleimido-longPEG-TKPR**

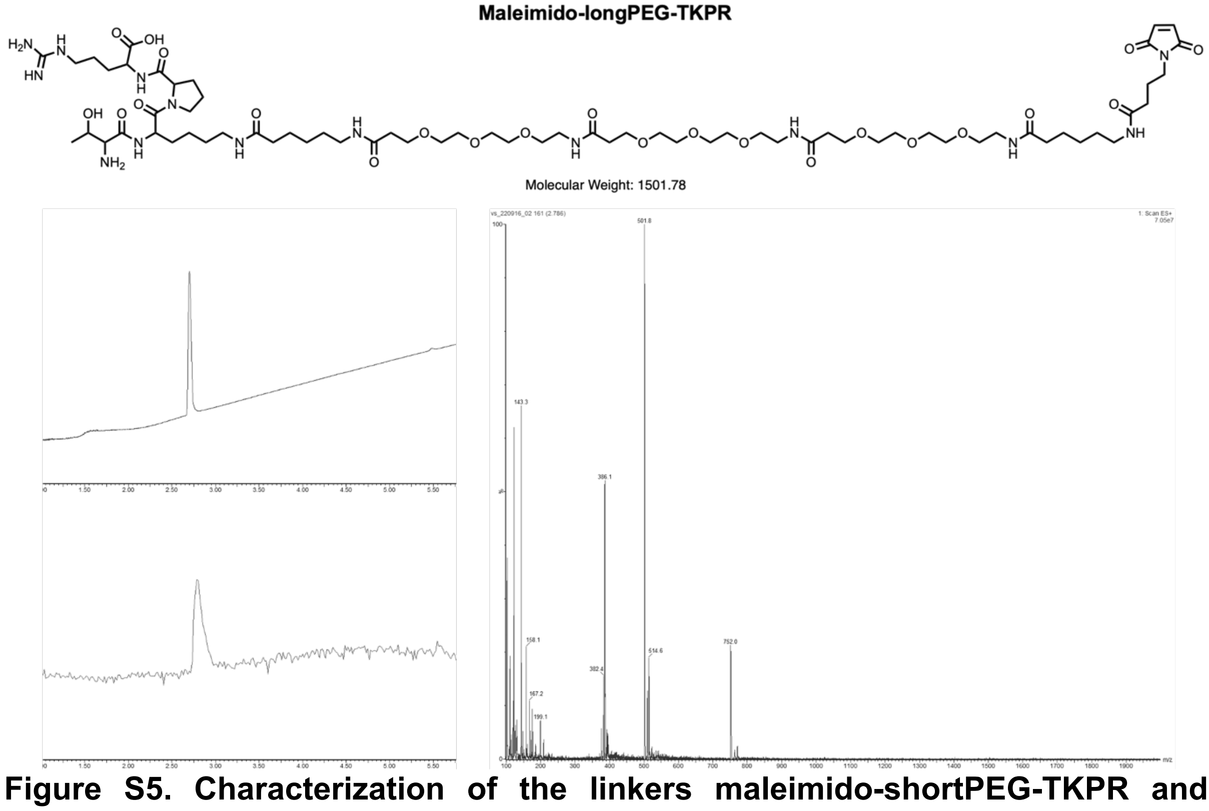

**Figure S5. Characterization of the linkers maleimido-shortPEG-TKPR and maleimido-longPEG-TKPR.**

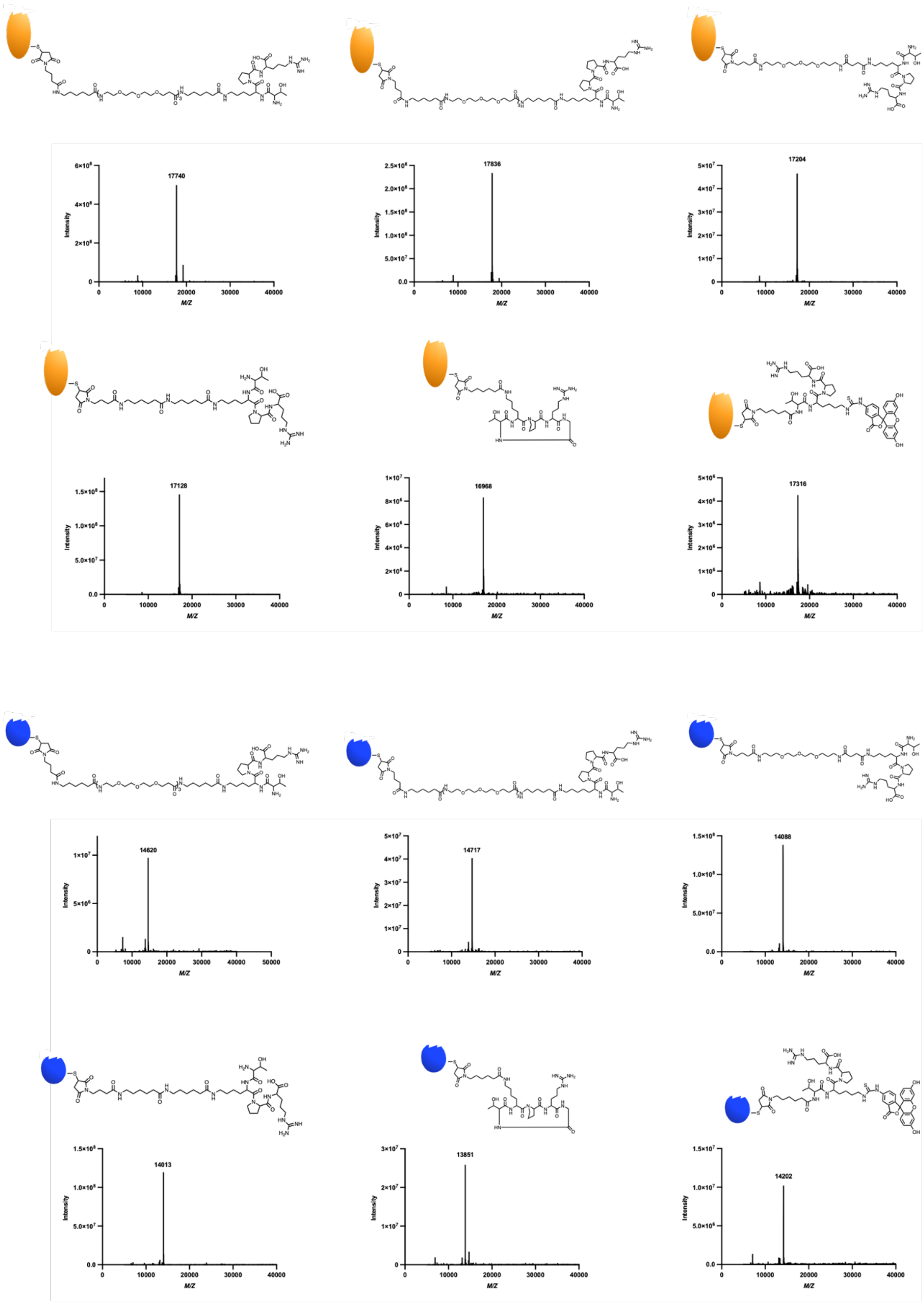

**Figure S6. LC-MS analysis and characterization of PSEs**

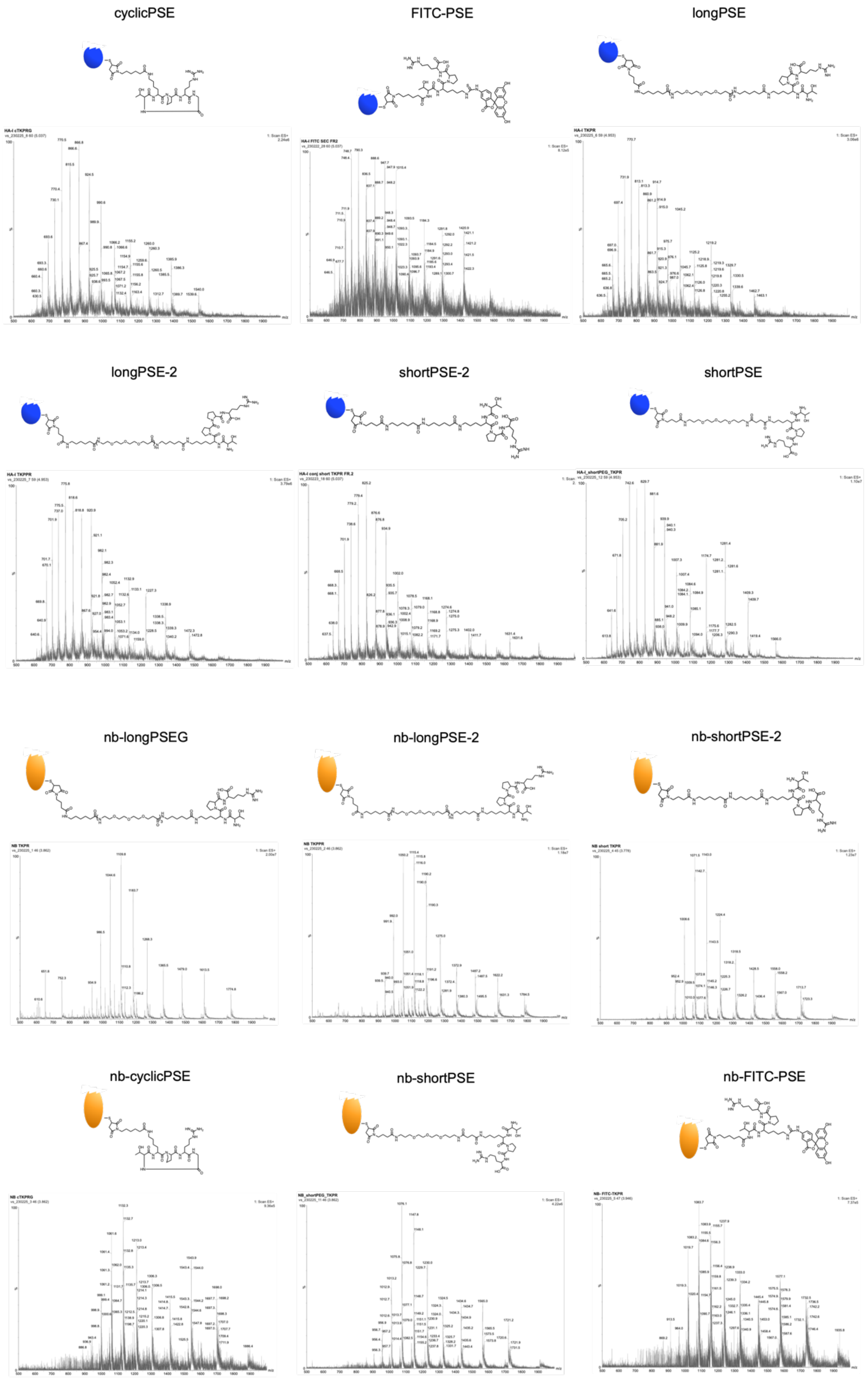

**Figure S7. LC-MS analysis and characterization of PSEs.**

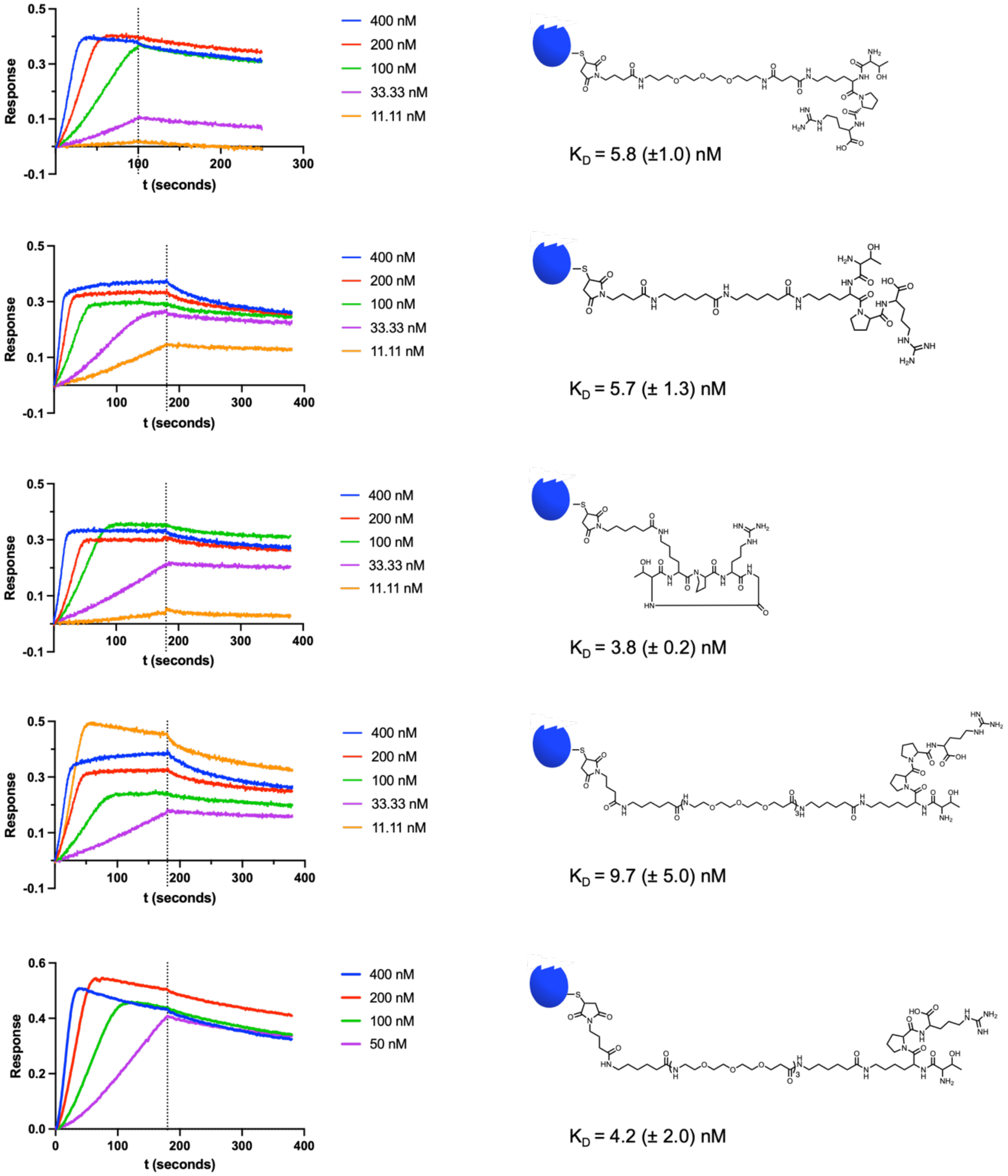

**Figure S8. Octet BLI analysis and $K_D$ determination of PSEs.** Biotinylated ligand loaded onto the strepatividin sensors at 2 μg/mL concentration. To determine the dissociation constants, the binding curves were fitted by non-linear regression analysis on prism.

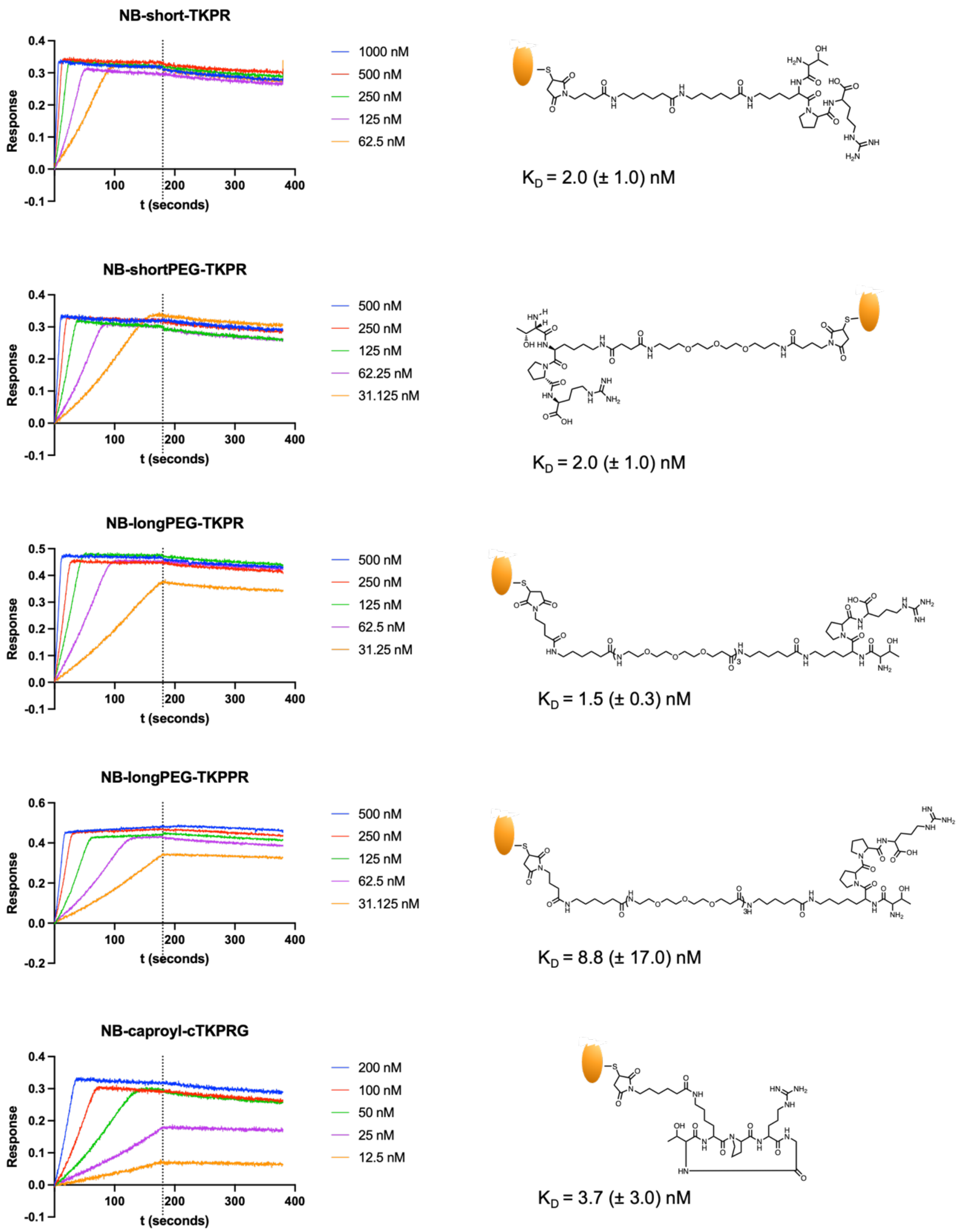

**Figure S9. Octet BLI analysis and K$_D$ determination of nb-PSEs.** Biotinylated ligand loaded onto the streptavidin sensors at 2 μg/mL concentration. To determine the dissociation constants, the binding curves were fitted by non-linear regression analysis on prism.

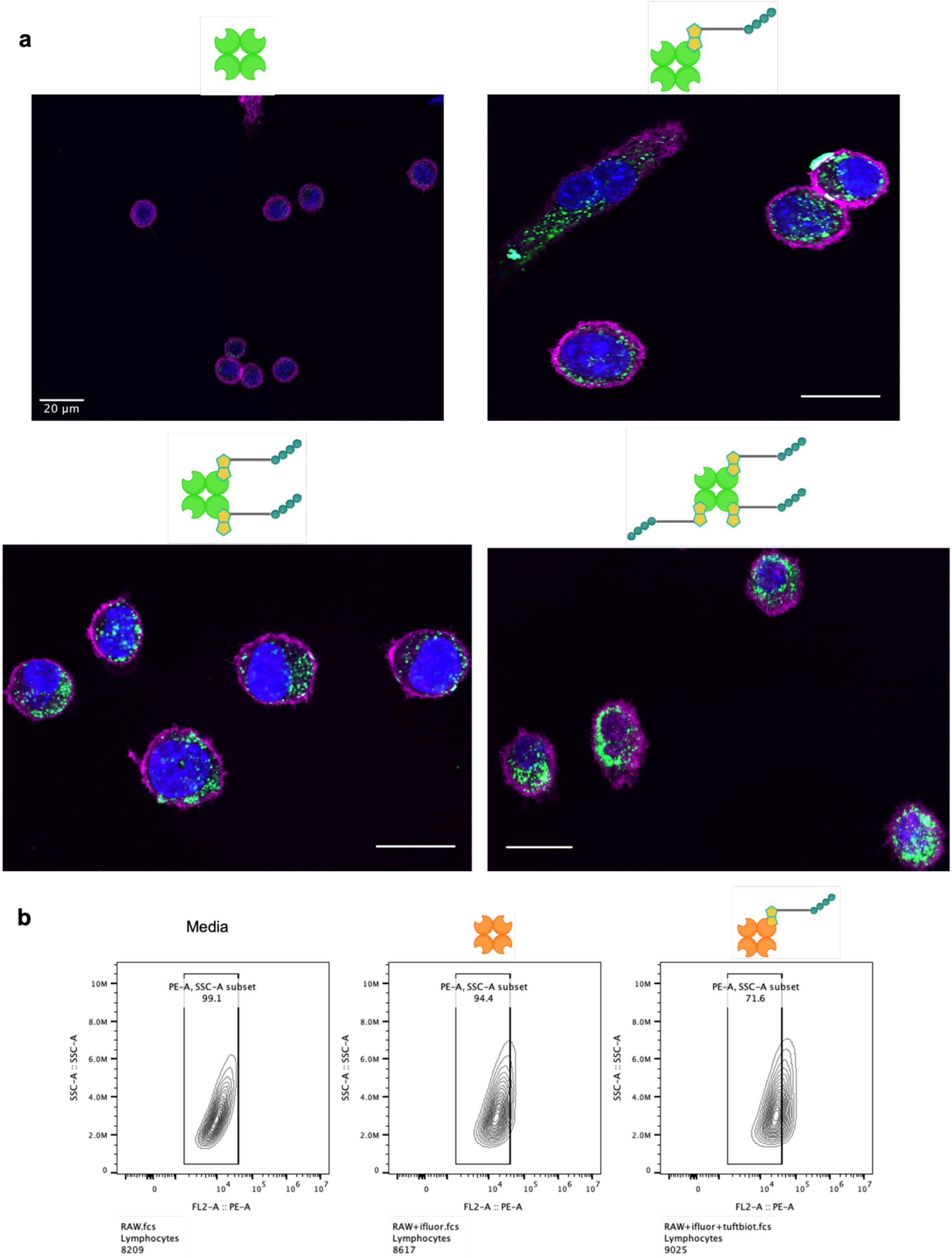

**Figure S10. RAW fluorescent protein uptake studies a**) RAW cells uptake of the fluorescent protein $NOG_{488}$ by confocal microscopy analysis. Scale bar: 20 μm. **b**) RAW cells uptake of the fluorescent protein Streptavidin $iFluor_{555}$ by flow cytometry analysis. The percentage of PE-positive cells is given by: (% PE-negative cells treated with only media) – (% PE-negative cells treated with $iFLuor_{555}$/Biot-$iFLuor_{555}$).

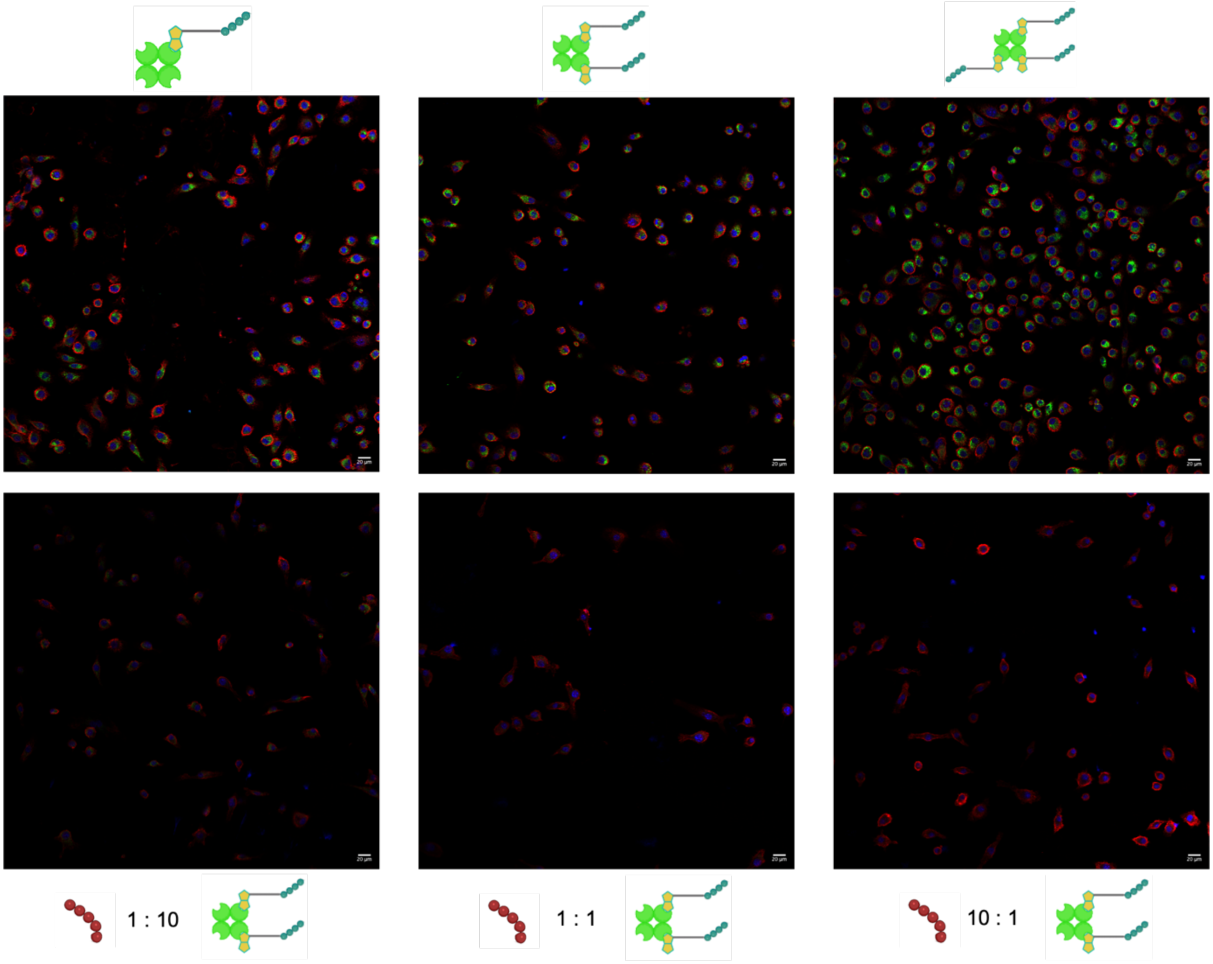

**Figure S11. Inhibition of the uptake of the fluorescent protein NOG$_{488}$ in the presence of the antagonist TKPPR by confocal microscopy analysis.** Confocal images of Biot-TKPR complex internalizing in RAW macrophages (upper panel) and Biot-TKPR in the presence of antagonist TKPPR (lower panel, from left ot right ratios TKPPR : Biot-TKPR 1:10, 1:1, 10:1). Scale bar: 20 μm.

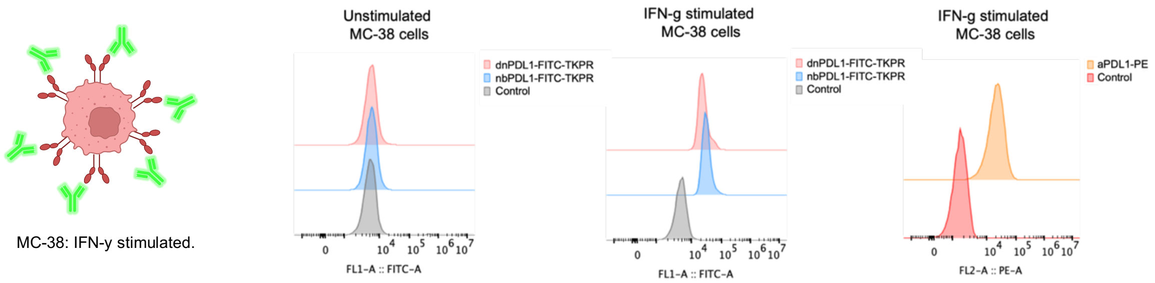

**Figure S12. PD-L1 binding of MC-38 cells stimulated with IFN-g.** The probes FITC-PSE, nb-FITC-PSE and the commercial antibody aPDL1-PE were incubated for 4 hours at 37 °C.

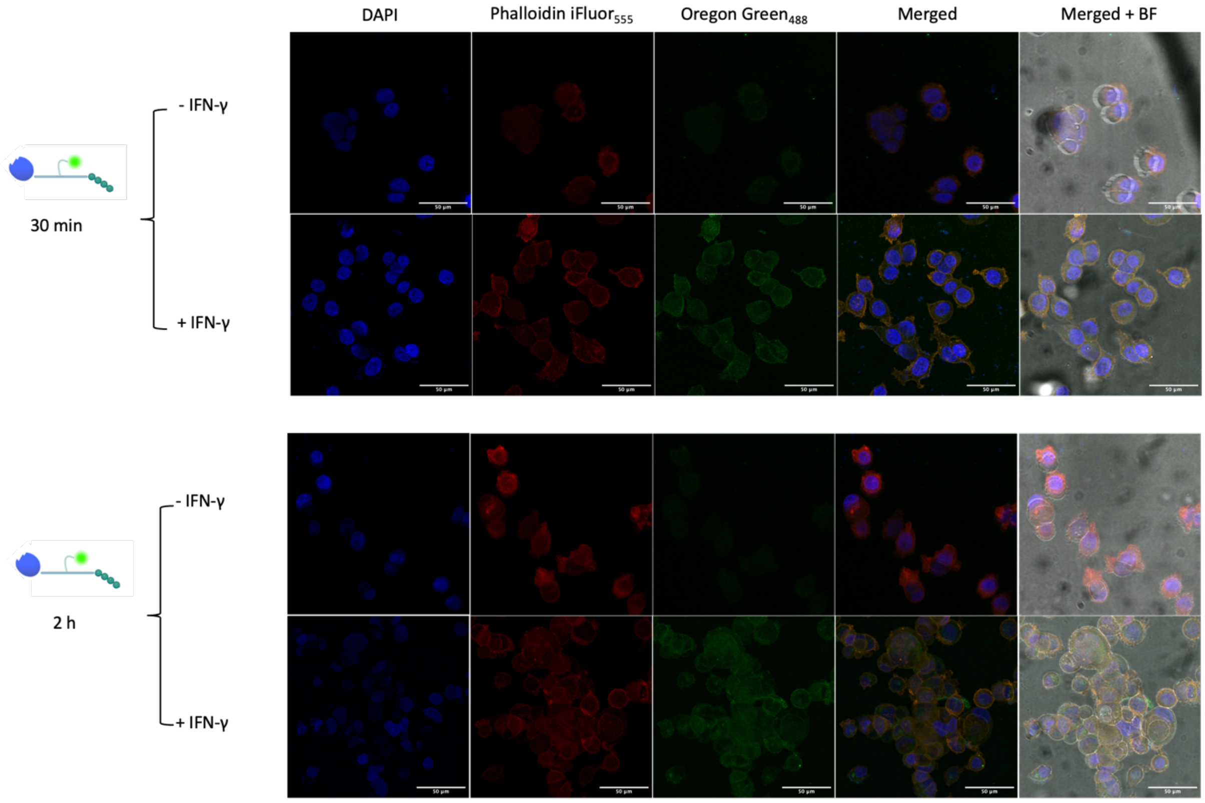

**Figure S13.** PD-L1 binding assays with MCF-7 unstimulated and stimulated with IFNγ FITC-PSE is incubated at 30 min and 2h with MCF-7 cells either unstimulated or stimulated for 24h with IFNγ to overexpress PD-L1. The cell membrane is stained with the fluorescent probe FITC-PSE only when MCF-7 cells are treated with IFNγ. FITC-PSE concentration: 100 nM. Scale bar: 50 μm.

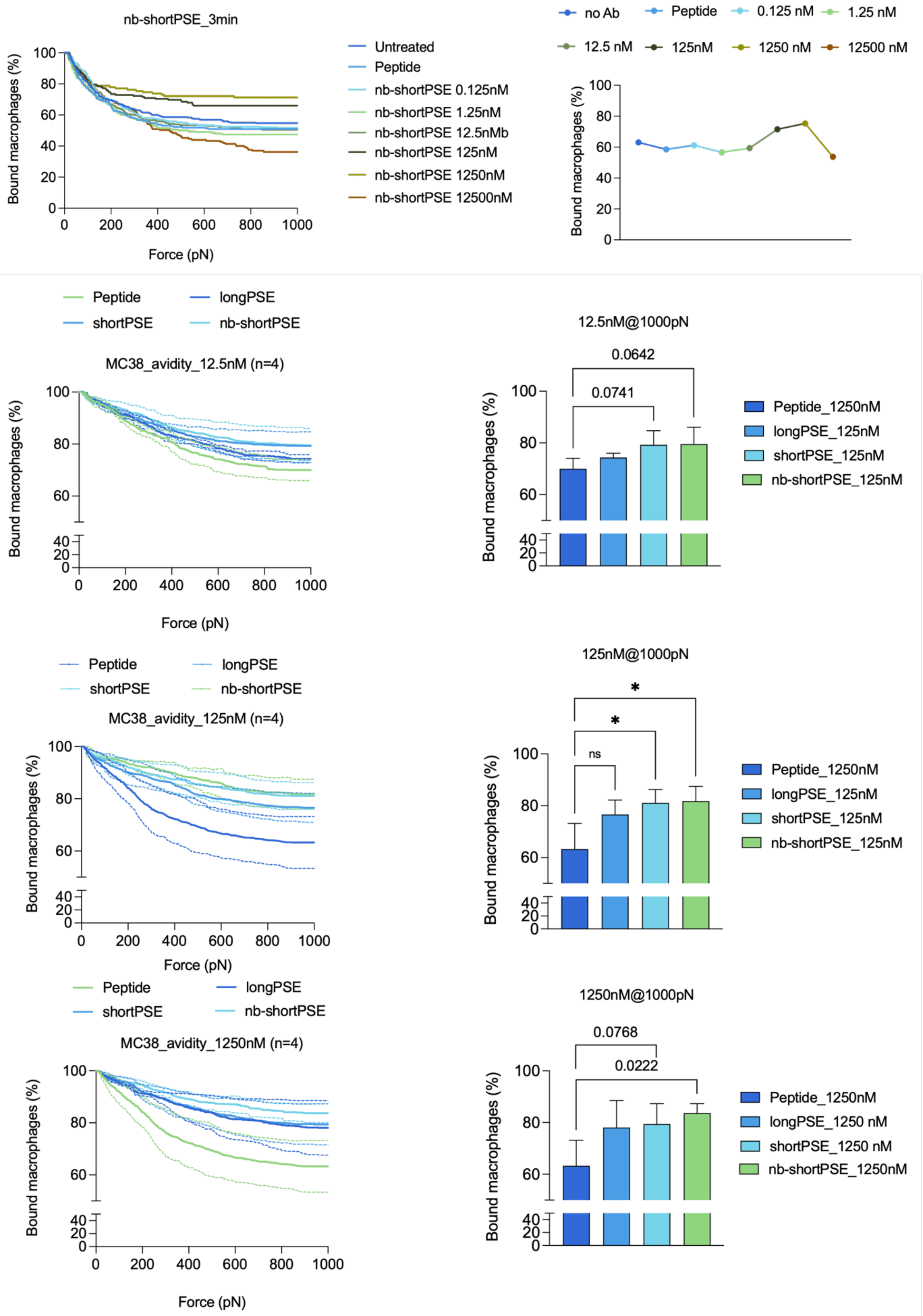

**Figure S14. Titration and avidity measurements with selected molecules at 12.5 nM, 125 nM, and 1.25 μM.** Titration curve of nb-shortPSE in co-culture system MC-38/RAW 264.7. The effect of concentration on the binding of Mϕ to cancer cells was measured by z-movi cell avidity analyser after 3-min incubation with the molecule. The effect of avidity was compared to a blank (no Ab) and the peptide tethered to the Short linker (Peptide).

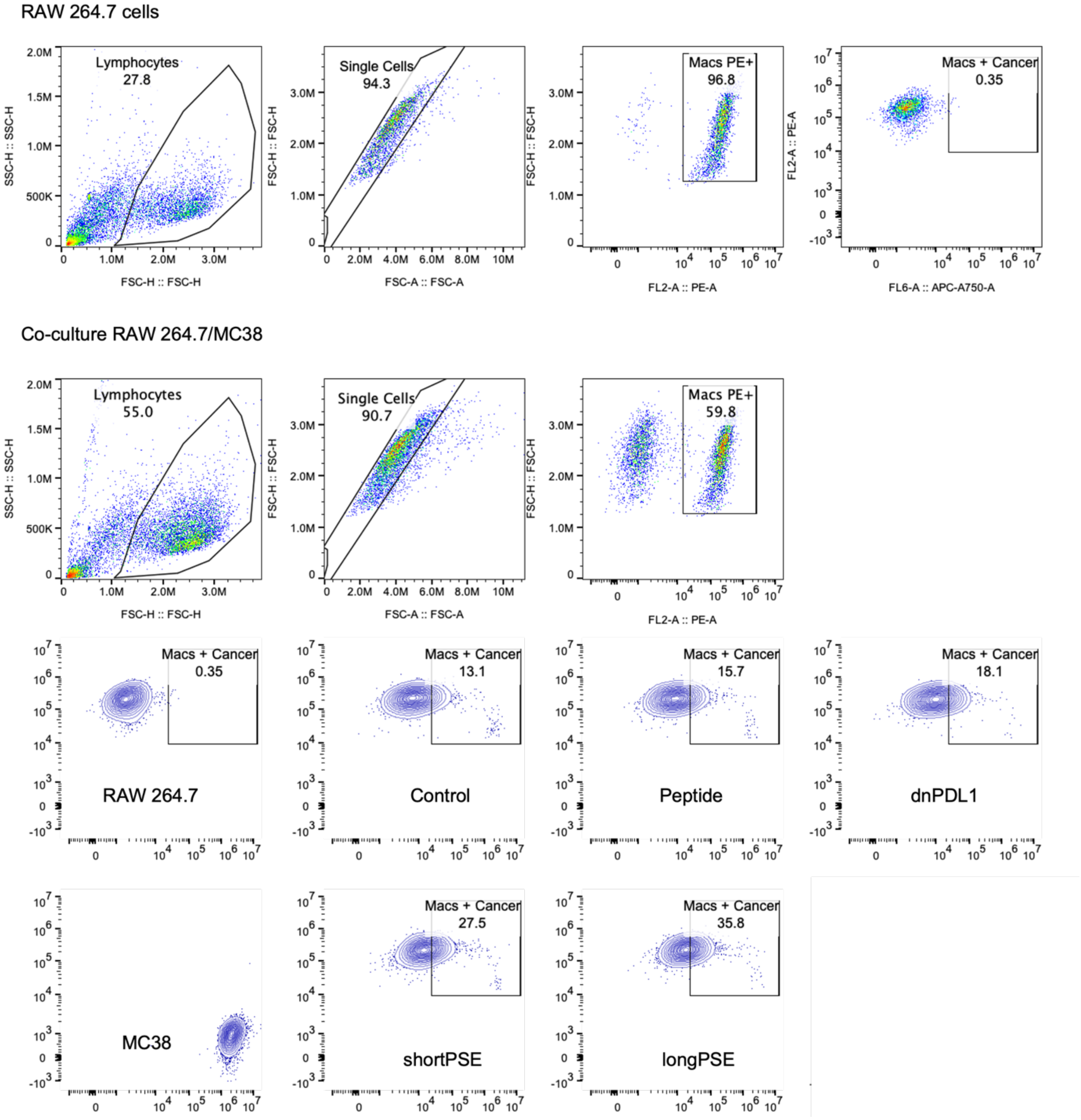

**Figure S15. Gating strategy phagocytosis assay with RAW 264.7 Mϕ.**

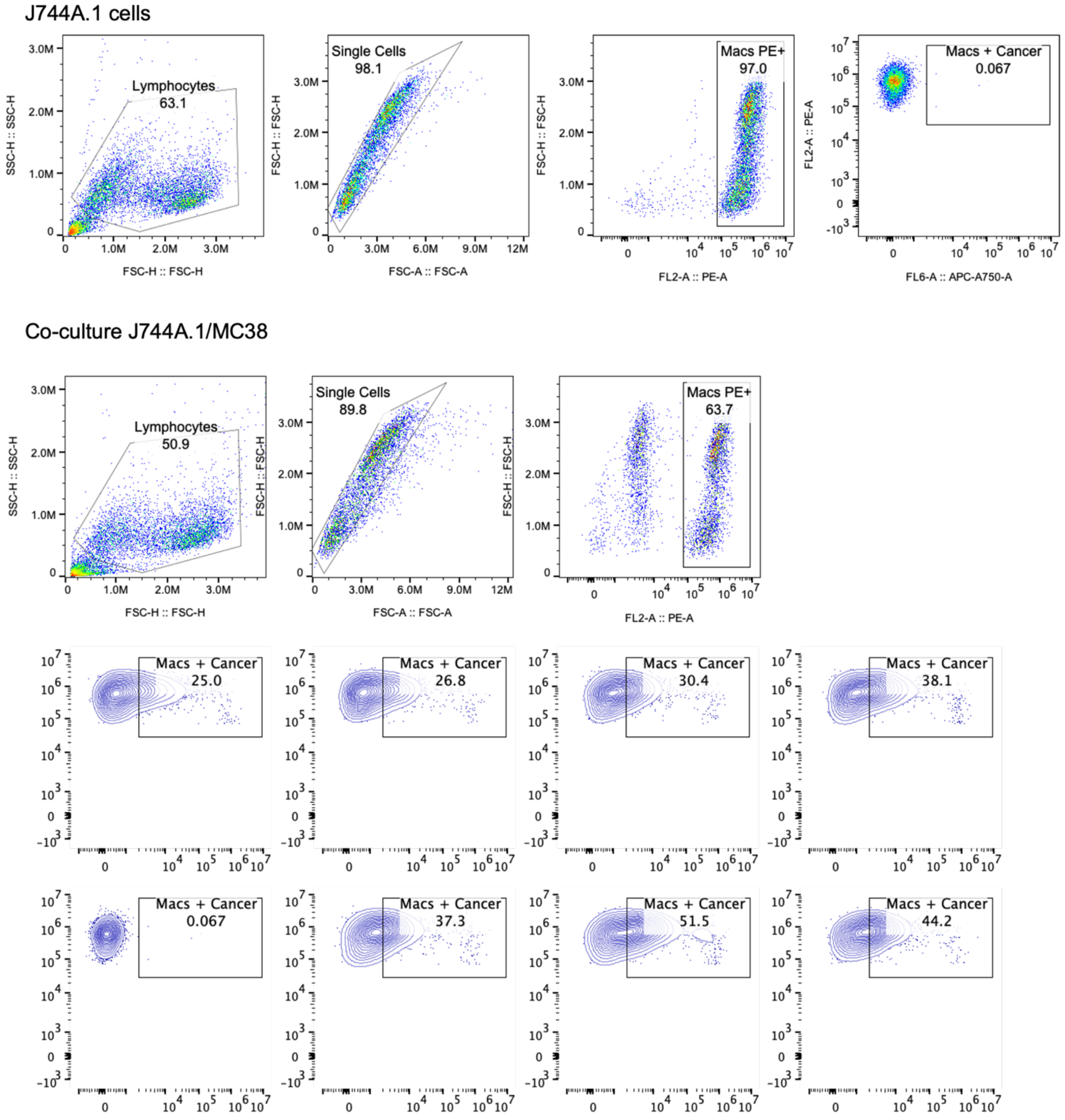

**Figure S16. Gating strategy phagocytosis assay with J774A.1 Mϕ.**

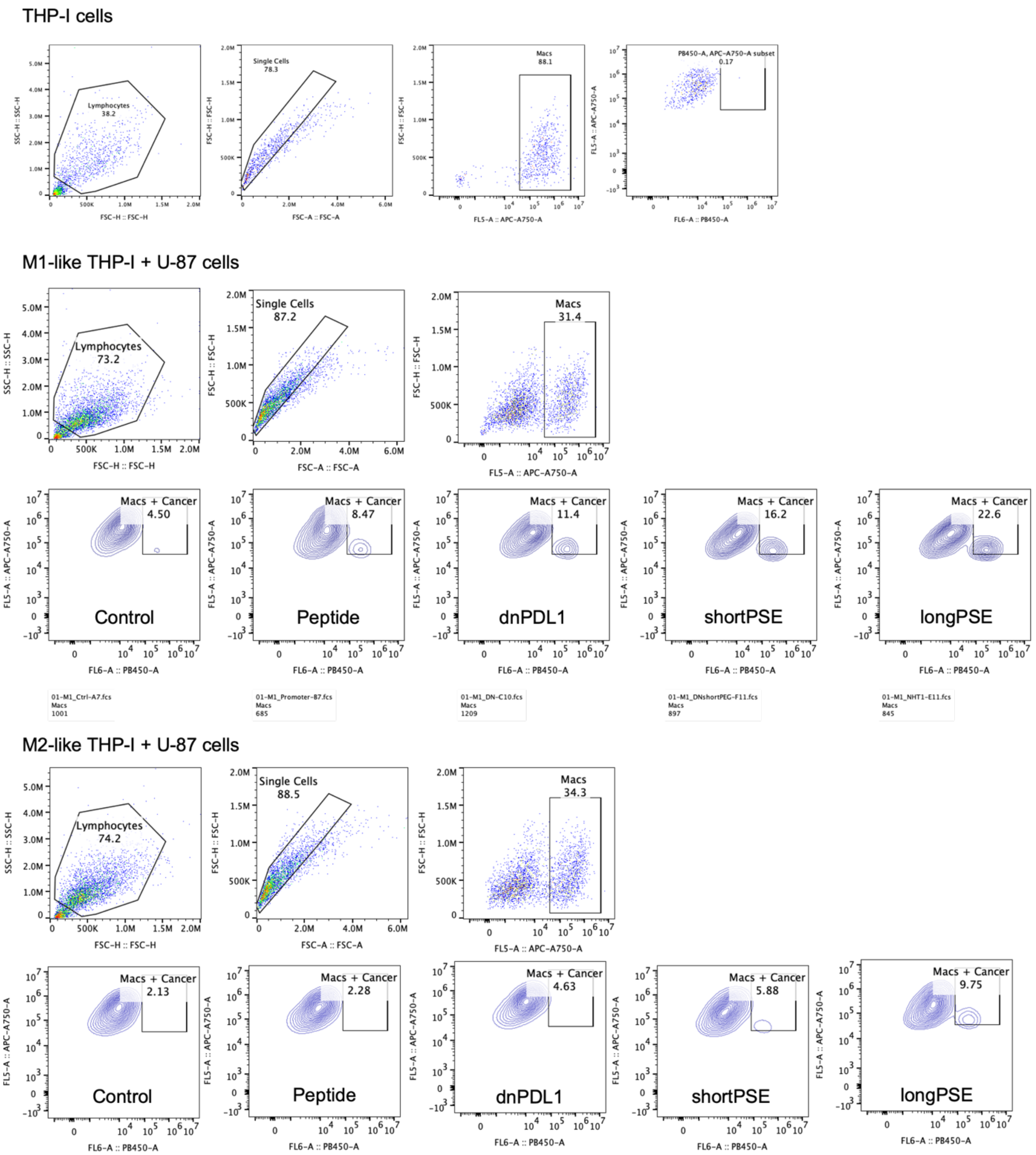

**Figure S17. Gating strategy phagocytosis assay with THP-I Mϕ.**

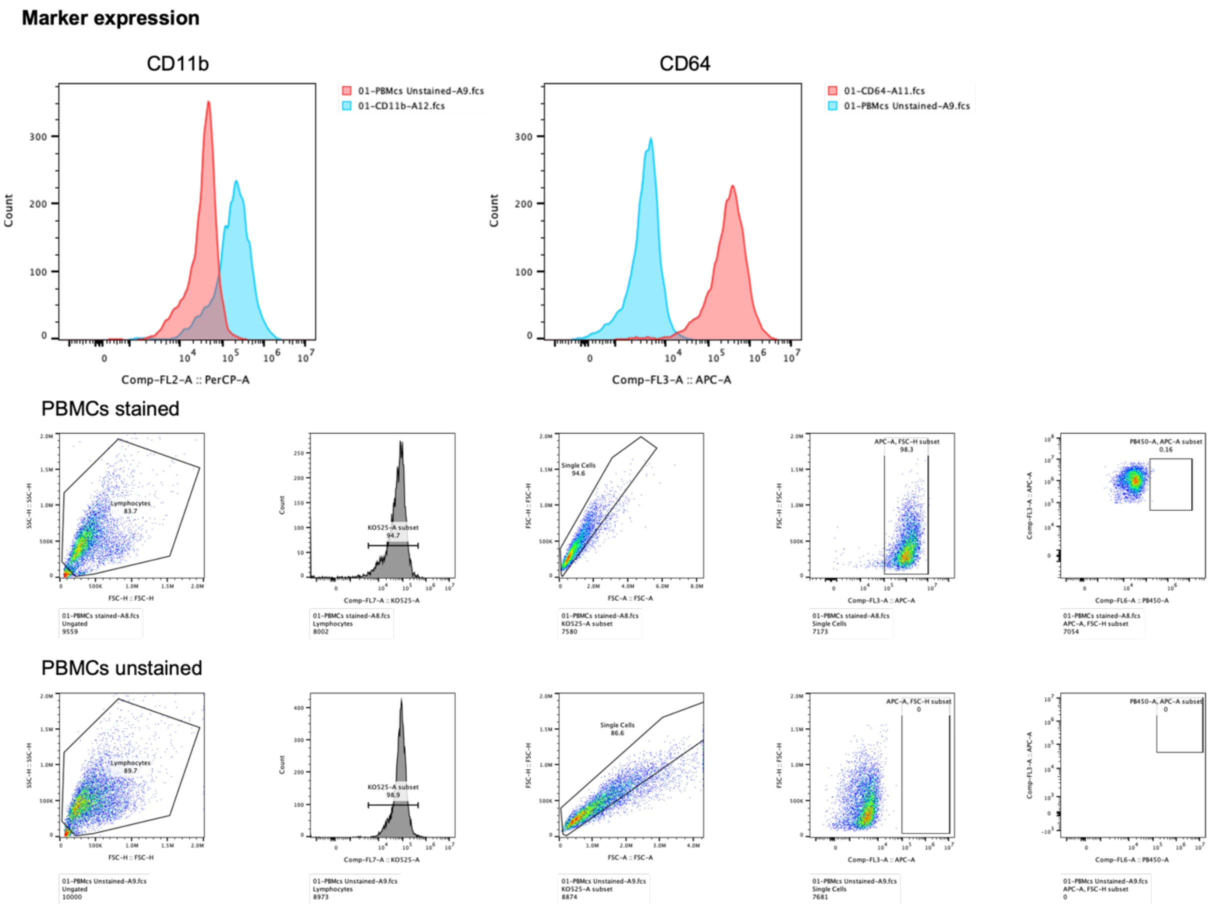

**Figure S18. Characterization of isolated PBMCs and staining for phagocytosis assays.**

a

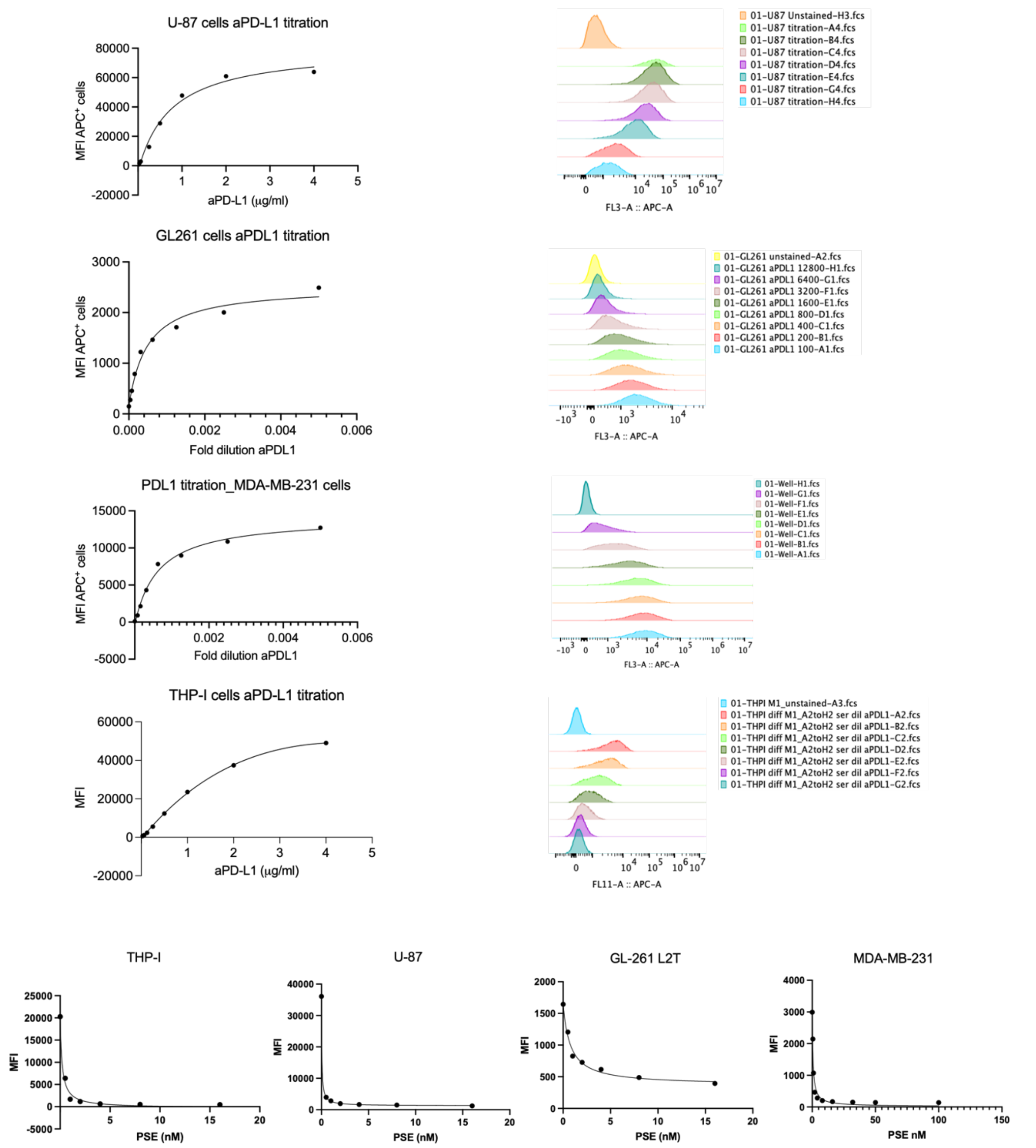

**Figure S19. a,** Titration assays of aPDL1-APC with PD-L1-expressing cell lines. aPDL1-APC dilutions used: 1:100, 1:200, 1:400, 1:800, 1:1600, 1:3200, 1:6400, 1:12800. **b**, Blocking assays using fixed concentration of antibody (aPDL1-APC, 1:500 dilution, 133 ng/ml) with different concentrations of longPSE (0.5-16 nM).

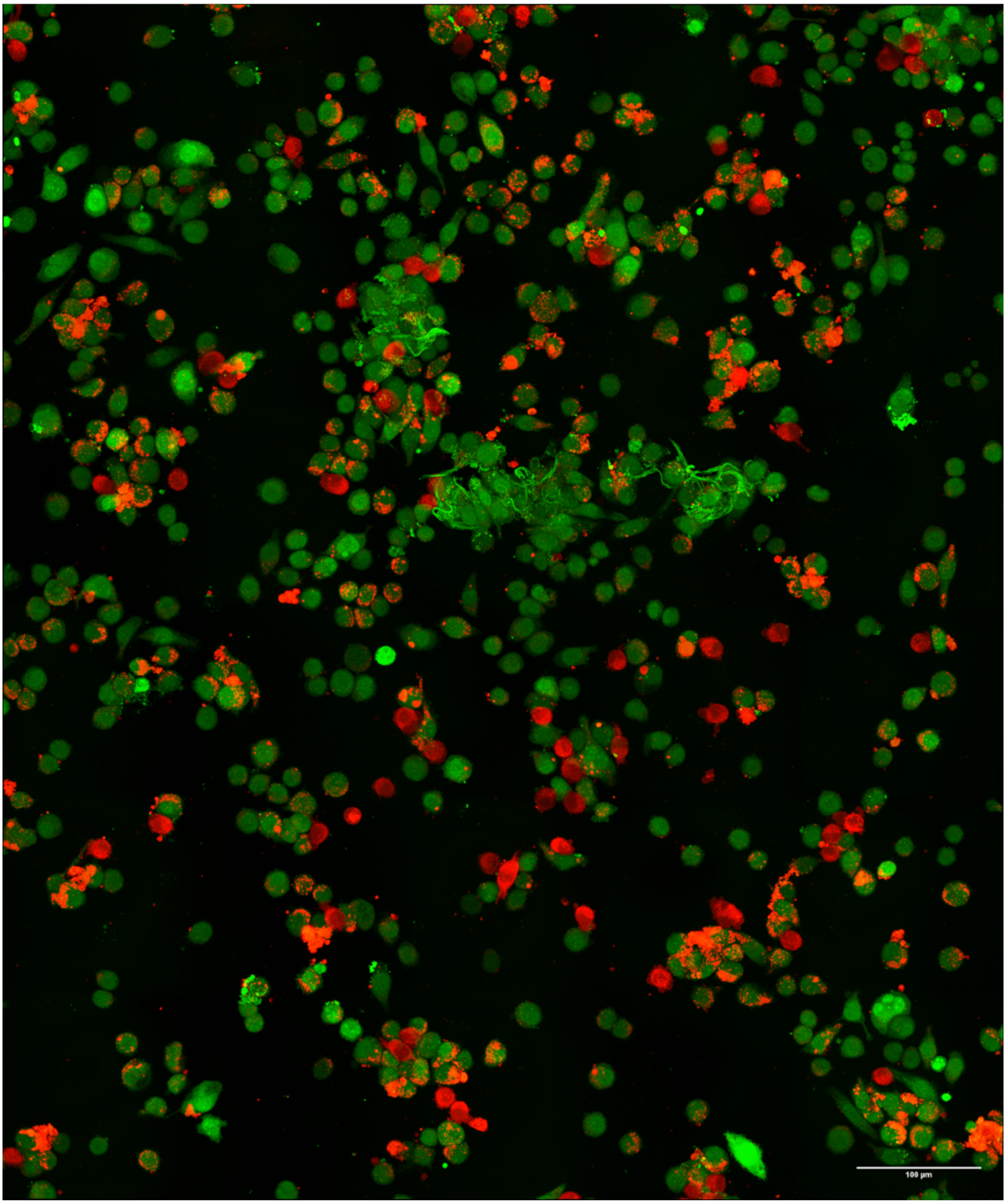

**Figure S20. Image relative to Figure 2b in the main text. Control (A).**

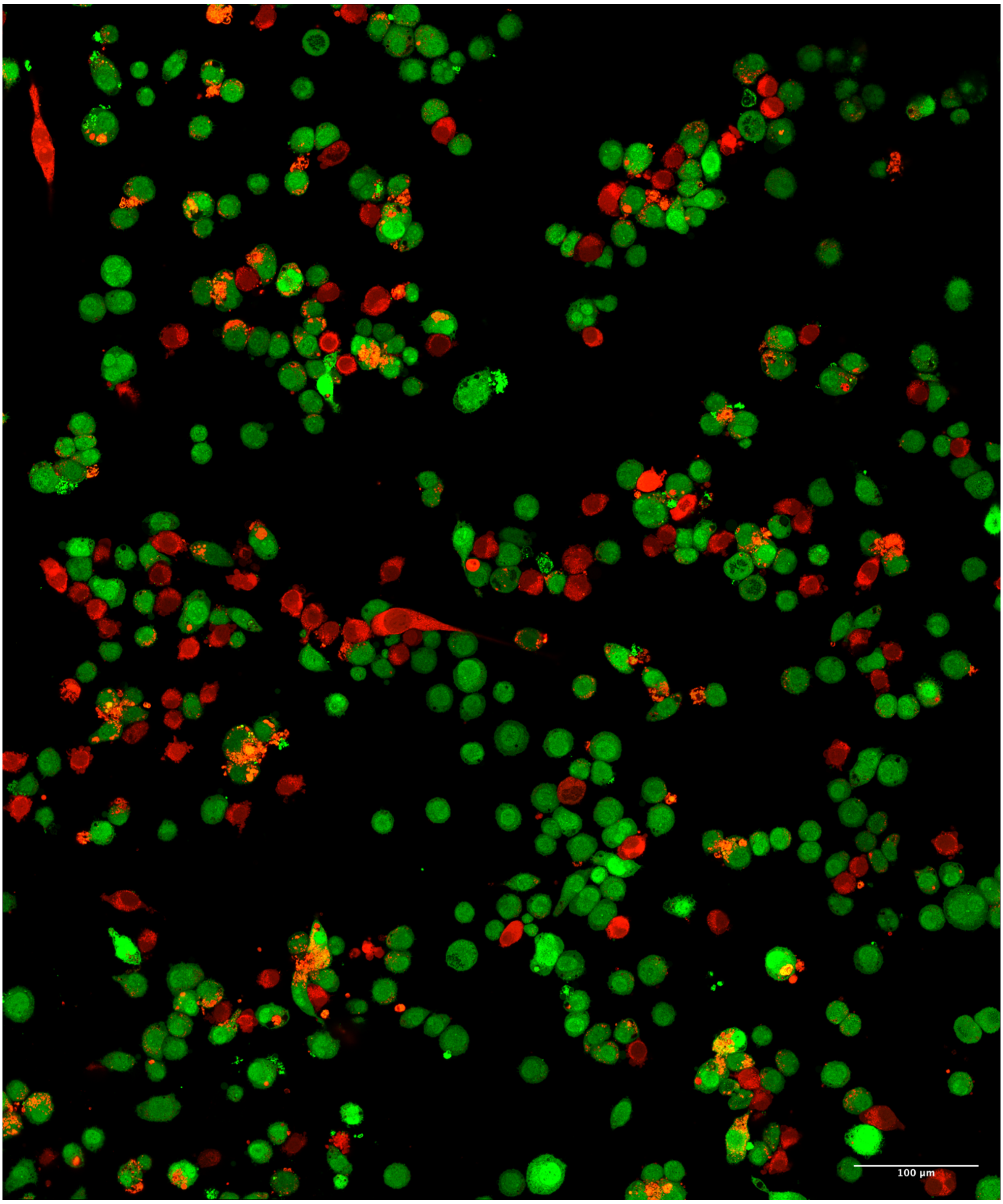

**Figure S21. Image relative to Figure 2b. Peptide (B).**

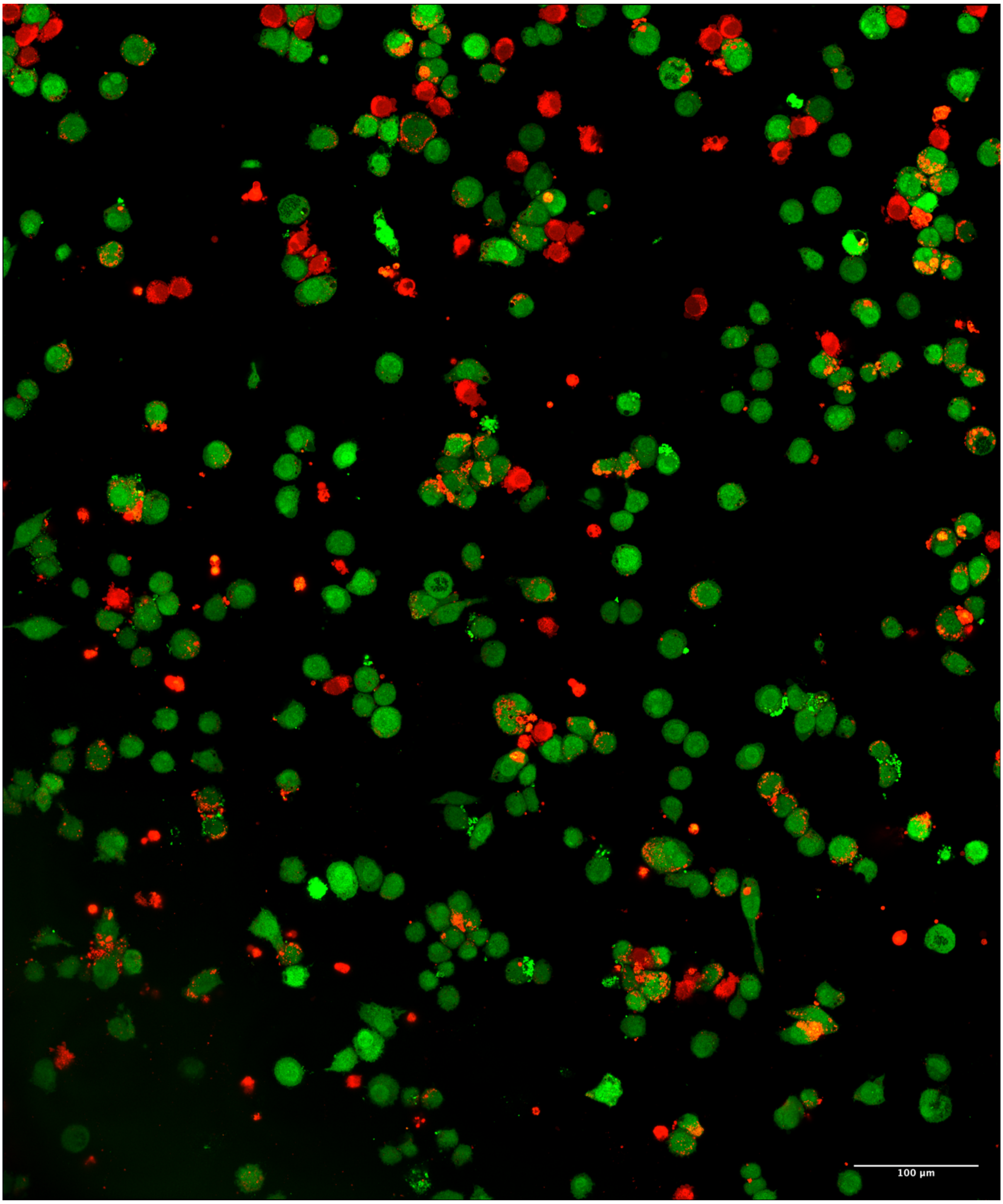

**Figure S22. Image relative to Figure 2b. dnPDL1 (C).**

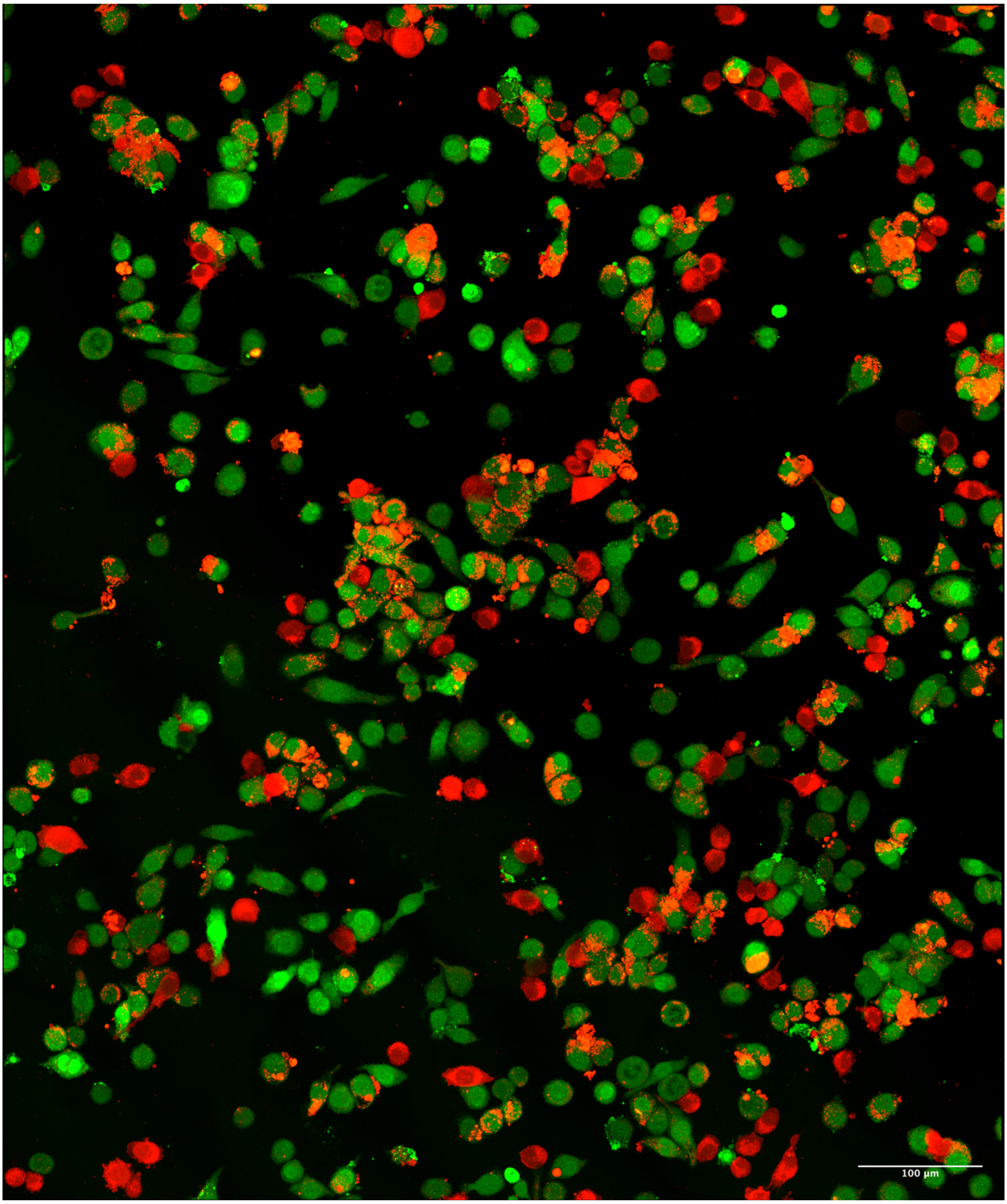

**Figure S23. Image relative to Figure 2b. shortPEG (D).**

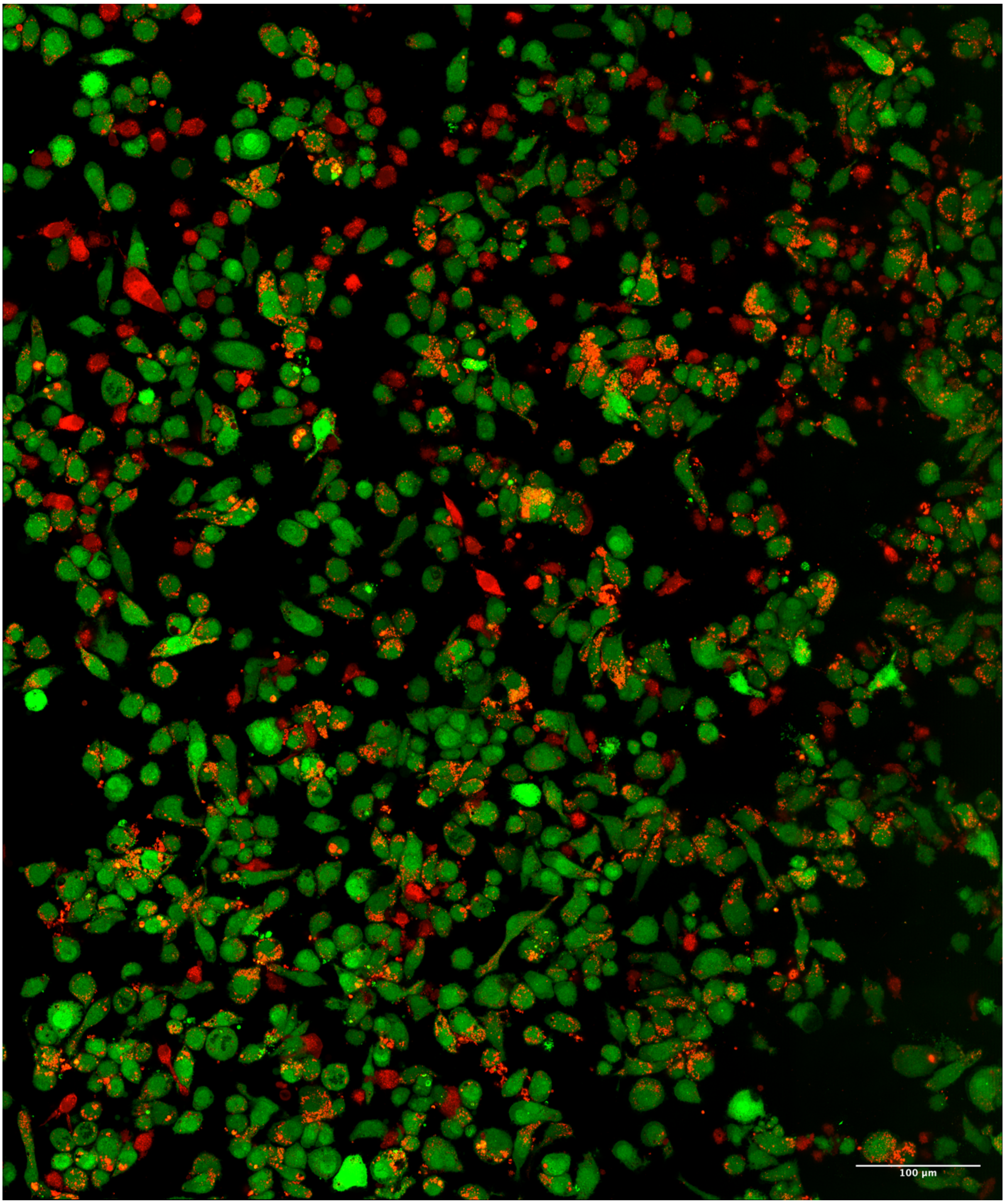

**Figure S24. Image relative to Figure 2b. longPSE (E).**

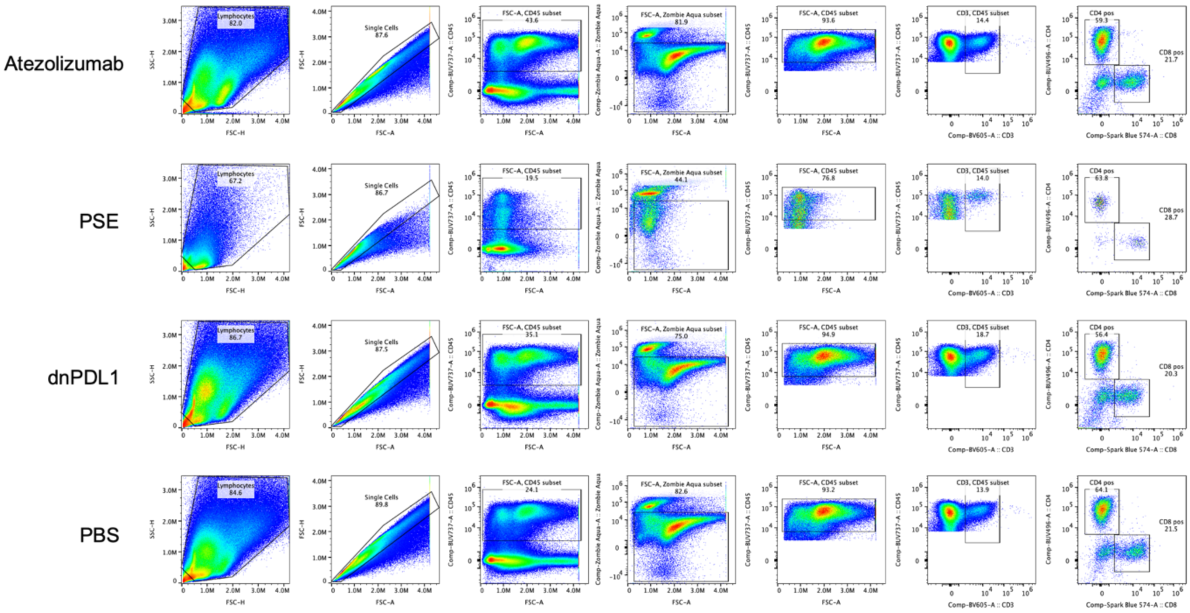

**Figure S25. Gating strategy for the analysis of the lymphoid compartment of the tumor-bearing hemisphere of mice treated with PSE (longPSE) and relative controls.** T-cells were gated as Single/Live/CD45+/CD3+.

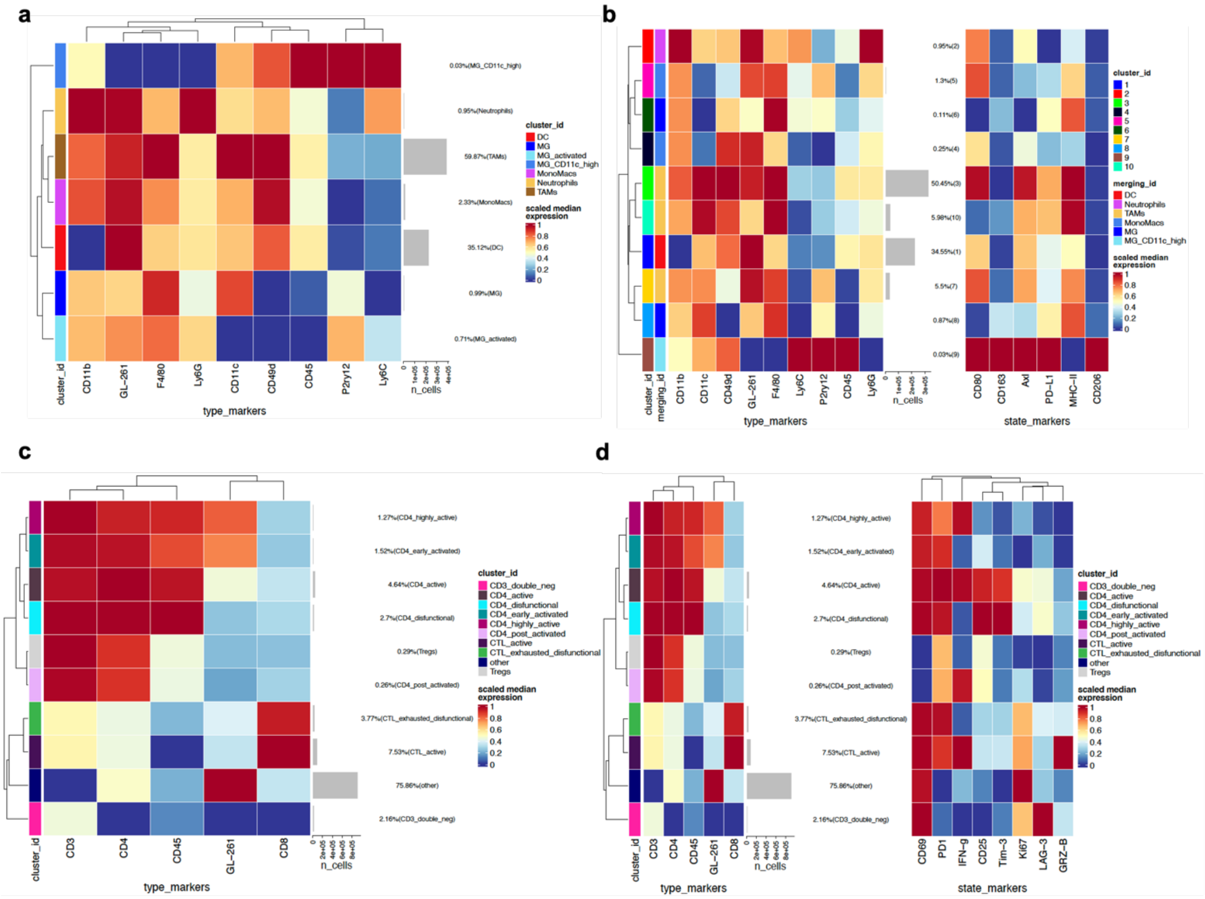

**Figure S27. Clustering analysis and cell annotation by FlowSom analysis.**
**a,** Myeloid panel clustered heatmap of scaled median marker expression per cluster. The number of cells per cluster is indicated on the right y-axis. **b,** Merged clustered heatmap of scaled median marker expression per cluster separated by type and state. **c,** Lymphoid panel clustered heatmap of scaled median marker expression per cluster. The number of cells per cluster is indicated on the right y-axis. **d,** Merged clustered heatmap of scaled median marker expression per cluster separated by type and state.

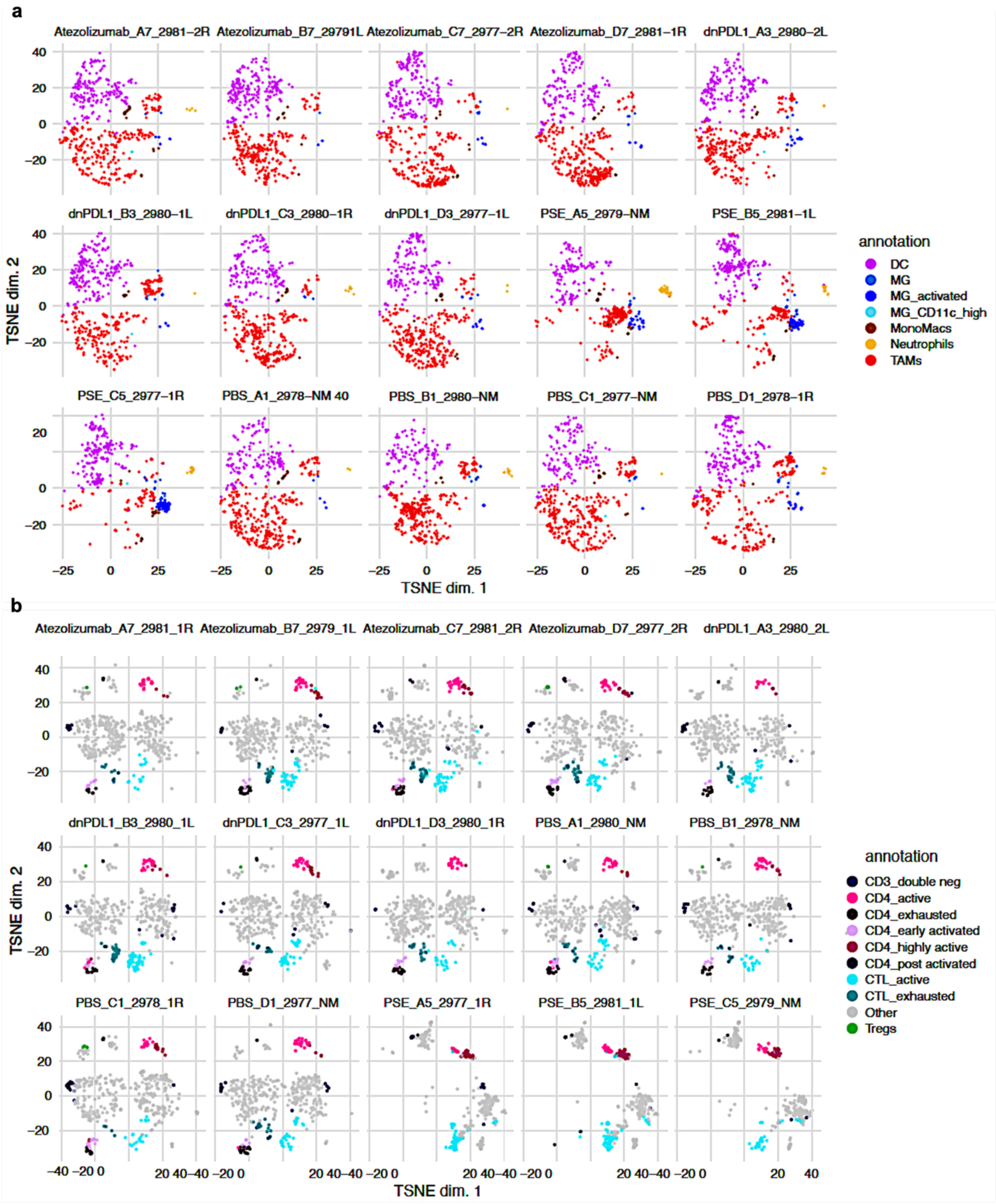

**Figure S28. Clustering analysis of myeloid and lymphoid cell populations of tumor bearing hemisphere treated samples. a,** t-distributed stochastic neighbor embedding (TSNE) plots of all animal samples depicting 7 main myeloid populations after initial merging and manual annotation from 13 populations originating from unbiased clustering. **b,** t-distributed stochastic neighbor embedding (TSNE) plots of all animal samples depicting 7 main lymphoid populations after initial merging and manual annotation from 20 populations originating from unbiased clustering.

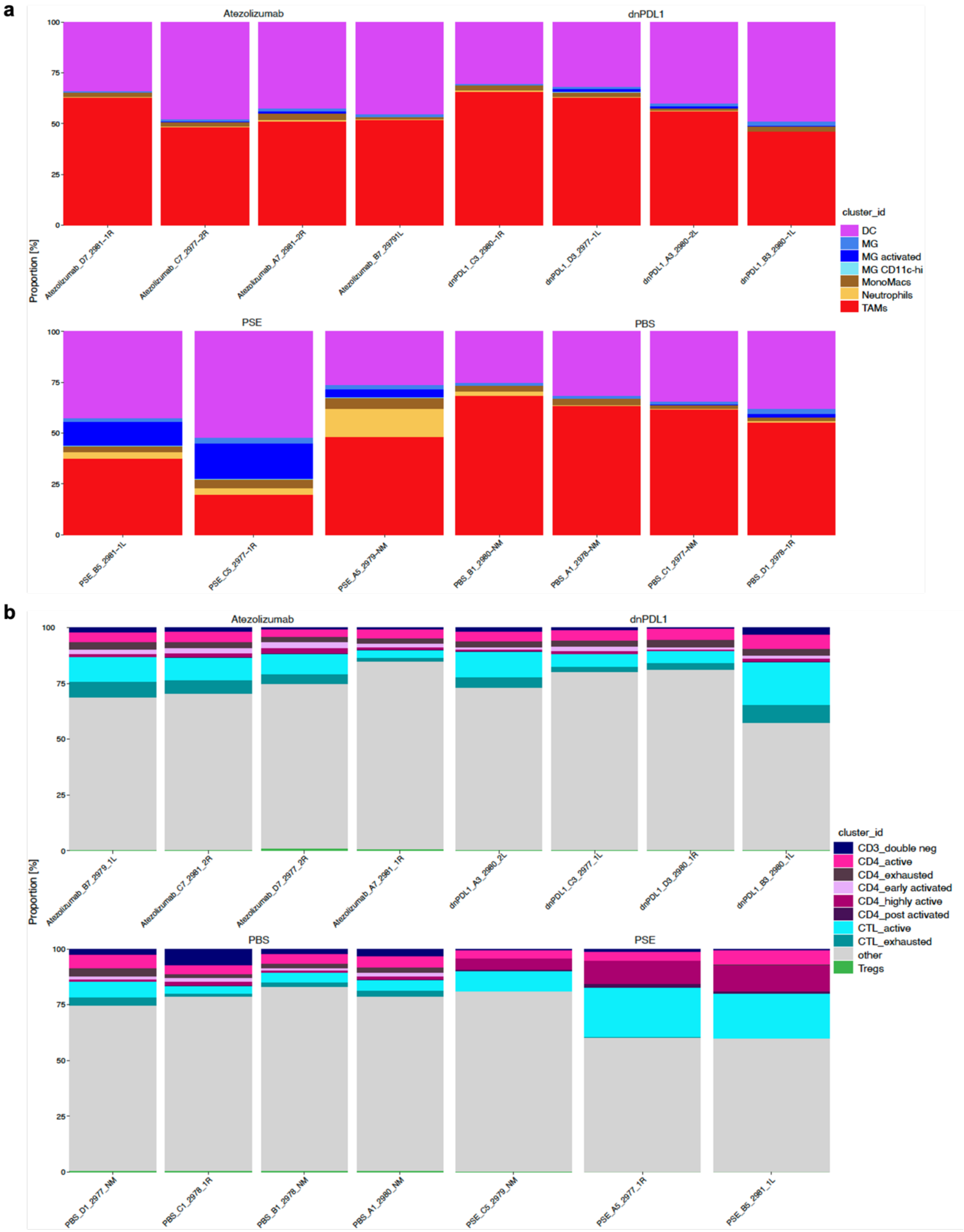

**Figure S29. Stacked bar plots representing cluster frequencies in the myeloid compartment (a) and lymphoid compartment (b) per each experiment and condition.**

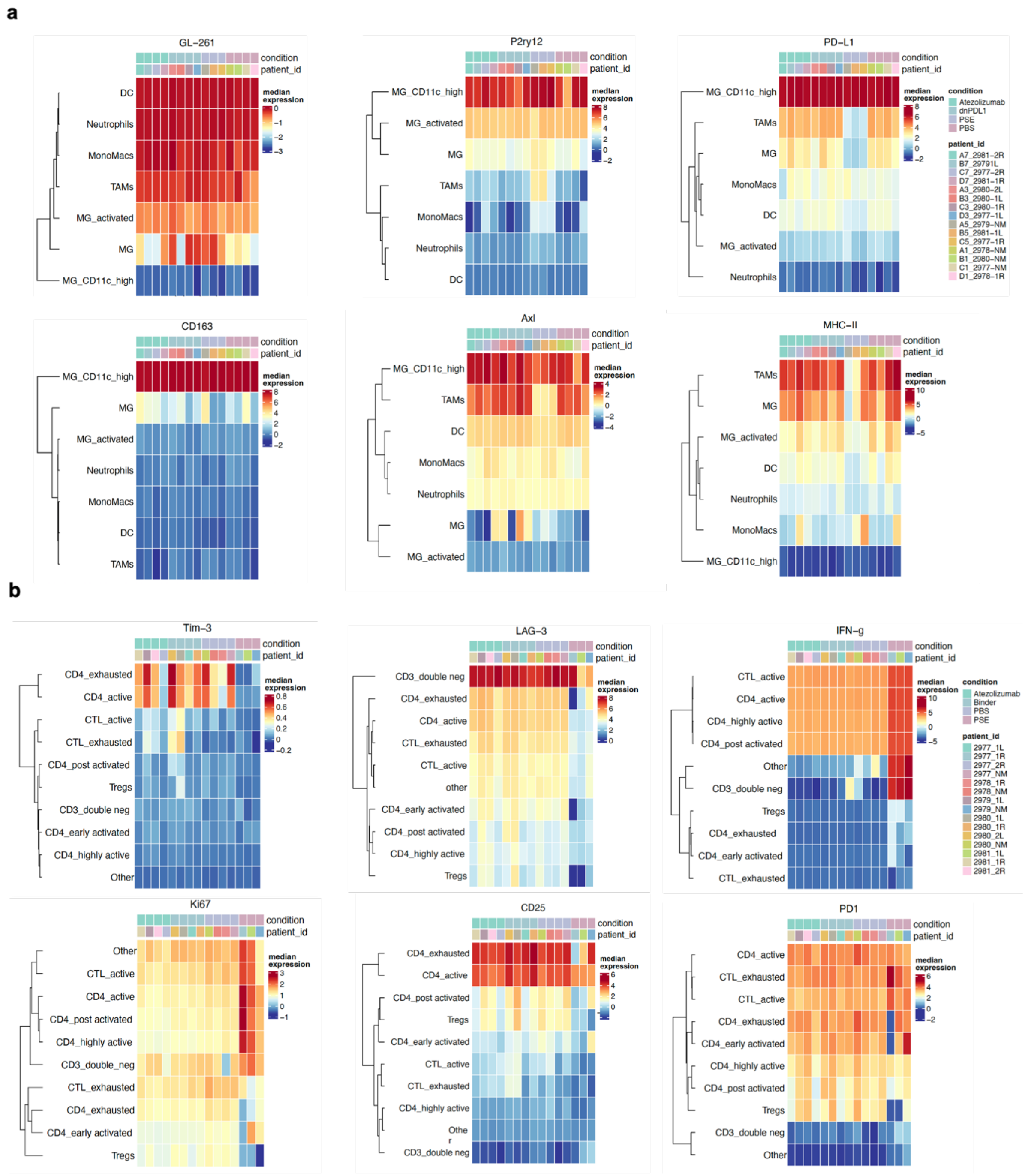

**Figure S30. Differential expression analysis in myeloid and lymphoid cell subsets reveal key phenotypic differences in PSE-treated mice.**

**a**, Heatmap of median expression for marker expression per individual sample depicting the changes in myeloid markers expression in PBS (n=4), PSE (n=3), dnPDL1 (n=4), Atezolizumab (n=4). PSE-treated brains show an increase of phagocytosed GL-261 cells in MG (left diagram), TAMs show higher expression of P2ry12 (middle diagram), and PDL1 is downregulated in TAMs, MG and DCs. **b,** Heatmap of median expression for marker expression per individual sample depicting the changes in lymphoid markers expression in PBS (n=4), PSE (n=3), dnPDL1 (n=4), Atezolizumab (n=4). Key changes in mice treated with the PSE are characterized by the decrease in exhaustion markers TIM-3 and LAG-3 (left and middle diagram) and increase of IFNγ (right diagram) and the proliferation marker Ki67 in CD4/CD8 T cells.

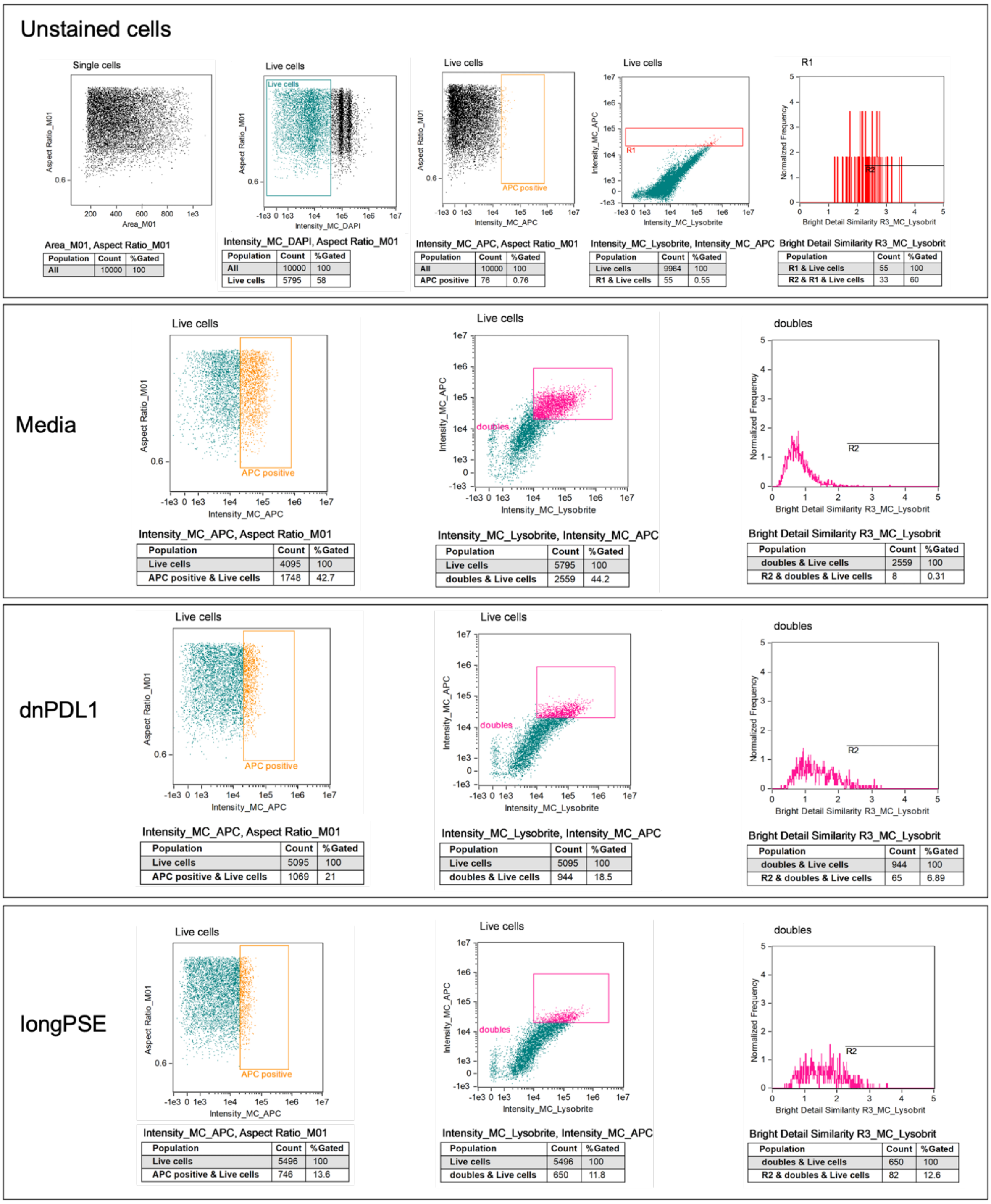

**Figure S31. Imagestream analysis of THP-I cells treated with Media only, dnPDL1 and longPSE**. Unstained diagrams show the gating of single live cells and gates used for APC+ cells. Double positive cells (Lysobrite+ APC+) were gated for colocalization analysis using Ideas2.0 software. Cells with brigh detail similarity greater than 2 were gated as colocalized cells.

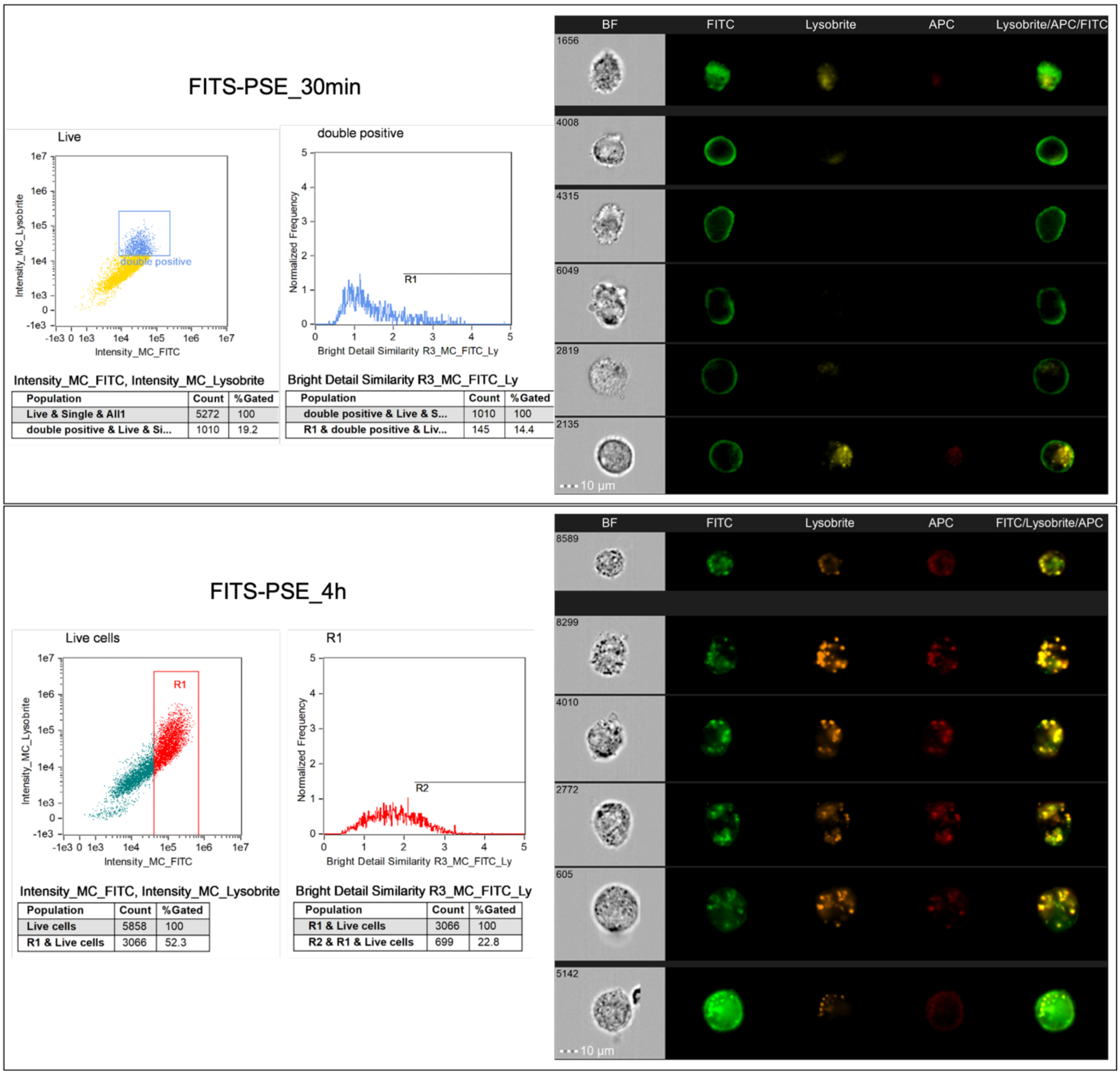

**Figure S32. Imagestream analysis of THP-I cells treated with FITC-PSE at 30min and 4h incubation**. Diagrams show the co-localization of double positive FITC+ Lysobrite+ cells at each timepoint. Channels from left to right: BF – brightfield; FITC – cells stained with FITC-PSE; Lysobrite - lysosome tracker; APC – cell stained with aPDL1-APC; FITC/Lysobrite/APC – all fluorescent channels merged. Analysis were performed using Ideas 2.0 software.

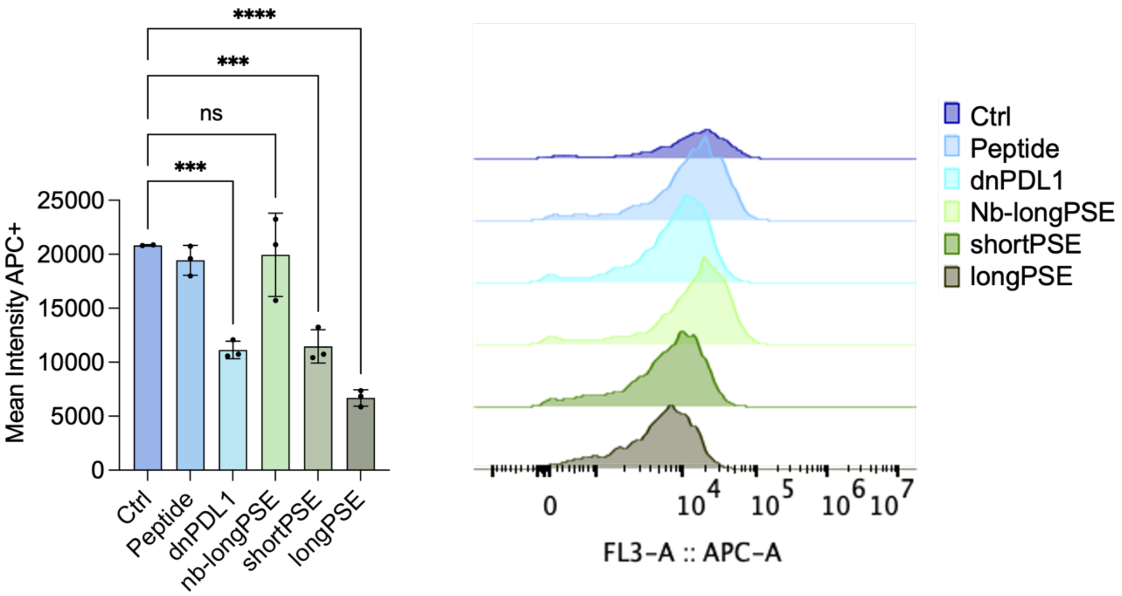

**Figure S33. Analysis of surface PDL1 by flow cytometry after 4h treatments.** aPDL1-APC was used to stain membrane PD-L1 after incubation with the molecules. The single-cell suspensions were analysed by flow cytometry. The bar graph on the left represents the mean intensity of membrane PD-L1 after each treatment. The histograms on the right are representative of the mean APC intensity for each treatment. Statistical analysis was performed in Prism (version 10) ordinary one-way ANOVAs were performed with Tukey's multiple comparison's test to compare treatment groups. In every instance the asterisk * indicates a $p<0.05$, ** indicates $p<0.01$, *** indicates $p<0.001$, and **** indicates $p<0.0001$.

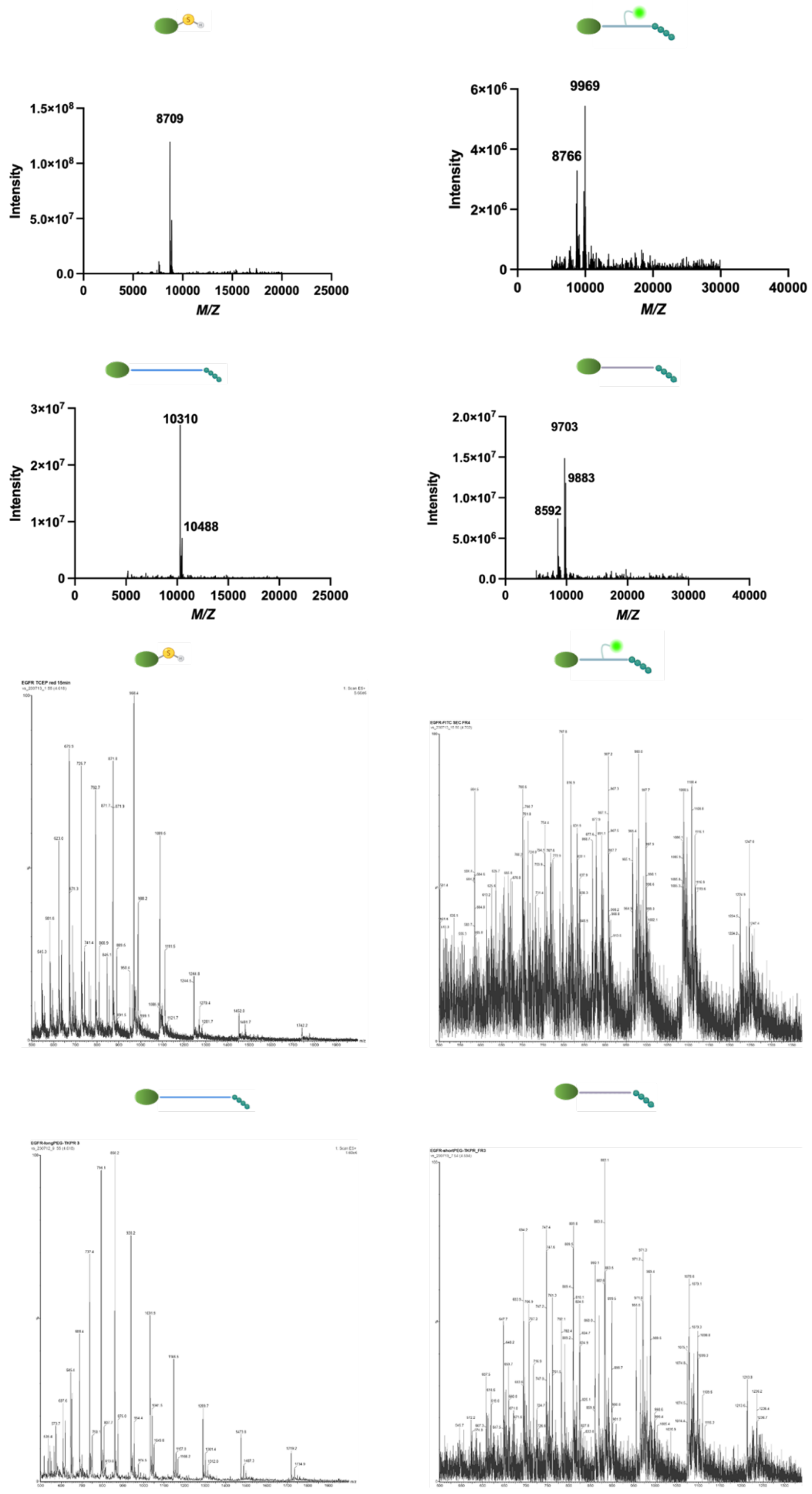

**Figure S34. LC-MS analysis of EGFR conjugates. Mass sequence of the EGFR conjugates.**

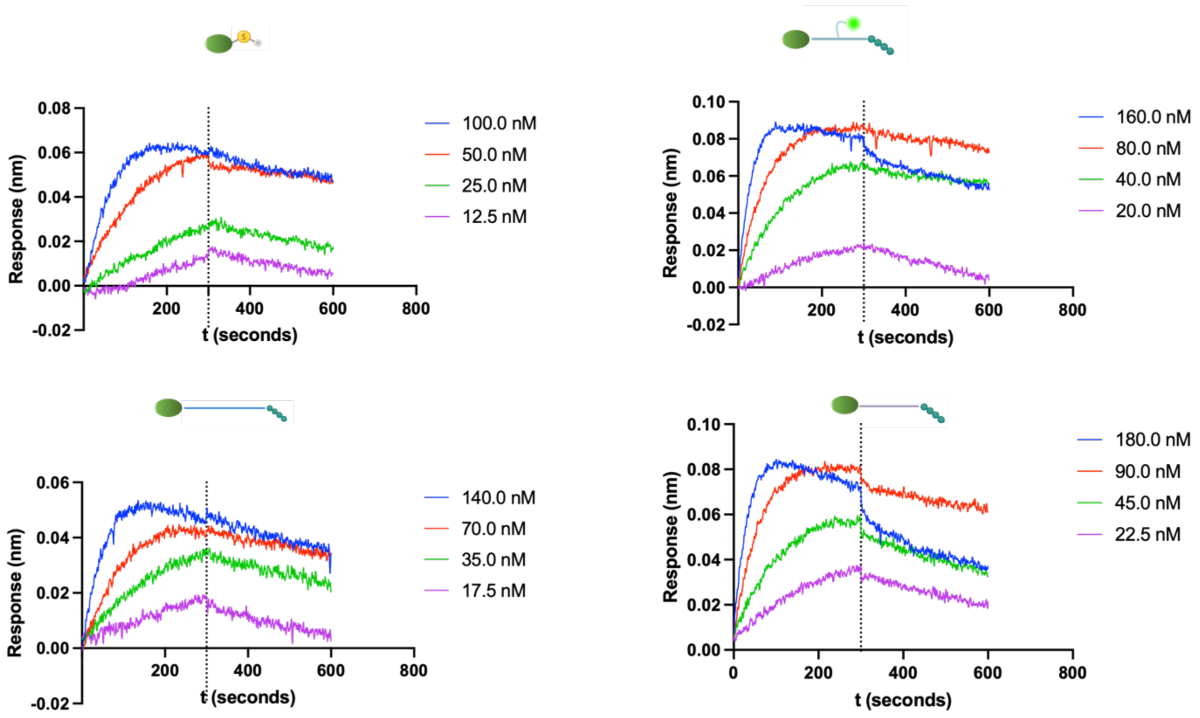

**Figure S35. BLI analysis of EGFR-conjugates.** Biotinylated ligand loaded onto the streptavidin sensors at 2 μg/mL concentration. Biotinylated ligand loaded onto the streptavidin sensors at 2 μg/mL concentration. The association and dissociation steps were analysed on prism.

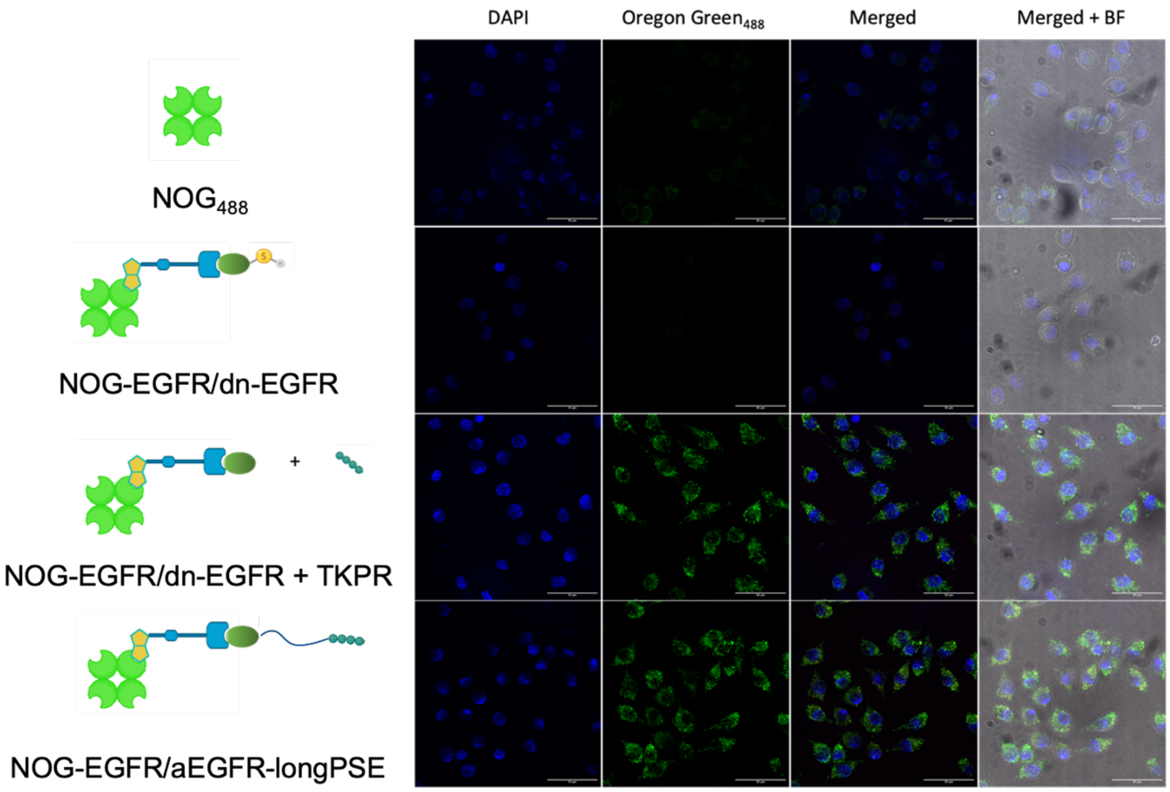

**Figure S36.** Confocal analysis of RAW 264.7 Mϕ treated with: $NOG_{488}$; $NOG_{488}$-EGFR complex + dnEGFR; C) $NOG_{488}$-EGFR complex + dnEGFR + tuftsin; D) $NOG_{488}$-EGFR complex + aEGFR-longPSE. Images contain the merged channels of i) blue channel - nuclei stained with DAPI; ii) green channel – $NOG_{488}$-EGFR complex iii) Merged fluorescent channels; iv) Merged fluorescent channels + brightfield.

**Table S1. Myeloid panel**

| Antigen | Fluorochrome | Species | Vendor | Clone | Dilution |
|---|---|---|---|---|---|
| CD11b | BUV395 | mouse | BD Horizon (563553) | M1/70 | 1:1600 |
| CD11c | BUV496 | mouse | BD Horizon (750450) | N418 | 1:100 |
| CD49d | BUV563 | mouse | BD Horizon (741243) | 9C10(MFR4.B) | 1:800 |
| Ly6G | BUV661 | mouse | BD Horizon (741587) | 1A8 | 1:500 |
| F4/80 | BUV805 | mouse | Biolegend (123133) | BM8 | 1:400 |
| MHC-II | Pe/Cy7 | mouse | Biolegend (107629) | M5/114.15.2 | 1:4000 |
| Ly6C | BV785 | mouse | Biolegend (128041) | HK1.4 | 1:10000 |
| CD80 | BV605 | mouse | Biolegend (104729) | 16-10A1 | 1:200 |
| CD163 | APC/Cy7 | mouse | Biolegend (155323) | S15049I | 1:400 |
| PDL1 | APC | mouse | Biolegend (124311) | 10F.9G2 | 1:400 |
| CD45 | BUV737 | mouse | BD Horizon (568344) | 30-F11 | 1:4000 |
| Axl | AF488 | mouse | R&D (FAB8541G) | 175128 | 1:200 |
| CD206 | AlexaFluor700 | mouse | Biolegend (141734) | C068C2 | 1:100 |
| P2ry12 | APC-Fire810 | mouse | Biolegend (848013) | S16007D | 1:100 |

**Table S2. Lymphoid panel**

| Antigen | Fluorochrome | Species | Vendor | Clone | Dilution |
|---|---|---|---|---|---|
| CD3 | BV605 | mouse | Biolegend (100351) | 145-2C11 | 1:100 |
| CD4 | BUV496 | mouse | BD Horizon (612952) | GK 1.5 | 1:800 |
| CD8 | Spark Blue 574 | mouse | Biolegend (344783) | SK1 | 1:400 |
| CD69 | BV650 | mouse | Biolegend (104532) | H1.2F3 | 1:400 |
| IFN$\gamma$ | Pe/Cy7 | human/mouse | Biolegend (502527) | 4S.B3 | 1:200 |
| GRZ-B | PE | human/mouse | Biolegend (372207) | QA16A02 | 1:500 |
| CD25 | APC | mouse | Biolegend (102012) | PC61 | 1:400 |
| PD1 | APC/Cy7 | mouse | Biolegend (135224) | 29F.1A12 | 1:1000 |
| Tim-3 | BV711 | mouse | Biolegend (119727) | RMT3-23 | 1:400 |
| LAG-3 | BV421 | mouse | Biolegend (125221) | C9B7W | 1:200 |
| CD45 | BUV737 | mouse | BD Horizon (568344) | 30-F11 | 1:4000 |
| Ki67 | FITC | mouse | Biolegend (652409) | 16A8 | 1:400 |

### Table S3. Distress score sheet

| | | | | | |
|---|---|---|---|---|---|
| Project/Cells injected: | | | | | |
| Date of injection: | | | | | |
| Drug administered at surgery: | Buprenorphin ip, or other | | | | |
| **Cage N°:** | | | | | |
| | **Animal ID (or ear mark)** | | | | |
| | **Weight at surgery** | | | | |
| | Date/Time: | | | | |
| Parameters | Weight: | | | | |
| Drug(s) | Drug's name: route, dose | | | | |
| Appearance | Normal posture and smooth fur<br>Hunched posture OR ruffled fur<br>Hunched posture AND slightly ruffled fur<br>Hunched posture AND all piloerection | 0<br>1<br>2<br>3 | 0<br>1<br>2<br>3 | 0<br>1<br>2<br>3 | 0<br>1<br>2<br>3 |
| Body condition | Normal<br>Loss of 10% body weight<br>Loss of 15% body weight<br>loss of 20% body weight | 0<br>1<br>2<br>3 | 0<br>1<br>2<br>3 | 0<br>1<br>2<br>3 | 0<br>1<br>2<br>3 |
| Clinical signs | Normal respiratory rate and pattern<br>Slight changes, increased rate only<br>Increased rate with abdominal breathing<br>Decreased rate with abdominal breathing | 0<br>1<br>2<br>3 | 0<br>1<br>2<br>3 | 0<br>1<br>2<br>3 | 0<br>1<br>2<br>3 |
| Activity | Normal: moves around the cage<br>Moves slowly around the cage<br>Moves only when touched or seizures (< 3 s)<br>Does not move (paralysis, lethargy) | 0<br>1<br>2<br>3 | 0<br>1<br>2<br>3 | 0<br>1<br>2<br>3 | 0<br>1<br>2<br>3 |
| Provoked behaviour | Normal (moves when cage is disturbed, runs from hand)<br>Attenuated or exaggerated responses (moves away briskly)<br>Moderate change (moves away slowly)<br>Reacts violently, or unresponsive | 0<br>1<br>2<br>3 | 0<br>1<br>2<br>3 | 0<br>1<br>2<br>3 | 0<br>1<br>2<br>3 |
| Score for euthanasia | If 3 is scored in one category, immediate euthanasia | □ | □ | □ | □ |
| Score: 0-15 | 0-4 No action needed | 5-9 Monitor carefully | | day 10-15 Suffering, euthanasia | |

### Table S4. Protein sequences and molecular weights

| Construct | Sequence | MW (Da) |
|---|---|---|
| dnPDL1 | SSGMEEEIECAYDLVEEAEKTGDTRLLKKAYELLEKVAEE ATKSGNPIVIIRLIIILIKIVRNSGDQRVAKLARELLEKLEEH DEKEGNRFVEAMAEALRTQIERALGSHHHHHH | 13119 |
| nbPDL1 | MAQVQLVETGGGLVQPGGSLRLSCTASGFTFSMHAMT WYRQAPGKQRELVAVITSHGDRANYTDSVRGRFTIS RDNTKNMVYLQMNSLKPEDTAVYYCNVPRYDSWGQGT QVTVSSSPSTPPTPSPSTPPCGENLYFQGLEHHHHHH | 16239 |
| dnEGFR | GHHHHHHGSSSDHWEEVFRNALEHLQEATQQNDPQKA KKILCEAHKKLRRELSEEEARSVVRWLKQLVDREKS | 8706 |
